# Spatial control of karyopherin binding avidity within NPC mimics revealed by designer FG- Nucleoporins

**DOI:** 10.1101/2025.04.21.649908

**Authors:** Hendrik W. de Vries, Anders Barth, Alessio Fragasso, Tegan A. Otto, Ashmiani van der Graaf, Eli O. van der Sluis, Erik van der Giessen, Liesbeth M. Veenhoff, Cees Dekker, Patrick R. Onck

## Abstract

Nucleocytoplasmic transport occurs *via* nuclear pore complexes (NPCs), ∼40-60 nm wide pores lined with intrinsically disordered proteins that are rich in Phe-Gly motifs (FG-Nups) that form a selective barrier. Molecules larger than ∼50 kDa are increasingly blocked for transport unless they are bound to a nuclear transport receptor (NTR). How the amino acid sequence of FG-Nups contribute to this is not fully understood. Here, we present *de novo* designed artificial FG-Nups with a systematically varied FG-repeat spacing and charge-to-hydrophobicity ratio (C/H). Starting from a reference sequence termed ‘NupY’ (with the average properties of natural yeast GLFG-Nups), we designed, synthesized, and experimentally tested a library of NupY variants using QCM-D experiments and phase separation assays. We find that the spacing between FG-motifs governs Kap95 absorption into the FG-Nup phase, while increasing C/H results in higher avidity for Kap95 due to an increased accessibility of FG-motifs. Molecular dynamics simulations of transport through NupY-coated pores show a reduced barrier function for noncohesive high-C/H-ratio variants and the highest transport selectivity for designs close to native GLFG-Nups. We postulate that a balance between entropic repulsion and enthalpic gain from multivalent Kap-FG-Nup interactions drives the spatial and temporal partitioning of Kaps in the NPC.

## Introduction

The nuclear pore complex (NPC), a massive protein complex (∼52 MDa in yeast) in the nuclear envelope of eukaryotic cells, regulates the bidirectional transport between the nucleus and the cytoplasm^1,2^. Molecular transport across the NPC is known to be fast and highly selective, but the underlying physical mechanism remains elusive. Intrinsically disordered FG-Nucleoporins (FG-Nups), rich in motifs containing Phe and Gly (*i.e.*, FxFG, GLFG with x any residue), form a selective and dynamic network in the 40 to 60-nm wide central channel^3,4^. This network allows small molecules to pass through while gradually hindering the passage of large macromolecules (masses of ∼20-70 kDa and higher^5,6^) unless these are bound to nuclear transport receptors (NTRs). NTRs are primarily constituted by the karyopherin protein family (Kaps). Interestingly, transport selectivity of Kaps through the NPC is remarkably robust to deletions of FG-Nups^7,8^, dilation or constriction of the central channel^4^, or variations in its structural composition, for example, in the number of symmetric spokes^9^.

The complexity of the NPC and the challenge of probing nucleocytoplasmic transport *in vivo* have inspired many *in vitro* efforts that study assemblies of FG-Nups and their interactions with Kaps, aiming to understand the molecular underpinning of the fast and selective transport. Examples are wide-ranging, both in terms of the employed geometries as well as complexity^10^. As a first example, planar FG-Nup brushes have been shown to selectively interact with different Kaps^11–16^, exhibiting a varying affinity and binding behavior for different pairs of Kaps and FG-Nups^11,17^. Second, condensates and hydrogels formed by native FG-Nups^18–21^ or synthetic constructs^22–25^ showed the selective uptake of Kaps over inert proteins^26^. Lastly, NPC-like selective transport (*i.e.*, allowing transport of Kaps while blocking nonspecific proteins of similar size) has been achieved by coating solid-state nanopores with FG-Nups^27–33^. These *in vitro* efforts have significantly contributed to the understanding of the physical interactions between FG-Nups and Kaps.

Furthermore, synthetic ‘designer FG-Nups’ have successfully recapitulated key aspects of transport and selectivity using simplified sequences of native FG-Nups with homogenized FG-repeat and spacer composition^22,24,25,31^ and variations of the FG-repeat type, number, and density^23,29,34^. The sequences of these synthetic constructs were mostly derived from FG-Nups containing GLFG repeats (GFLG-Nups, such as Nup100) while less attention was given to the role of the spacer regions which contained a low amount of charged amino acids. An important second class of FG-Nups^35^, however, contains FxFG repeats and a high charge content within the spacer regions. Of these, the FxFG Nup Nsp1 was shown to have a modulatory effect on condensates of other FG-Nups such as Nup100 and Nup116^36^ where mutations in the charged residue patterning within the spacer sequences greatly affected its role as phase state modulator. This makes it important to further assess how the physicochemical properties of the spacers affect the self-interaction of FG-Nups, the structure and dynamics of the dense FG-Nup mesh within the central transporter, and the selectivity and efficiency of transport.

Here, we probe the complete parameter space of FG-nucleoporins in a systematic bottom-up study by rationally designing a library of synthetic FG-Nups with varying physicochemical properties. In earlier work^31^, we illustrated that a *de novo* artificially designed FG-Nup, coined ‘NupX’, could form a selective barrier. In the present work, we resolve the essential FG-Nup sequence features that lead to selective transport by designing a much longer 803-residue successor, which we name ‘NupY’. Compared to NupX, the NupY-protein comprises a length and number of FG-motifs that more closely resemble the average values of native GLFG-type Nups from yeast. With this NupY template, we systematically and independently vary two parameters that are key features of FG-Nups, *i.e.*, the spacing between adjacent FG-motifs (d_FG_) and the charged-to-hydrophobic amino acid ratio (C/H). We hypothesize that these two parameters are the key determinants for the selective transport properties of FG-Nup assemblies with Kaps that efficiently traverse the NPC by transiently binding to FG-motifs^34,37,38^. Both C/H and d_FG_ may control Kap diffusivity through and Kap affinity for the FG-mesh, while we expect C/H to be an important driver of the barrier function (the ability to repel inert molecules beyond a soft size barrier of ∼50 kDa) by controlling the density of the FG-Nup network.

Our study presents coarse-grained molecular dynamics (MD) simulations as well as experiments (QCM-D quartz-crystal-microbalance with dissipation monitoring, and phase separation assays) on a large variety of NupY proteins and their interaction with Kap95 (yeast homolog of Importinβ). We find that d_FG_ strongly affects the amount of absorbed Kap95 and Kap95-bound cargo in assemblies of NupY variants, where a d_FG_-value close to the native FG-Nup average yields the highest Kap95 mass flux, while increases in C/H firmly reduce the barrier function and lead to notably enhanced Kap-FG-Nup binding kinetics. The most pronounced transport selectivity appears for d_FG_ and C/H-values close to the native GLFG-Nup average. Notably, we find that Kap avidity to assemblies of FG-Nups strongly depends on the density of the FG-Nup mesh. This implies that entropic repulsion, in addition to posing a permeability barrier to inert probes, also needs to be considered to understand the behavior of Kaps where it can counteract the enthalpic gain of the multivalent Kap-FG interaction. Our insights into the sequence-function relationship of designer FG-Nups suggest that different classes of natively occurring FG-Nups play functionally distinct roles within the NPC, collectively maximizing both transport selectivity (through the cohesive Nups with small FG-spacing) as well as Kap absorption kinetics (through the noncohesive, high-C/H Nups with larger FG-spacings).

### Design of artificial FG-Nucleoporins with controlled sequence properties

To disentangle the sequence-function relationship for the formation of the selective transport barrier, we constructed a library of artificial FG-Nucleoporins by systematically varying two parameters: 1) the FG-spacing (d_FG_), namely the number of amino acids between consecutive FG-repeats, which sets the total number of FG-motifs within the FG-mesh, and 2) the ratio of charged-to-hydrophobic amino acid content (C/H) of the full sequence (Figure 1a). Together, we expected that the FG-spacing strongly controls the multivalency of the network towards Kap95^34^, whereas the C/H would most strongly affect the cohesivity of the FG-Nups, thereby modulating the passive permeability barrier formed by the FG-Nup mesh^5,29,39^.

**Figure 1:**
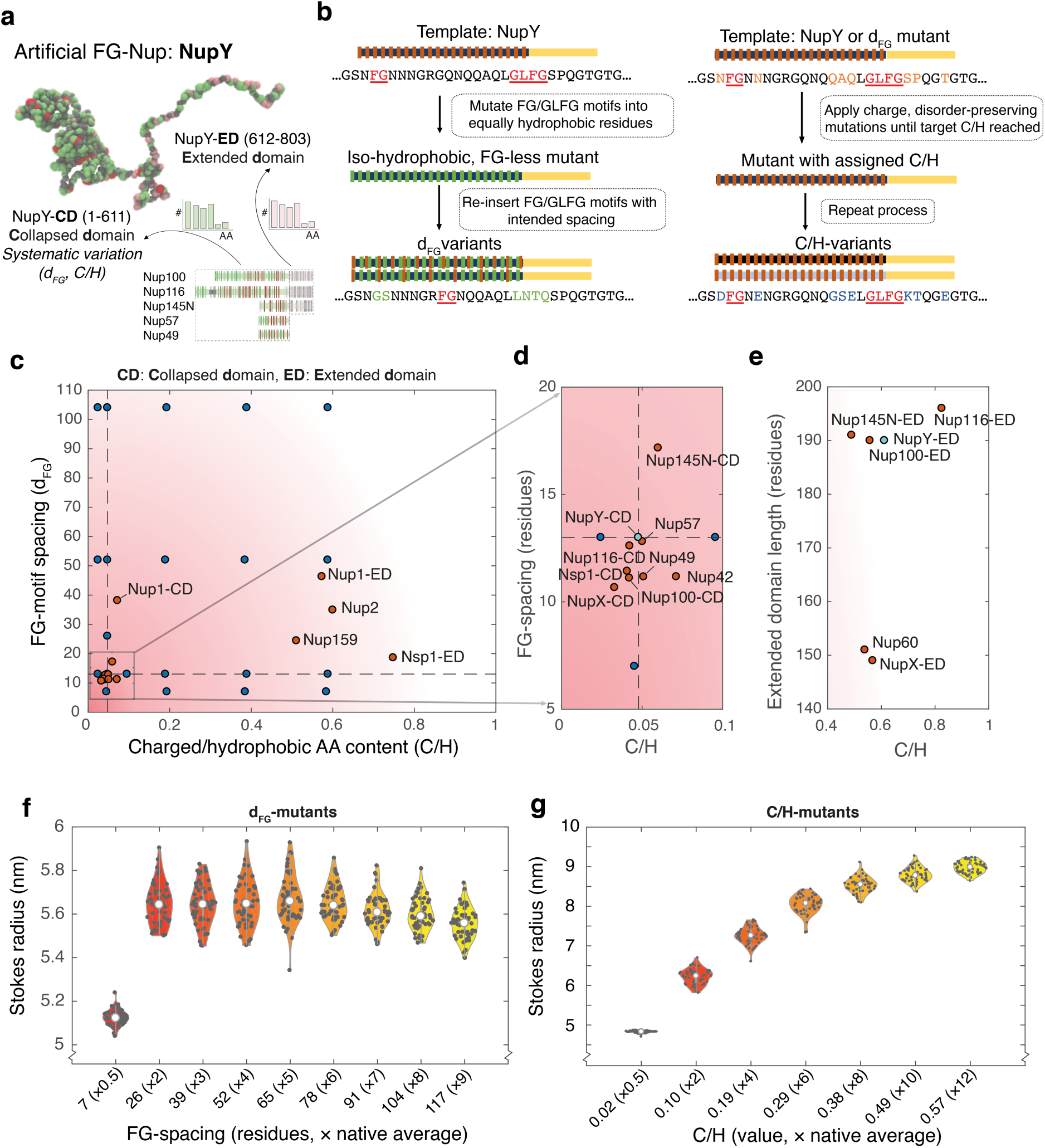
Study approach and design of a collection of artificial FG-Nups a: Design of the artificial FG-Nup ‘NupY’, with a two-domain structure: a collapsed domain (CD) and extended domain (ED). We mutated the CD (residues 1:610) throughout this study via a systematic variation of the FG-motif spacing and the composition of the spacers (total C/H). F-residues are shown in red, polar uncharged residues (N, Q, T, S) in green, and charged residues (D, E, K, R) in pink. The picture represents a simulation snapshot. **b:** Schematic overview of the generation of artificial FG-Nup sequences with designated FG-motif spacings and C/H-values, by performing various mutation steps on the NupY-sequence. The yellow region represents the extended domain, devoid of FG-motifs. **c:** Comparison between the C/H-values and FG-motif spacings in native *saccharomyces cerevisiae* FG-Nup domains (dark orange) and our NupY variants (blue). Our designs span a range of C/H-values and FG-motif spacings encompassing virtually all native FG-domains. A dashed line indicates the average C/H and d_FG_ for native GLFG-Nup domains, which underlie the NupY template. Definitions of collapsed domain (CD) and extended domains (ED) were adopted from Yamada *et al.*^35^ (Supplementary Table 2), where we note that we adhered to a different definition of C/H (methods). **d:** Close-up of cohesive native yeast FG-Nups or collapsed domains (in case of bimodal Nups). Included are also an artificial FG-Nup ‘NupX’ designed earlier^31^, and NupY(13;0.05), which forms the basis for all mutations in this study (cyan). **e**: Close-up of select FG-Nup extended domains (EDs) that fell out of panel (c). Instead of FG-spacing, we display the entire domain length since these domains do not comprise FG-motifs. **f:** Violin graph showing the calculated Stokes radii (Methods), *R_S_*, for 50 designs per FG-motif spacing, each with the GLFG-Nup average C/H of 0.05. Each dark scatter point indicates the average *R_S_* of one design. The *R_S_*-distribution for variants with a specific FG-spacing is narrow, indicating that the polymer properties of these designs are similar. The effect of FG-motif spacing on *R_S_* is minor for variants with d_FG_>13 (native average) since the mutations preserve the C/H and patterning of hydrophobic residues. Our designs for d_FG_=7 did not fully preserve the patterning of hydrophobic residues, leading to additional compaction. **g:** Violin graph showing the calculated *R_S_* for 50 different designs per C/H, each with the GLFG-Nup average d_FG_ (13 residues). The polymer properties are similar for designs with the same C/H; increasing the C/H leads to a monotonic increase in *R_S_*. The scaling of the *R_S_*-values with C/H is similar for different d_FG_-values (Supplementary Figure 2).

Our design procedure took place in two distinct stages. First, we adopted the design rules applied for the artificial FG-Nup ‘NupX’ in earlier work^31^ and designed a full length 803-residue counterpart that we term ‘NupY’. Both proteins derive from a class of FG-Nups rich in GLFG-motifs that are deemed essential for cell viability^7^. Briefly stated, the design method randomly assigns residues to an extended and collapsed domain following the statistical distribution of amino acids in such domains in yeast GLFG-Nups (for a full description, see Methods). Whereas the NupX-protein considered the ‘average’ sequence properties of GLFG-Nups in terms of FG-motif spacing and overall sequence composition, it was notably shorter (with only 311 residues) than most native GLFG-Nups and did not consider the ratio between FG and GLFG-motifs. The NupY-protein (Figure 1a) resembles an average GLFG-Nup (Methods) more closely: it comprises a 610-residue collapsed domain that contains FG and GLFG-motifs in a 4:3 ratio, and a 190-residue extended domain that is devoid of FG-motifs. The final amino acid sequences included a Cys-residue (for end-grafting to surfaces) at the C-terminus, and a Gly-Pro-motif at the N-terminus, which remained after cleaving off a tag used in purification (Methods), putting the total sequence length at 803 residues. To produce NupY-variants, we varied only the collapsed domain that contains the FG-motifs, while we kept the extended domain without FG-motifs (611-800) the same for all the variants. To distinguish between the different variants, we label each variant by its spacer length and C/H-value, *e.g.,* ‘NupY(13;0.05)’ for the NupY variant with d_FG_=13 and C/H=0.05.

Starting from the ‘average GLFG-Nup’ NupY(13;0.05), we designed variants with a varying d_FG_ by performing mutations that replaced FG and GLFG-motifs with isohydrophobic groups of residues such as AA, MQ, IS, VT and reinserting FG or GLFG-motifs at designated positions (Methods). In these variants (Figure 1b-c, Supplementary Figure 1a), we maintained a constant C/H (namely at the GLFG-Nup average of 0.05) while varying d_FG_ between 7 and 117 residues, covering a range from approx. 0.5 to 9 times the GLFG-Nup average. Interestingly, coarse-grained molecular dynamics simulations (Methods) of 50 design variants for each value of d_FG_ (Figure 1f) illustrate that the Stokes radius (*R_S_*) remains highly similar between variants with different d_FG_ values but with similar C/H. We postulate this to be a consequence of the conserved C/H and spacing of groups of hydrophobic residues (Methods), preserving the collapsed state of the 1-610 domain despite reducing the number of FG-motifs. NupY(7;0.05) displayed a reduced *R_S_*-value due to the increased density and high degree of patterning of hydrophobic residues, despite maintaining a constant C/H by the insertion of more hydrophilic residues in the spacer regions (see Methods).

Using the set of d_FG_-variants as inputs, we then generated designs with varying C/H, achieved by iteratively replacing spacer residues for disorder-promoting^35^, non-aromatic residues (including charged and polar residues) while preserving the net charge of the protein (Figure 1b-c, Supplementary Figure 1b, Methods). For example, if an increase in C/H was desired and a hydrophobic I-residue was picked, a replacement was chosen from the remaining pool of less cohesive, disorder-promoting amino acids (A, G, Q, S, P, M, T, H, E, K, D). The pool of available residues depended on the hydrophobicity of the picked residue and the direction of the C/H mutation. We designed NupY-variants with C/H ranging from 0.02 (0.5x the GLFG-Nup average) up to 0.57 (12x), a range large enough to also include sequences similar to the extended domains of FG-Nups (Figure 1c). We did not further study designs for NupY(7-0.02) due to insufficient predicted disorder (Methods). Coarse-grained modeling of C/H variants at constant d_FG_=13 illustrated minor variations in *R_S_*between design variants with identical target C/H, whereas the *R_S_*-value monotonically increased with increasing C/H due to the decreasing cohesivity (Figure 1g, Supplementary Figure 2). In the following, we present results for one sequence for each combination of C/H and d_FG_ for further studies (Supplementary Table 1).

The complete set of designs spans almost the entire physiological range of native *Saccharomyces cerevisiae* FG-Nups in terms of d_FG_ and C/H (Figure 1c-e) and provides us with the opportunity to characterize the effect of the two parameters on the functionality and selectivity of the NupY protein. In the following, we performed experimental studies on polymer brushes using QCM-D and condensates formed via phase separation (PS) and evaluate the functional consequences for transport in a nanopore geometry using coarse-grained modeling.

### FG-spacing and C/H impact Kap95 selectivity and adsorption kinetics in FG-Nup brushes

To assess the effect of FG-spacing and C/H variations on the polymer properties of NupY and the affinity to Kap95, we performed QCM-D experiments on a subset of seven different NupY variants that includes the template protein NupY(13;0.05), the FG-spacing variants NupY(52;0.05) and NupY(104;0.05), the C/H variants NupY(13;0.09), NupY(13;0.19) and NupY(13;0.38), as well as a high C/H variant with large FG-spacing, NupY(104;0.57) (Supplementary Table 1). FG-Nup proteins were first grafted to a clean gold-coated quartz sensor (Figure 2a), followed by titration of Kap95 at increasing concentrations (Figure 2d, Methods)^31^.

**Figure 2:**
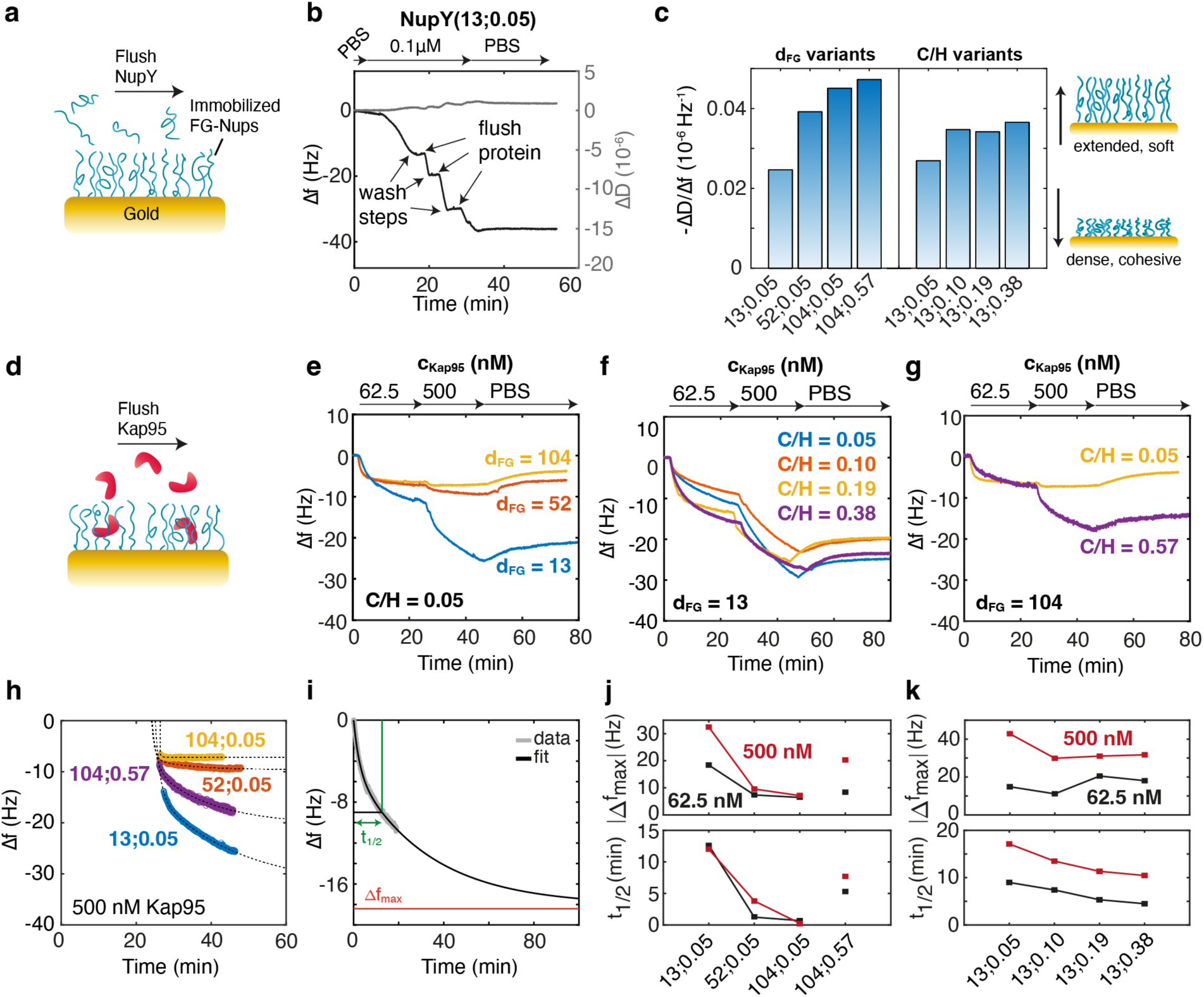
QCM-D characterization of NupY variants. a: NupY proteins self-assembled onto the gold chip surface through thiol-gold chemistry to form a dense protein brush (Methods). **b:** During the coating of the gold surfaces, changes of the frequency (Δf, black) and dissipation (ΔD, gray) report on the binding of FG-Nups. Shown is the variant NupY_13–0.05_. All NupY variants showed qualitatively similar Δf and ΔD, indicating that similar grafting densities could be achieved (Supplementary Figure 15). **c:** Average dissipation-to-frequency ratio (-ΔD/Δf) of NupY brushes. We observed an increase in - ΔD/Δf both as a function of FG-spacing and C/H ratio, indicating reduced stiffness. **d:** Binding of Kap95 to pre-formed FG-Nup brushes. (**e-g**) Kap95 is titrated to pre-formed NupY brushes at 62.5 nM and 500 nM, followed by a buffer wash. Larger FG-spacing reduces Kap95 binding **I** while variations of the C/H ratio shows no systematic change **(f)**. In the case of NupY(104;0.57) (**g**) the high C/H ratio appears to rescue the Kap95 binding to the brush. **h)** Bi-exponential fits of the Δf curves for Kap95 binding at 500 nM to the indicated NupY variants, extracted from the curves shown in **e-f**. Fits to all binding curves are shown in Supplementary Figure 15. **i)** Example of fitting and extraction of the parameters Δf_max_ and t_1/2_ from the Δf curve for Kap95 binding to NupY(13;0.05) at 62.5 nM. **j-k)** Extracted parameters |Δf_max_| (top) and t_1/2_ (bottom) for the different variants at Kap95 concentrations of 62.5 nM (black) and 500 nM (red) for FG-spacing variants **(j)** and C/H variants **(k)**. All experiments were performed once (N=1).

For a direct comparison between the NupY variants, we aimed, to the extent possible, to obtain comparable grafting densities by reaching a similar frequency shift of-40 Hz during the NupY coating step. This required different protein concentrations ranging from 0.1 μM for NupY(13;0.05) up to 1 μM for NupY(104;0.05) and NupY(104;0.57) (Supplementary Figure 3a-h). We generally found that the higher the d_FG_ or C/H ratio, the higher the protein concentration needed to reach a certain frequency shift. This trend is expected and consistent with previous work^16^ that reported a positive correlation between surface coverage and the protein’s propensity to self-interact, which in the present case is mainly a result of inter-and intra-chain hydrophobic interactions. In support of this, we also observe that the dissipation-to-frequency ratio (-ΔD/Δf, Figure 2c), a measure of brush softness and level of hydration (Methods)^16,40^, increased both with d_FG_ and C/H, suggesting that the NupY brushes are becoming more extended.

After an additional passivation step to minimize nonspecific interactions between proteins and the gold surface^11,13,17,31,41^ (Supplementary Figure 4b), we sequentially titrated Kap95 (the main protein importer in yeast) at concentrations of 62.5 and 500 nM, which resulted in different frequency responses for the tested NupY variants (Figure 2e-g). As observed in earlier work^13,31^, dissociation of Kap95 from the FG-Nup brush was slow when washed in PBS buffer (see for t>40min), but Kap95 was reversibly released upon flushing NaOH (Supplementary Figure 4c). When adding BSA as an inert control protein, we observed a complete lack of interaction for all NupY variants (Supplementary Figure 5), indicating that all NupY variants show clear selective behavior, qualitatively similar to native FG-Nups. While Kap95 molecules bound well to our template NupY(13;0.05), with frequency shifts comparable to those observed for native FG-Nups in previous studies^15,31,42^, binding to the FG-spacing variants NupY(52;0.05) and NupY(104;0.05) was severely reduced as evident from the reduced frequency shift (Figure 2e). On the other hand, frequency shifts for the C/H variants were similar as for the template (Figure 2f, top), consistent with the constant number of FG-repeats. To our surprise, the combination of high C/H ratio and large FG spacing in the variant NupY(104;0.57) significantly enhanced Kap95 binding compared to NupY(104;0.05), up to levels comparable to the original template NupY(13;0.05s) (*cf.* Figure 2e and g). In contrast, for an FG spacing of 13 the increase in C/H did not further boost the adsorption of Kap95 (Figure 2f).

To further quantify these trends, we fitted the frequency response at the different Kap95 concentrations to a bi-exponential function (Figure 2h-i, Methods, Supplementary Figure 6, Supplementary Table 6), which allowed us to determine two parameters: Δf_max_, which corresponds to the maximal Δf reachable for 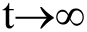indicative of the maximal Kap95 occupancy at a given concentration, and t_1/2_, which is the time to reach Δf_max_/2 and which quantifies how fast binding saturation occurs (see Figure 2i). In agreement with surface-plasmon resonance experiments^41^, the deviation of the frequency response from single-exponential behavior suggests the existence of a spectrum of affinities as the NupY brushes saturate with Kap95. For the FG spacing variants (Figure 2j), Δf_max_ decreased with increasing FG-spacing, consistent with a lower amount of Kap95 binding to the FG-Nup brush due to a depletion of available FG-repeats. Interestingly, also t_1/2_ decreased with d_FG_ due to a faster binding saturation for variants with less FG repeats. For the C/H variants (Figure 2k), no clear trend for Δf_max_ was evident, with similar saturation values for all tested C/H-values. However, t_1/2_ steadily decreased with increasing C/H variants (Figure 2k, bottom). We attribute the latter effect to the more extended and dynamic brushes formed by the higher C/H variants, reasonably causing FG-motifs to be more accessible to Kap95 and causing it to be incorporated faster into the protein brushes. C/H thus is an important parameter for tuning the binding rate of Kap95 to the FG-Nup brushes by increasing the availability of FG repeats, while having no effect on the total amount of bound Kap95.

Notably, for the variant with the largest FG-spacing of 104, we observed an increased Δf_max_ and t_1/2_ at high C/H (NupY(104;0.57)) compared to low C/H (NupY(104;0.05), see Figure 2j). On the contrary, for an FG-spacing of 13, an increase of C/H only resulted in faster binding, but not increased uptake of Kap95 (Figure 2k). This apparent difference suggests that Kap95 readily partitions into the dense brushes of NupY(13;0.05) due to the high FG-motif density (which does not increase further at higher C/H), whereas the low FG-motif density for the variant NupY(104;0.05) is not sufficient to overcome the steric repulsion experienced by Kap95 in the cohesive FG-Nup mesh. The latter is reduced in the high C/H variant NupY(104;0.57), which rescues Kap95 binding to a level comparable to NupY(13;0.05) but with slower binding saturation.

In summary, the characterization of NupY brushes reveals that, while FG-spacing is an important parameter for tuning the binding strength towards Kap95, C/H also plays a crucial role as it can promote efficient Kap95 binding due to enhanced accessibility of FG-motifs, even for very sparse FG-patterning.

### FG-spacing and charge to hydrophobicity ratio determine Kap95 and cargo uptake into condensates

We studied the interaction of Kap95 with condensates that were formed via PS by rapid dilution of NupY protein into denaturant-free buffer. We reached final concentrations of FG-Nups (after dilution) of 200 nM. Different to the FG-domain of the native Nup100, neither the template NupY(13;0.05) nor any of the other NupY variants formed detectable amounts of condensates at this concentration as assessed by a sedimentation assay and brightfield microscopy (Figure 3a-b). An exception was the variant NupY(7;0.05), which, likely due to the high amount of cohesive FG motifs, formed irregularly shaped particles that were more resistant to disruption of hydrophobic interactions by the aliphatic alcohol 1,6-hexanediol (Supplementary Figure 7c-d). This observation is in line with the reduced Stokes radius seen in the MD simulations (Figure 1f).

**Figure 3:**
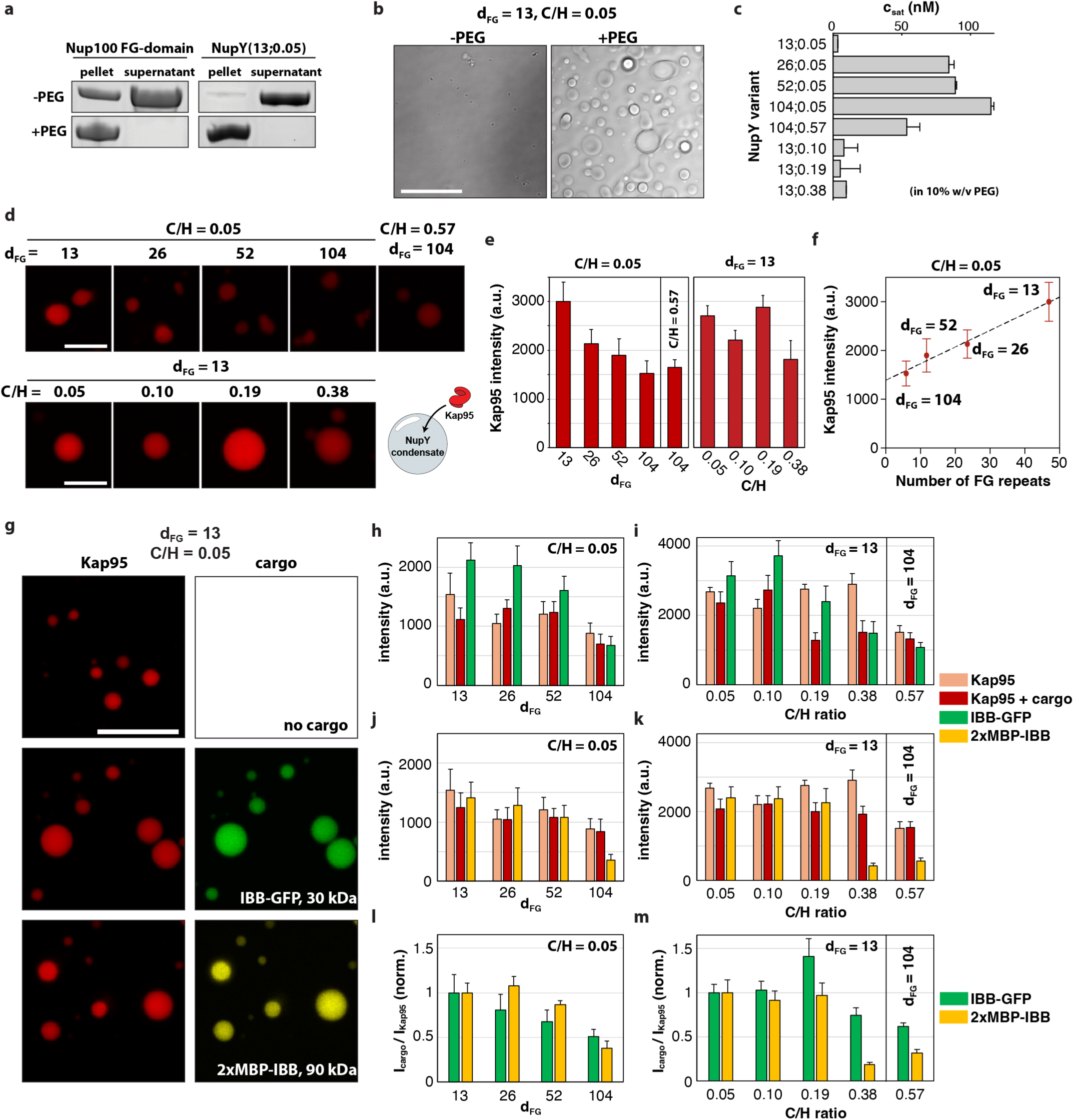
Phase separation of NupY variants. a: Sedimentation assay to probe phase separation of FG-Nups. The FG-domain of Nup100 (a.a. 1-610) and NupY(13;0.05) were diluted to a concentration of 200 nM in buffer containing 150 mM NaCl with or without 10% w/v PEG-8000 and condensates were pelleted by centrifugation. **b**: Brightfield images of NupY(13;0.05) condensates formed at 1 µM in the absence or presence of 10% w/v PEG-8000. Scalebar: 20 µm. **c:** Concentration of the dilute phase c_sat_ for the different NupY variants in the presence of 10% w/v PEG-8000. **d:** NupY condensates were formed at 200 nM and challenged with 1 µM Kap95-Alexa647. Scalebar: 5 µm. **e:** Kap95 intensity in NupY condensates for the different variants. **f:** Linear fit of the Kap95 intensity as a function of the number of FG-repeats. **g:** NupY condensates were challenged with 1 µM Kap95-Alexa647 without cargo and in presence of 1 µM IBB-GFP or 1 µM 2xMBP-IBB labeled with Alexa488. Scalebar: 10 µm. **h-j**: Intensities of Kap95 alone, Kap95 in the presence of cargo, and cargo for the different NupY variants. Note that cargo intensities are not comparable between IBB-GFP and 2xMBP-IBB due to differing brightness of the fluorescent label. **l-m:** Ratio of the cargo intensity to the Kap95 intensity. To facilitate a comparison, the intensity ratios are normalized to the intensity ratio obtained for the template NupY(13;0.05) and hence represent the relative change of the cargo uptake per transporter compared to the template.

Addition of a macromolecular crowder (PEG-8000 at 10% w/v) sufficiently lowered the saturation concentration c_sat_, i.e., the concentration above which the system phase separates, such that condensates formed for all variants (Figure 3a-c, Supplementary Figure 7a-b). Control experiments using fluorescently labeled PEG-8000 did not show significant partitioning of the crowder into the condensates (Supplementary Figure 8). We observed differences in the c_sat_ in the presence of molecular crowding (Figure 3c). While the template NupY(13;0.05) showed negligible solubility under the experimental conditions due to the high self-interaction propensity, c_sat_ increased significantly for larger FG spacing, in line with the higher concentrations required in the QCM-D experiments to reach equal surface coverage (Supplementary Figure 3a-h). This indicates that FG-FG interactions are the main factor for self-interaction of FG-Nups at low C/H. At native FG-spacing (d_FG_ = 13), C/H had no clear effect on the PS propensity and all C/H variants showed near-complete condensation with negligible protein amount in the dilute phase. On the other hand, for d_FG_ = 104 an increase of C/H from 0.05 to 0.57 resulted in a lower c_sat_ and larger PS propensity, indicating that electrostatic interactions between charged residues or cation-π interactions between arginines and the phenylalanines of the FG-repeats promote self-interaction in the dense phase. Condensates formed by the NupY variants, with the exception of NupY(7;0.05) (Supplementary Figure 7c) showed a spherical morphology. However, merging of droplets generally remained incomplete, likely owing to the high viscosity of the condensates and their strong adhesion to the cover glass (Figure 3b).

We then challenged condensates of the different NupY variants with 1 µM fluorescently labeled Kap95 (Figure 3d). We detected negligible signal of Kap95 in solution as the protein efficiently partitioned into the condensates but did not yet reach saturation (Supplementary Figure 7e). Increasing the FG-spacing gradually lowered the density of Kap95 within the condensates while variation of the C/H ratio did not reveal a clear effect on Kap95 density (Figure 3e). While the high C/H variant with d_FG_ = 104 rescued the binding of Kap95 in the QCM-D experiments, we did not observe significant differences in the uptake of Kap95 into the condensates between the variants NupY(104;0.05) and NupY(104;0.57) in the LLPS assay. Kap95 intensity in the condensates depended linearly on the number of FG repeats and hence FG-motif density (Figure 3f). However, the non-zero offset indicates that non-FG-binding site interactions of Kap95 with the condensates also contribute under the experimental conditions.

We next assessed the Kap95-mediated uptake of two model cargoes that were fused to the importin-beta binding (IBB) domain, namely (i) GFP (IBB-GFP, 30 kDa) and (ii) a concatemer of the maltose-binding protein labeled with the organic dye AlexaFluor 488 (2xMBP-IBB, 90 kDa). Both cargoes showed negligible interactions with the condensates in the absence of Kap95 (Supplementary Figure 9) but partitioned efficiently into the condensates of all tested NupY variants when Kap95 was present (Figure 3g), showing that NupY proteins can form a selective phase akin to native FG-Nups^21,26,43^. The amount of cargo in the condensates generally followed the partitioning of Kap95 and decreased with increasing FG-spacing (Figure 3h,j). Upon comparing the Kap95 signal in the absence and presence of cargo, we did not observe any significant difference for variations of d_FG_ at a constant C/H of 0.05 (Figure 3h,j), but uptake of Kap95 in the presence of cargo was hindered at high C/H ratio for d_FG_ = 13 (Figure 3i,k). This suggests that a high C/H ratio poses a larger barrier for partitioning of the bulky Kap95-cargo complex. To directly compare the uptake of the two cargoes, we computed the intensity ratio of cargo to Kap95 and normalized all values to the intensity ratio obtained for the template NupY(13;0.05) (Figure 3l,m). The resulting quantity corresponds to the cargo uptake per Kap95 protein, normalized to the value obtained for the template NupY(13;0.05). For IBB-GFP, the cargo uptake per Kap95 molecule steadily decreased with increasing FG-spacing, as expected from the reduced enthalpic gain of Kap95-FG interaction due to the lower FG-motif concentration. On the other hand, the larger cargo 2xMBP-IBB still showed similar cargo uptake for d_FG_ = 26 and 52 compared to the template (Figure 3l) despite lower FG-motif concentration for these variants. This suggests that variants with larger FG-spacing could more frequently form wider openings within the network that can accommodate the large cargo and hence reduce entropic repulsion. The C/H variants showed a small reduction in uptake for the small cargo but a drastic reduction for the large cargo at a C/H ratio of 0.38 (Figure 3m). The lowered uptake of the large cargo at high C/H is also seen for the variants NupY(104;0.05) and NupY(104;0.57), which otherwise behaved similarly. This suggests that high C/H ratio results in a denser network that is harder for the bulky Kap95-cargo complex to penetrate.

To gain insights into the mobility of Kap95 within the condensates, we performed fluorescence recovery after photobleaching (FRAP) experiments (Figure 4a-b). For all tested NupY variants, recovery was found to be slow and incomplete and did not depend on FG spacing or C/H, with a mobile fraction of only 10-20% and recovery times of several minutes. This indicates a very low mobility of Kap95 within NupY condensates that are populated by transport receptors. To assess the kinetics of Kap95 uptake into empty NupY condensates, we added 1 µM of Kap95 immediately before imaging. Contrary to the FRAP experiments, influx of Kap95 into unchallenged condensates was fast and complete on the timescale of ∼30 s (Figure 4c-d). These results suggest a strong hindrance of the mobility of Kap95 on the mesoscopic scales probed in the FRAP experiments. To confirm this result, we performed a competition experiment with differently labeled Kap95 by first challenging condensates of NupY(13;0.05) with Kap95-Alexa488 (cyan), followed by incubation with Kap95-Alexa647 (Figure 4e). As expected, Kap95-Alexa488 was initially quickly taken up by the condensates, however Kap95-Alexa647 only bound to the surface and was unable to partition into the pre-challenged condensates even over the timescales of hours.

**Figure 4:**
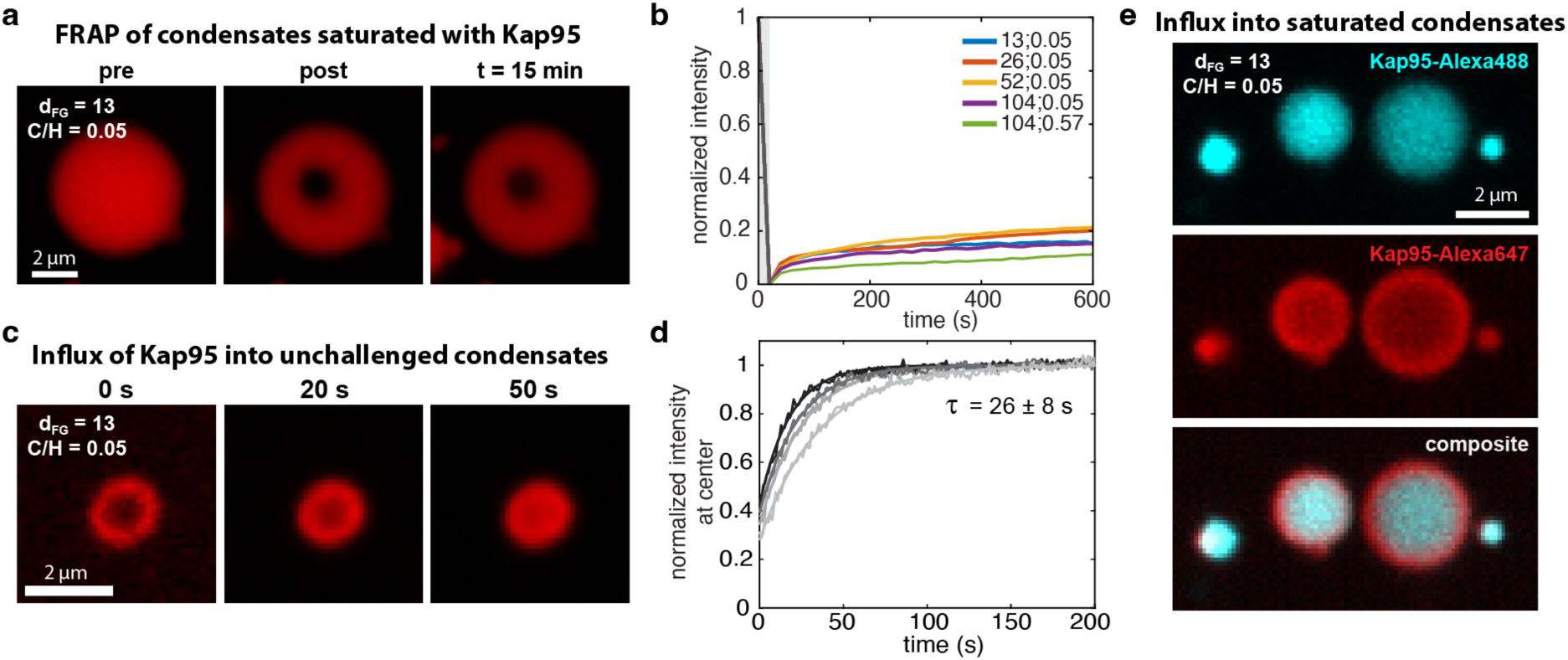
Mobility of Kap95 in NupY condensates. **a)** Fluorescence recovery after photobleaching experiment on a condensate of NupY(13;0.05) challenged with Kap95 at a concentration of 1 µM. **b)** FRAP curves of Kap95 obtained for Kap95-challenged condensates of different NupY variants show incomplete recovery. See Supplementary Figure 10 for details. **c)** Influx of Kap95 into a condensate of NupY(13;0.05). **d)** Kap95 influx into condensates of NupY(13;0.05) is monitored by the intensity at the center of the condensate, normalized to the maximum intensity value at the end of the experiment. Individual curves belong to different condensates. **e)** Condensates of NupY(13;0.05) were first challenged with Kap95-Alexa488 at 1 µM (top) and subsequently exposed to Kap95-Alexa647 at 1 µM (middle). A composite image is shown below.

In summary, these data revealed the FG-motif density as the main driving force of phase separation and showed that Kap95 efficiently partitioned into condensates formed by NupY in an FG-repeat-density-dependent manner, despite potential steric hindrance due to high FG-Nup density in the condensates. Cargo uptake was likewise efficient, however high C/H-values increasingly hindered the uptake of bulky Kap95-cargo complexes. Despite fast influx of transporters, condensates showed low mobility of Kap95 after having taken up sizeable amounts of transporters, most likely due to their high affinity to the FG-Nup phase, resulting in low off-rates, or mutual hindrance of diffusion between Kap95 molecules.

### Simulations show that FG-Nup cohesivity controls the dynamic network morphology of NPC mimics

To assess the functional consequences of variations of d_FG_ and C/H on transport through the NPC, we used residue-scale coarse-grained molecular dynamics simulations to study the internal structure of nanopores coated with the set of designed NupY variants (Figure 5a, Supplementary Figure 11). We selected a nanopore diameter of 55 nm, comparable to the *in situ* diameter of the transport channel in the NPC^3^, and a grafting distance between adjacent NupY anchor points of 5.5 nm, in line with our earlier work on FG-Nup coated nanopores (Materials and Methods)^29,31^. We first describe the FG-Nup distribution inside pores coated with the template protein NupY(13;0.05) (second panel in Figure 5b-e). The extended anchoring domains (residues 612-803), combined with a grafting density comparable to the Stokes radius (Figure 1f), caused the FG-rich collapsed domain (1-611) to localize away from the pore wall. This resulted in low protein densities near the rim of the pore (comprising the extended anchoring domain). This general feature appeared in all nanopore simulations in this work since pore diameter, grafting density and the composition of the extended anchoring domain remained unchanged. The collapsed FG-rich domain, on the other hand, formed a dense region towards the pore center, with a hyperboloid shape that protruded slightly out of the pore membrane (see Figs. 4c and e).

**Figure 5:**
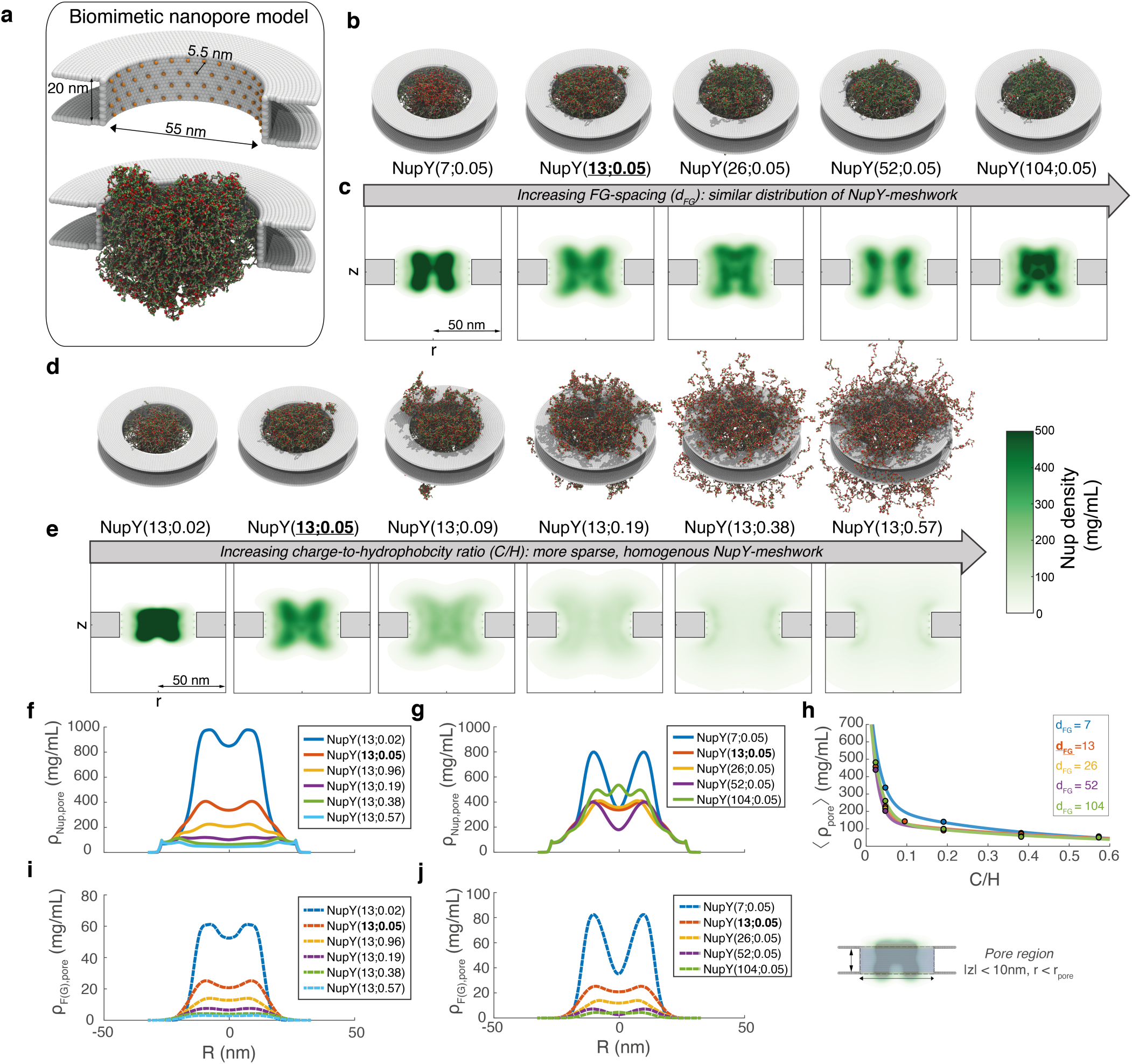
Effect of C/H and FG-spacing on the internal organization of FG-Nup coated nanopores. a: Snapshot of the computational model for FG-Nup coated nanopores. Red spheres highlight the FG-motifs. **b-c:** Snapshots (top) and axi-radial, time-averaged density graphs (bottom) of NupY-coated nanopores with native GLFG-Nup average C/H (0.05) and varying d_FG_. The extended NupY-anchoring domains caused the dense, FG-rich domains to accumulate in a hyperboloid structure centrally within the pore, rather than towards the pore wall. In line with the single-chain simulations (Figure 1f-g), increasing d_FG_ from the GLFG-Nup average did not strongly affect the structure of the FG-Nup network, whereas a decrease in FG-spacing enhanced the density of the FG-Nup network. **d-e:** Snapshots (top) and axi-radial, time-averaged density graphs (bottom) of NupY-coated nanopores with native GLFG-Nup average d_FG_ and varying C/H. Changes in C/H strongly affected the localization of the FG-Nup domains. Starting from the ring-like structure formed by the collapsed, FG-rich domains of the NupY-proteins, a decrease in C/H lead to the formation of a dense, central plug that localized almost entirely within the pore membrane. Increasing C/H decreased the interaction strength of the FG-containing domains, leading to a an increasingly sparse and homogenous distribution of protein mass that covers a large region outside of the pore interior as well. **f,g:** Time-averaged radial FG-Nup density profile within the pore region (defined as the cylindrical volume within the pore membrane) under varying C/H (f) and d_FG_ (g). **h**: Protein density, averaged over the pore region, as a function of C/H, for NupY-variants with combined variations in C/H and d_FG_. A bi-exponential fit is provided as a guide to the eye. **i,j:** Time-averaged radial F(G)-motif density profile within the pore region under varying C/H (i) and d_FG_ (j). Importantly, the density of FG-motifs scaled with both C/H and d_FG_, where the shape of the distribution correlated strongly with that of the total Nup density, due to the presence of FG-motifs in the collapsed domain.

At a constant C/H of 0.05 (GLFG-Nup average), the average density and spatial distribution of the protein network in the nanopore interior were relatively unaffected by step-wise increases in d_FG_ from 13 up to 104 (Figure 5b-c,g). This finding is consistent with the low variability of the Stokes radius (Figure 1f, Supplementary Figure 2) between d_FG_-variants. The smallest value of d_FG_=7 provided an exception; the compaction due to an increased density of (evenly spaced) hydrophobic motifs along the chain similarly lead to an increased protein density in the central annular structure (Figure 5b-c first panel, Supplementary Figure 11). Overall, the density distribution of FG-motifs decreased with increasing d_FG_ (Figure 5j, Supplementary Figure 13).

Next, we considered the effect of variations of C/H at a constant d_FG_ of 13 residues. Changes in C/H caused a remarkable change in the morphology of the FG-Nup network over the range from 0.02 (0.5x native average) to 0.57 (12x, Figure 5d-f). Compared to the protein distribution for NupY(13;0.05), a decrease in C/H caused the FG-rich domain to form a dense central plug with densities approaching 1000 mg/mL, rather than a hyperboloid structure, featuring a density distribution that almost entirely localized inside the pore (Figure 5d-e, first panel). On the other hand, increases in C/H beyond the GLFG-Nup average caused the dense central annular structure to dissolve (Figure 5e-f). This transition took place when increasing C/H from 0.09 to 0.19. The least cohesive variants (C/H=0.38 and 0.57, rightmost panels in Figure 5d-f) displayed a homogenous distribution of FG-Nup mass with the Nups sampling a large volume of space outside of the pore interior. The distribution of FG-motifs (Figure 5i, Supplementary Figure 13) as a function of C/H largely coincided with that of the FG-Nup mass (Figure 5f).

Finally, we assessed whether the effects of d_FG_ and C/H on the distribution of NupY-variants was similar across combined variations of d_FG_ and C/H. The average densities inside the pore interior as a function of C/H displayed bi-exponential behavior: The pore average density (Figure 5h) showed a bi-exponentially decreasing trend with increasing C/H. The average protein density was insensitive to d_FG_, indicating that the properties of the protein networks are determined by the C/H value, as supported by the similar axi-radial density graphs (Supplementary Figures 11-12). We note that C/H variants under a constant d_FG_ of 7 residues were always more compacted, consistent with our findings on isolated NupY-variants (Figure 1f) and d_FG_-scaling (Figure 5b-c, g). The spatial distribution of FG-motifs under combined variations of C/H and d_FG_ largely followed that of the total FG-Nup density (Supplementary Figure 13), where we note that some differences exist for noncohesive variants.

The protein distributions in NupY-coated pores thus most strongly depended on C/H, with a less pronounced role for d_FG_. While the changes in NupY network morphology with varying C/H are reminiscent of earlier modeling results, we note that the exact organization of FG-Nups inside a nanopore geometry depends critically on the nanopore dimensions and grafting density^31,44^, Nup length and cohesivity^39,45,46^, and the domain structure of the FG-Nups^47^. For example, protein networks in earlier work displayed central plugs (e.g., Nsp1^29,32,48^, NupX^31^), ring-like structures (NupX^31^, yeast NPC^49–53^, Nup98^29^), or sparse and homogenous structures (Nsp1_FILV→S_^29^) depending on the anchoring pattern, protein length, protein cohesivity or pore diameter.

#### FG-repeat density and FG-Nup cohesivity govern the distribution of proteins in NupY-coated pores

To assess the transport selectivity of nanopores coated with NupY-variants, we performed simulations in the presence of the transport receptor Kap95 and inert probes of different sizes. In our residue-scale coarse-grained models (Methods), we explicitly considered interactions of three groups of amino acid residues in folded proteins and the NupY variants, namely charged residues^54^, aromatic residues^21^, and FG-specific binding sites^55–57^, that all affect a protein’s ability to interact with FG-Nups.

First, we performed single-molecule binding simulations between single copies of NupY-variants and Kap95 (Figure 6a-b) to obtain the apparent dissociation constant *K_D_* (under the assumption of binary complex formation, see Methods) between Kap95 and all NupY-variants. Our simulations showed a strong dependence of the binding affinity between Kap95 and NupY on the number of FG-motifs and the C/H value (Figure 6b). Increasing the number of FG-motifs by reduction of the FG-spacing led to more frequent interactions between FG-motifs and the binding sites on the surface of Kap95 (and thus a lower *K_D_*-value), consistent with nuclear magnetic resonance (NMR) and isothermal titration calorimetry (ITC) measurements on FG-Nup segments with varying number of FG-motifs^34^. Interestingly, an increase of C/H resulted in increased affinity, especially for variants with fewer FG-motifs (large d_FG_). In agreement with the larger Stokes radius for higher C/H (Figure 1g), the larger extension of the noncohesive NupY-variants increased the availability of FG-motifs for interactions with Kap95 compared to more cohesive variants. This suggests a combined role of d_FG_ and C/H in governing NupY-Kap95 binding, where the binding strength increases both with the number of FG-motifs and the accessibility of such motifs (C/H). Interestingly, this effect seems to manifest itself in the QCM-D measurements (Figure 2g), where the absorption of Kap95 greatly increased when the C/H-value of a large spacing variant NupY(104;0.05) was increased from 0.05 to 0.57.

**Figure 6:**
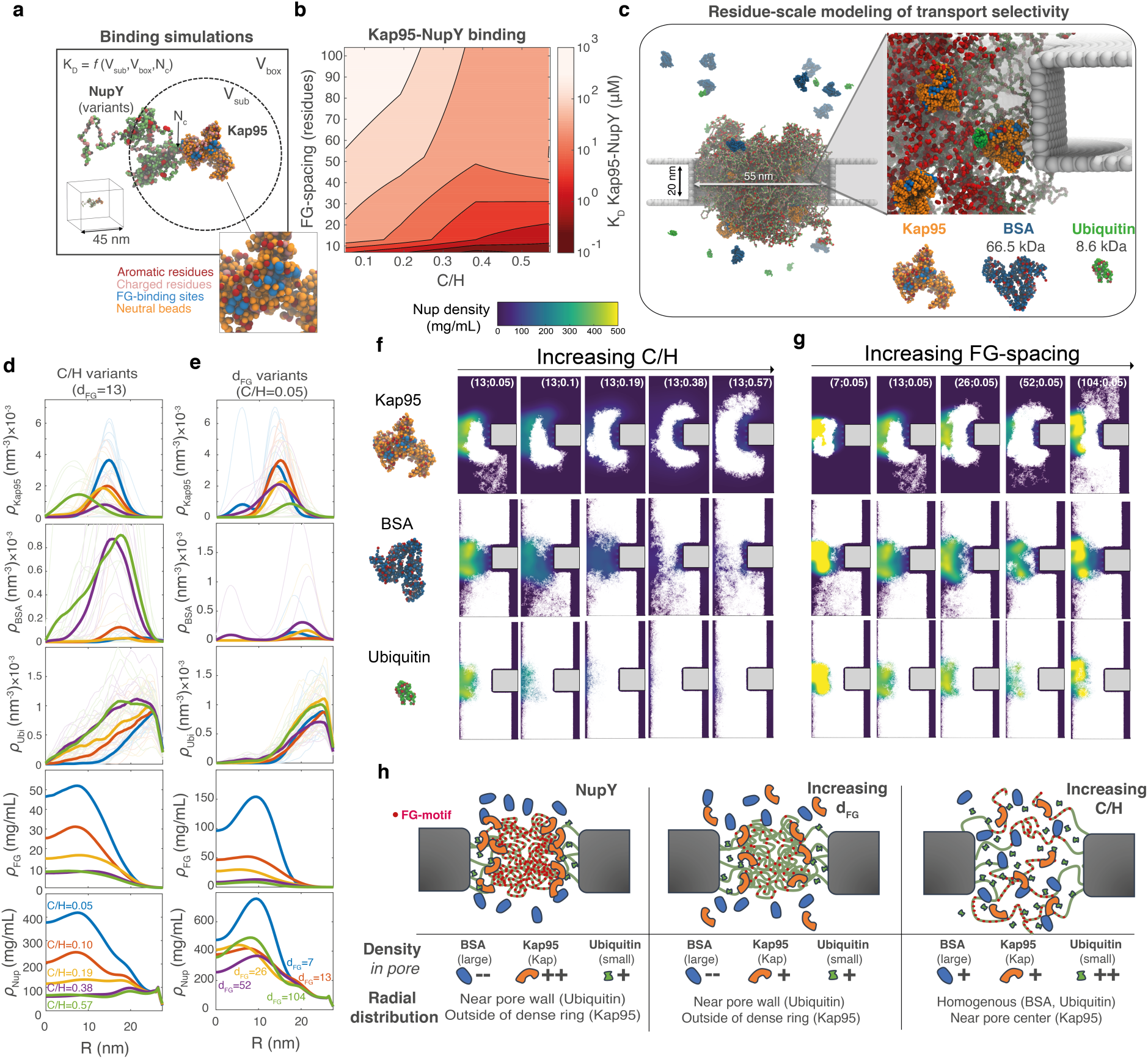
Localization of FG-Nups, Kaps, and inert proteins in nanopores lined with artificial FG-Nups. a: We obtained the dissociation constant (*K_D_*) between Kap95 and NupY variants using single-molecule binding simulations. *K_D_* was calculated from the number of binding and non-binding contacts within a protein pair. **b:** *K_D_*(log-scale) as a function of d_FG_ and C/H. The *K_D_*-value decreased with decreasing d_FG_ due to the increased number of FG-motifs that can participate in Kap95 binding. Increases in C/H enhanced binding to Kap95 due to an enhanced exposedness of the FG-motifs, an effect that is most pronounced for weak binders with high d_FG_. **c:** 1-BPA computational model of transport selectivity. *Left:* Nanopore coated with NupY-variants in the presence of the folded proteins. Our transport simulations comprised the yeast Kap Kap95 and inert proteins smaller (ubiquitin) and larger (BSA) than the soft size cut-off for passive transport. We released ten copies of each cargo type into the FG-Nup network. *Right:* Zoomed-in view of binding site-FG-motif interactions that allow Kap95 particles to localize inside the pore network. For clarity, FG-Nups are transparent, with FG-motifs highlighted in red. **d:** Effect of varying C/H on the time-averaged radial number densities profiles of inert proteins or Kap95 (averaged over all copies), and the radial mass densities of FG-motifs and FG-Nups within the pore region (|z|<10 nm). Lighter shadings indicate the number density profiles of individual folded proteins. Increases in C/H lead to a sparser and homogenous FG-Nup network (bottom panels). For Kap95 (top panel), the flattened density profiles with increasing C/H reflect this trend, whereas, for the least cohesive NupY-variant, Kap95 molecules localized in the pore center due to the low Nup density. The in-pore densities of BSA (second panel) and ubiquitin (middle panel) increased with C/H, where both molecules localized in the sparsest regions away from the pore center. **e:** As in d, but for varying d_FG_. Kap95 (top panel) localized at the boundary of the dense, ring-like structure formed by the FG-Nups (see bottom two panels), where the average density decreased with increasing d_FG_. No apparent effect of d_FG_ on the localization or density of inert molecules is visible: in all cases, any permeation by BSA (second panel) or ubiquitin (middle panel) occurred in the peripheral channel formed by the extended NupY-anchoring domains. Local trapping of inert molecules explains the outliers in the density profiles for individual molecules. **f:** Localization of the three molecules (Kap95, BSA, ubiquitin) inside NupY-variants with native GLFG-Nup average d_FG_ (13) and varying C/H. Increasing C/H caused Kap95 to sample a larger volume, whereas inert molecules increasingly localized inside the FG-Nup network. **g:** As in (f), but for native GLFG average C/H (0.05) and varying d_FG_. Depending on the local FG-motif density (controlled by d_FG_), Kap95 either sampled a small volume of the pore interior (d_FG_=7), localized near the dense lobes (d_FG_=13 to 52), or partially outside of the pore (d_FG_=104). The localization of ubiquitin and BSA was largely unaffected. **h:** Schematic overview of cargo localization for different NupY-variants, where ‘-‘ and ‘—’ indicate hindrance or blockage, and ‘+’ or ‘++’ reflect the ease of a molecule to permeate the pore. For pores with near-native d_FG_ and C/H, Kaps localized near (but did not necessarily permeate) the dense central structure formed by collapsed FG-domains. Increases in d_FG_ and C/H reduced the concentration of Kaps inside the pore lumen: either by reducing the ability of Kaps to associate with the dense ring-like structure (d_FG_ increase) or by spreading out more homogenously (C/H increase). We found that variation in d_FG_ did not directly affect the localization for nonspecific molecules: small molecules permeated mainly via the sparse part of the FG-Nup network near the pore wall, whereas large molecules were hindered. For increasing C/H, more inert molecules localized within the pore lumen.

Next, we performed transport simulations where ten copies each of three types of proteins were released into NupY-variant-coated pores simultaneously (Figure 6c, Methods). The three proteins studied in our nanopore simulations were Kap95 (94.7 kDa), BSA as a large inert protein (66.5 kDa), and ubiquitin as a small inert protein (8.6 kDa), which cover the mass range (∼20-70 kDa) associated with the onset of the NPC’s transport barrier (Figure 6c, Methods). We first assessed the effect of C/H variation on cargo localization by inspecting the radial density profiles and trajectories of the three probes (Figure 6d,f). In the template NupY(13;0.05)-coated pores, Kap95 avoided the regions with the highest FG-motif density (Figure 6f, top left and Figure 6d, top panel) and localized at the outer boundary of the dense, hyperboloid-shaped regions. This suggests that the binding interaction between Kap95 and FG-motifs was counteracted by steric hindrance due to high local protein density. While BSA was largely excluded from the densely-filled pore (Figure 6d,f, middle row), ubiquitin localized preferentially near the outer rim of the pore where FG-Nup density was lower due to the extended anchoring domains. Increases in C/H at a constant d_FG_ of 13 caused a reduced steric hindrance for all three probes due to the lowered FG-Nup and FG-motif density within the pore. At higher C/H, the distribution of Kap95 more closely followed the density of FG-motifs. The increasingly sparse and extended distribution of FG-Nups and FG-motifs at high C/H caused Kap95 to sample a larger volume of the FG-Nup network including regions outside of the pore membrane, while still being excluded from the pore center (Figure 6d,f top). For the least cohesive case NupY(13;0.57), however, the distribution of Kap95 shifts towards the pore center due to the lowered FG-Nup density and reduced cohesion (Figure 6d, fifth panel), despite the lowered FG-motif density throughout the pore (Figure 6d, fourth panel). The localization of inert cargo (BSA, ubiquitin) similarly varied with increasing C/H: the average density of BSA and ubiquitin increased notably beyond a C/H-value of 0.19, where both molecules localized more towards the pore center as the concentrated regions with FG-Nups disappeared.

The distributions of the three probes for d_FG_-variants at native C/H (0.05) showed similar behavior to pores comprising the baseline NupY(13;0.05) design since the spatial FG-Nup arrangement of the d_FG_-variants was similar (Figure 5b-c). For all variants, Kap95 did not permeate the dense central structure formed by the collapsed NupY-domains, and instead localized at its outer boundary where the Nup density is not too high and the FG-motif density not too low (Figure 6d, top panel, Figure 6g, top row). Increasing d_FG_ reduced the uptake of Kap95 and caused it to preferentially localize farther away from the dense central region due to the increasing difficulty of FG-Kap95 binding (due to the reduced FG-motif density) to overcome the steric hindrance of the FG-Nup network (which is only moderately affected by d_FG_). In accordance with the consistent spatial FG-Nup distribution, the radial density profiles were similar between all d_FG_-variants for both BSA and ubiquitin. BSA molecules were largely excluded in all cases (Figure 6e,g, second row), whereas similar amounts of ubiquitin localized near the pore rim for all variants (Figure 6e,g, third row). Ubiquitin displayed a higher number density than BSA due to its smaller size (Figure 6b).

The localization of the three cargo molecules under combined variations in C/H and d_FG_ (Supplementary Figures 16-18) displayed trends similar to those displayed in Figure 6d-g. Of interest is the combined effect of C/H and d_FG_ on the localization of Kap95: in noncohesive variants with large FG-motif spacings (NupY(52;0.57) and NupY(104;0.57)), Kap95 localized homogenously throughout the FG-Nup mesh, which was not seen for d_FG_ or C/H variants with the native GLFG-Nup average as fixed parameter. This can be traced back to the high availability of the FG-motifs in the dilute network at high d_FG_ and C/H (Figure 6b), featuring dissociation constants that are similar to the template NupY.

We summarize the effect of C/H and d_FG_ variations in Figure 6h; spatial variations in FG-Nup and FG-motif density led to spatial variation in the localization of the three studied cargoes. Inert molecules preferentially localized towards sparse regions with low FG-Nup density, which existed for all NupY variants near the pore wall due to the common extended anchoring domain, and throughout the pore for noncohesive variants (C/H of 0.19 and higher). Kap95 localized near but not inside the dense, FG-rich central structure for cohesive variants due to a competition between steric hindrance (Nup density) and binding with FG-motifs. For increasing C/H, Kap95 spread more homogenously throughout the FG-Nup network and closely followed the density of FG-motifs.

### Native GLFG-Nups provide optimal transport selectivity

We next assessed the role of d_FG_ and C/H in transport selectivity by calculating translocation rates for the inert probes, defined as the total number of full traversals in any direction across the pore, per microsecond of simulation time, as well as the mass flux for all types of molecules. The translocation rates of ubiquitin showed a gradual increase with increasing C/H, with non-zero translocation rates for all NupY variants (Figure 7a). In addition, there was a small but significant increase of the ubiquitin translocation rate with increasing d_FG_ as there was less Kap95 in the peripheral channel for these variants (Figure 6e). The translocation rates of BSA showed a similar trend and increased with C/H while no clear trend was evident with respect to the value of d_FG_ (Figure 7b). Notably, the translocation rates for BSA showed a step-wise behavior and increased significantly beyond a C/H value of 0.19, while the rates remained near-zero at a C/H of 0.19 and below. Pores comprising NupY-variants with C/H-values above 0.19 showed a loss of barrier function as the translocation rates and permeation of BSA into the pore network increased notably for these systems (Figure 7b, Figure 6d,f). The translocation rates of the inert probes BSA and ubiquitin can be understood intuitively as a function of the average FG-Nup protein density within the pore (Figure 7c). For both proteins, transport rates decreased significantly when the average FG-Nup density in the pore region was in a range between 75 and 100 mg/mL. Above this value, the transport rates plateaued at zero for BSA (blue curve, top panel) and a finite value for ubiquitin (orange curve, bottom panel) due to permeation via the sparse regions at the pore wall, formed by the extended anchoring domains.

**Figure 7:**
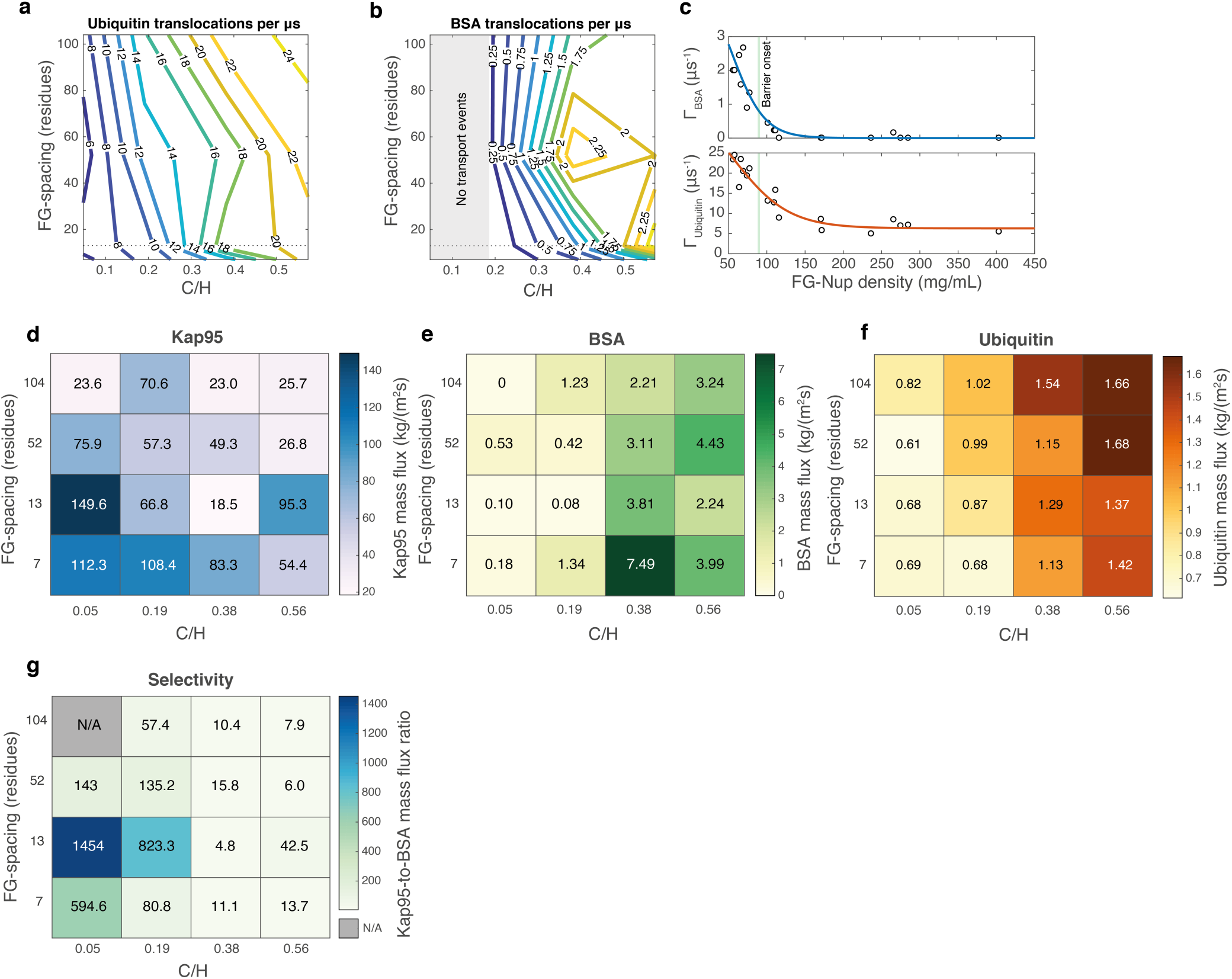
Quantifying transport selectivity in nanopores with FG-Nup variants. a: Contour plot of translocation events per microsecond of simulation time for BSA as a function of C/H and d_FG_. The data indicate that a leakage (non-zero translocations on average) starts to occur for C/H larger than 4x the native GLFG-Nup average (C/H=0.19), where further increases of the C/H (to 12x) lead every studied pore to leak BSA. The change in slope for contour lines at low d_FG_ is a consequence of the increased Nup density for NupY-variants with d_FG_ = 7. **b:** Translocation events per microsecond simulation time for ubiquitin as a function of C/H and d_FG_. Between the native average (C/H=0.05) and the 12xC/H-variant (C/H=0.57), the translocation rate increases up to fourfold. The slight diagonal orientation of the contour lines indicates that d_FG_ affects the permeability of ubiquitin.**c:** Permeability ρ (translocation events per microsecond of simulation time of inert proteins) as a function of FG-Nup density within the nanopore. BSA permeability (top panel) rapidly dropped between 50 and 100 mg/mL average density, being essentially blocked in pores with an average FG-Nup density beyond ∼120 mg/mL. Ubiquitin (bottom panel) translocations still occurred at high average protein densities due to sparse regions (extended NupY anchoring domains) near the pore wall (see Figure 6), explaining the non-zero plateau. **d:** Mass flux of Kap95 as a function of d_FG_ and C/H. A global maximum is clearly visible for the native GLFG-Nup average NupY(13;0.05). **e-f:** Mass fluxes of the inert proteins BSA and ubiquitin as a function of d_FG_ and C/H. The mass flux of BSA and ubiquitin increased notably with C/H. The barrier function of NupY-variant pores strongly decreased between C/H-values of 0.19 and 0.38. A local optimum exists for BSA (NupY(7;0.38)), due to a slightly attractive interaction between BSA and NupY(7;0.38) driven by cation-pi interactions **g:** Transport selectivity, defined as the ratio of the Kap95 and BSA mass flux, as a function of d_FG_ and C/H. We found an optimal transport selectivity for NupY(13;0.05), which corresponds to the native GLFG-Nup average of d_FG_ and C/H.

For Kap95, the direct calculation of translocation rates was not possible as Kap95 molecules remained in the pore interior for the entire duration of the simulation due to their high affinity to the FG-Nup mesh. To quantify the transport behavior of Kap95 molecule, we calculated the mass flux across the pore from the spatial distributions of the protein’s density and its velocity (see Methods). This analysis was also performed for the BSA and ubiquitin molecules in parallel to the translocation rate analysis. The mass flux of Kap95 decreased with either increasing C/H or increasing d_FG_, (Figure 7d). These trends can be understood as follows: with increasing C/H or d_FG_, the average magnitude of the Kap95 velocity along the pore axis increased, an effect counteracted by the decreased Kap95 concentration, yielding an overall decrease in mass flux (see Supplementary Figure 20 for individual trends in Kap95 occupancy and velocity).

Importantly, the highest Kap95 mass flux was obtained for the template NupY(13;0.05), corresponding to the native GLFG-Nup average (Figure 7d). The cohesive variant NupY(7;0.05) with the smallest FG-spacing resulted in a lower mass flux than NupY(13;0.05) due to the reduced mobility of Kap95 molecules (Supplementary Figure 20b), which localized on the boundary of the dense central structure where the local FG-motif density was high. Unexpected local maxima in Kap95 mass flux occurred for NupY(104;0.19) and (NupY(13;0.56). Compared to variants with the same C/H but smaller spacings, the Nup distribution of NupY(104;0.19) shows a more strongly localized pattern, leaving a well-defined peripheral channel for the Kap95 molecules. For the NupY(13;0.56) variant, the NupY density was the lowest (Figure 6d) and Kap95 accumulated in the pore center. For BSA and ubiquitin, the trends in the mass flux under varying C/H and d_FG_ (Figure 7e-f) showed generally similar behavior as the translocation rates in Figure 7a-b yet provide a more nuanced view on selectivity. Since abortive transport events still contributed to the mass flux, we found that even in absence of BSA translocations, a non-zero (yet very low) BSA was present. The BSA mass fluxes in the ‘leaky’ pores NupY(7;0.38) and NupY(13;0.56) formed outliers. In the case of NupY(7;0.38), a slightly attractive electrostatic (cation-pi driven) interaction existed between this NupY-variant (which carried relatively more cationic residues than variants with larger spacings) and BSA (Supplementary Figure 17), which led larger amounts of BSA to associate with the FG-Nup network. For NupY(13;0.56) the accumulation of Kap95 in the center of the pore caused reduced the Finally, we defined a selectivity score as the ratio of the mass fluxes of Kap95 and BSA (Figure 7g). The optimal transport selectivity occurred for NupY(13;0.05), which corresponds to the average properties of GLFG-Nups, consistent with the idea that this may have been evolutionary optimized. The scaling of selectivity with C/H and d_FG_ reconciles the trends we identified for Kap95 mass flux (Figure 7d) and BSA permeability (Figure 7b,e). For d_FG_=13 and higher, the selectivity decreased with larger d_FG_ owing to the reduced affinity of the FG-Nup mesh to Kap95, resulting in a reduced mass flux of Kap95 across the pore (Figure 7d). For d_FG_=7 at low C/H, the selectivity slightly decreased as Kap95 molecules effectively get stuck due to the high FG-motif density. Selectivity decreased strongly with higher C/H, mainly due to a higher leakage of BSA through the less cohesive FG-Nup mesh (Figure 7b,e).

In summary, our transport modeling highlighted that the barrier function is predominantly determined by C/H, while FG-spacing had a minor effect. Translocations of the small probe ubiquitin was present throughout and increased further with reduced protein density in the pore, while the larger probe BSA showed a stepwise behavior with full blockage at small C/H and significant leakage beyond a C/H of 0.19. The mass flux of Kap95 was determined by its affinity to the FG-Nup mesh, which decreased with both the FG-repeat density (d_FG_) and C/H. Transport selectivity was optimal for d_FG_ and C/H-values corresponding to the native GLFG-Nup averages, which combined a high mass flux of Kap95 with a low leakage of BSA (d_FG_=13, C/H=0.05, Figure 7g).

## Discussion

In this work, we rationally designed artificial FG-Nups (‘NupY’) to identify the role of FG-spacing (d_FG_) and cohesivity (C/H) on selective nuclear transport. Following the sequence of essential GLFG-Nups, NupY has a bimodal structure consisting of a 190-residue high charge (extended) domain with fixed sequence and no FG-repeats and a 610-residue designer (collapsed) domain in which we independently varied d_FG_ and C/H while controlling other sequence properties like number of aromatic residues and net charge. Our design differed from previous studies in two aspects^25^. First, we considered a significantly longer FG-domain, which allowed us to probe a broad range of FG-repeat spacings. Second, we performed systematic variations of the C/H ratio. The resulting library of NupY proteins enabled us to cover the full physiological range of C/H-values and d_FG_-values of native FG-Nups (Figure 1c). We first evaluated various polymer properties of the different NupY variants. An increase of C/H both increased the Stokes radius in simulations (Figure 1f-g) and the softness/extension of polymer brushes in QCM-D experiments (Figure 2c), while variations of the FG spacing showed a minimal effect in simulations but a strong softening of the polymer brushes in QCM-D. Condensation experiments showed an increase of the saturation concentration with FG-spacing (Figure 3c), consistent with results on a similar designer FG-Nup^25^ and an NMR study displaying the role of FG-motifs in self-interactions^58^. Self-interaction/cohesiveness thus depends mainly on the density of FG-motifs in the FG-Nup brush or condensate, while C/H seems to predominantly affect the extension of the polymer.

Interestingly, the phase separation propensity showed little dependence on C/H at low FG spacing. Similar effects were seen in prion-like domains, where c_sat_ remained constant upon variation of the overall charge content as long as the net charge was unaltered^59^. Interestingly, the phase separation propensity was enhanced at large FG-spacing for the high C/H variant NupY(104;0.57) compared to NupY(104;0.05). We hypothesize that in this specific case, the more extended configuration of the NupY(104;0.57) variant increased the availability of FG-motifs, thus enhancing their contribution to phase separation, driving c_sat_ down. While the macromolecular crowder did not partition into condensates of NupY (Supplementary Figure 8), it remains unclear whether the increased self-interaction of NupY in the presence of crowding remains purely entropy-driven^60^ or whether crowding could also increase the protein density within the dense phase^61^ and promote a liquid-to-gel transition^62,63^.

The QCM-D experiments showed that the FG-spacing controls the binding of Kap95 to the NupY brushes with the expected trend of decreased binding with increasing FG-spacing (Figure 2 and 3), confirming that FG-repeats are the primary driver of the Kap-Nup interaction. Interestingly, we also found that increasing C/H in a weakly absorbing brush for the variant NupY(104;0.05) rescued the ability to bind Kap95 efficiently by increasing FG-motif accessibility by increased C/H in NupY(104;0.57). On the other hand, increasing the C/H of the spacers in a strongly absorbing NupY(13;0.05) brush results in similar saturation levels of bound Kap95 but notably faster adsorption kinetics (Figure 2). The experimental data thus indicate that FG-spacing and C/H critically tune the binding strength and adsorption rate of Kap95. Kap95 binding is qualitatively similar between the QCM-D measurements and k_D_ simulations: decreasing d_FG_ enhanced binding, while increasing C/H can lead to more FG-motifs being exposed and available for binding with Kap95 (Figure 6a-b), while reducing the energetic penalty of displacing FG-Nup mass, resulting in faster absorption (Figure 2k). This highlights the importance of FG-motif availability for Kap95 binding and underscores that local protein density and FG-motif concentration must be considered to understand transport selectivity.

The condensation assay also showed a clear reduction of Kap95 partitioning as the FG-spacing is increased (Figure 3e-f). Different from the QCM-D experiments, however, we observed significant Kap95 partitioning even at large FG spacing (Figure 3e). This highlights key differences in the experimental design of these assays. In the QCM-D experiments, the low density of FG-motifs in the polymer brushes for NupY(104;0.05) is not sufficient to overcome the entropic repulsion experienced by Kap95. However, considering high reported FG-Nup densities within condensates (up to 300-600 mg/ml for a similar system^25^), the density of FG-motifs seems to remain high enough for efficient partitioning into the condensates even at large FG-spacing. Our data also suggest a residual affinity of Kap95 even in the absence of FG-motifs (Figure 3f), which either indicates nonspecific interactions of Kap95 with other amino acid sidechains of NupY or a nonlinear behavior of Kap95 at low FG-repeat densities. As in the QCM-D experiments, we obtained similar Kap95 partitioning at a physiological FG-spacing of 13 regardless of C/H (Figure 3e). However, condensates showed similar Kap95 uptake for the variants NupY(104;0.05) and NupY(104;0.57). This suggests that the concept of FG-motif availability in polymer brushes or nanopore geometries does not apply at the high protein concentrations within the condensed phase. As entropic repulsion within the dense condensates is expected to be high for all NupY variants regardless of C/H, the affinity of Kap95 is primarily determined by the binding enthalpy and hence the FG-motif density.

The uptake of two model cargoes of different size (30 and 90 kDa) provided insights into the mesh size and density of the FG-Nup network. Cargo uptake showed a similar dependence on the FG-spacing as Kap95 alone (Figure 3h,j,l), however the larger cargo complex showed a less steep decrease with increasing FG-spacing with similar uptake at d_FG_ = 13 and 26 (Figure 3l). Together with the observation that FG-FG interactions are defining for self-interaction, we hypothesize that the FG-motif spacing could dictate the size of openings/voids in the dense condensate as well as how frequently they appear and reduce the entropic repulsion experienced by bulky cargo complexes. C/H had a less pronounced effect on cargo uptake, however at high C/H-values of 0.38 and above we observed a notably reduced cargo uptake.Lastly, FRAP experiments showed a strong hindrance of the diffusion of Kap95 within condensates, consistent with the idea of a resident population of Kap95 in the NPC^64–66^. In accordance with the coarse-grained modeling, efficient transport thus most likely occurs in regions where the FG-Nup network is less dense and cohesive and Kap95 molecules are more mobile.

Coarse-grained modeling of nanopores coated with the template NupY(13;0.05) revealed that the collapsed, FG-rich domains formed a hyperboloid-shaped density distribution within the 55-nm pore, while the extended FG-free anchoring domains cause low protein densities near the outer rim (Figure 5b-c). Whereas the distribution of FG-Nup mass showed modest changes when varying d_FG_, changes to C/H had a notable impact on the distribution of FG-Nups and FG-motifs (Figure 5d-e). As the value of C/H increased, the FG-Nup density in the central regions gradually diminished and the FG-Nup mass redistributed to regions outside of the pore (C/H of 0.19 and above, Figure 5f,h). Simulations of the multivalent interaction between individual molecules of the NupY-variants and Kap95 showed the expected scaling of the binding affinity with the number of FG-repeats. Increasing C/H promoted the extension of FG-Nup chain and enabled FG-motifs to become more easily available for the binding sites on Kap95, causing improved binding (lower *K_D_*, Figure 6a-b). Repeating the modeling of NupY variant-coated pores in the presence of the inert probes ubiquitin and BSA, as well as Kap95 uncovered how the changing FG-Nup and FG-motif distributions due to varying d_FG_ and C/H controlled protein translocation. FG-spacing strongly affected the localization and mass flux of Kap95, with Kap95 showing a tendency to populate the parts of the FG-Nup mesh with the highest FG-motif density yet sufficiently low steric hindrance. For cohesive NupY-variants, these regions predominantly occurred at the outer boundary of the dense region. C/H affected the localization (Figure 6d-g) and mass flux (Figure 7a-c) of both Kap95 and inert proteins. Interestingly, an optimum in Kap95 mass flux and transport selectivity occurred for NupY(13;0.05), which corresponded to the native average GLFG-Nup (Figure 7d). We also found that in pores with noncohesive FG-Nups comprising low d_FG_, Kap95 localized in the pore center and sequestered FG-Nups towards the pore interior (see Figure 6d,f, Supplementary Figure 16). Barrier functionality (blockage of BSA) strongly decreased beyond a C/H of 0.19 (Figures 5d-f, 6b,e), consistent with the value at which the dense region disappeared.

Throughout this study, we observed major effects of d_FG_ and C/H on the microenvironment that Kap95 and inert molecules experience, where the particular nature of this microenvironment differed (e.g., geometric constraints and anchoring density of the FG-Nups) between the various approaches used in this study (QCM-D, condensates, and nanopore simulations). A view that emerges from our findings is that large local variations in the FG-Nup and FG-motif density effectively cause large local variations in avidity between Kap95 and the FG-Nup network. Note that the avidity in this context corresponds to an effective binding strength (*i.e.*, a free energy) experienced locally by a Kap inside the FG-Nup network. It hence combines the counteracting effects of enthalpic attraction due to the favorable multivalent Kap-FG interaction and entropic penalties via steric repulsion due to the overall protein density of FG-Nups and Kap95 in the pore. A large Kap avidity thus corresponds to regions with a high concentration of binding sites combined with low density-induced steric repulsion. Local variations in Kap avidity, together with the steric repulsive effect of the FG-Nup mesh density on BSA and ubiquitin localization (barrier function), explain most of the key observations on selectivity in this study. For example, at the non-saturated Kap95 concentrations used in our modeling, Kap95 preferentially localized in regions where the FG-motif density was locally high, but at the same time the FG-Nup density sufficiently low (i.e., Figure 6d-h). This manifested in Kap95 localizing outside of the dense lobes in cohesive variants, yet in the pore center of noncohesive NupY-variants. We observed a similar effect in our recent work on Nsp1^32^, where yeast Kap95 localized in regions comprising extended, FG-repeat containing domains (unique to Nsp1 and Nup1), rather than inside dense regions (formed by Nsp1’s collapsed domains) with higher local FG-motif densities. In our QCM-D experiments, the local balance between steric hindrance and the local density of FG-motifs explains how NupY-variant brushes absorb less Kap95 with increasing d_FG_, have similar saturation amounts of bound Kap95 (but faster on-rates) for increasing C/H, and why increasing C/H for the weak-binding NupY(104;0.05) variant rescued Kap95 binding. In our condensation experiments, the counteracting effects of steric repulsion and binding enthalpy is evident from the observation that Kap95 uptake was strongly dependent on the FG-motif density (given that the overall FG-Nup density is expected to be similar between the different NupY variants^25^), and that uptake of large Kap95-cargo complexes was reduced due to increased steric repulsion.

The insights from all data, which cover the sequence space of native FG-Nup domains in yeast, allow us to describe an operating mechanism for the NPC central channel in the presence of Kaps. This model is based on the arrangement of the two categories of FG-Nups within the NPC, as determined from recent structure determination^3,4,67,68^ and simulation studies^52,69–71^ in combination with a spatially varying Kap avidity. The noncohesive FxFG-type Nups, located at the entrance and exit of the pore and along the central axis, sequester Kaps with a maximized on-rate (Figure 6b and 2j-k), and enable Kaps to enter the central pore region. The cohesive GLFG-Nups, forming a high-density region positioned near the inner ring^71^ provides the NPC with a steric barrier against non-NLS/NES cargo. At the same time, Kaps experience the highest binding avidity towards the FG-Nup network within the pore’s central channel, just outside the high-density GLFG-rich region. Recent studies underscore such a mechanism and suggested behavior of NTRs that can be explained by considering spatial variations of NTR avidity^65,71–74^. Specifically, one study found crowding-induced spatial segregation of NTRs within FG-Nup assemblies, where small NTRs such as NTF2 localized inside denser regions of these assemblies^72^. We postulate that smaller NTRs, which have a lower number of FG-binding sites and may compete with larger NTRs such as Kap95 or CRM1 in lower-density regions of the NPC^75^, would indeed be able to permeate further into the dense lobes by virtue of their smaller size, leading to avidity-driven spatial segregation of NTRs of different size in the NPC. Moreover, a recent experimental study^65^ showed that a significant Kap95 population is present centrally in the NPC’s FG-Nup network (populated by the largely non-cohesive Nsp1^71^), which was essential for maintaining transport selectivity. We hypothesize that structural variations^3,76^ or rearrangements^4^ of the NPC scaffold will leave the spatial variations in Kap avidity relatively unaffected due to simultaneous changes in both FG-motif and FG-Nup density. This may be an important contributing factor in redundancy and securing transport selectivity under a certain degree of structural variation.

Concludingly, our insights into the spatial control of Kap avidity provide a viewpoint of nuclear transport that couples the local microenvironment (as controlled by two types of native FG-Nups in yeast) to the transport behavior of NTRs. The importance of this coupling cannot be understated: the concept of Kap avidity connects the two major classes of nuclear transport models (FG-centric and Kap-centric), and in that way offers a route towards the long-sought consensus between transport models. Our findings and design approach can guide further studies that control explicitly the spatial arrangements of FG-Nups under the presence of a more diverse pallet of NTRs, which ultimately aids in completing our understanding of nuclear transport. Moreover, these principles can further be transferred to the design of well-controlled spatial arrangements of FG-Nups and FG-motifs such as in DNA origami nanostructures^77–80^, which could be used to achieve NPC-like functionality in synthetic cells or serve as highly selective and efficient membranes for molecular sorting^81^.

## Materials and Methods

### Design of the 800-residue artificial ‘NupY’-protein

We designed the NupY-protein according to the design rules in our earlier work on an artificial FG-Nup termed ‘NupX’^31^. NupY represents an ‘average’ of the GLFG-type Nups from *Saccharomyces Cerevisiae*. This class of Nups comprises Nup49, Nup57, Nup100, Nup116, and Nup145N, which are generally bimodal, with a collapsed (low charge content) N-terminal FG-rich domain and an extended (high charge content) FG-devoid C-terminal domain. Incorporating such structural bimodality, NupY comprises a 610-residue long collapsed FG-rich domain and a 190-residue long extended domain devoid of FG-motifs, where the lengths of the two domains were matched to those of Nup100, an essential yeast FG-Nup^35^. FG and GLFG motifs were placed in a 4:3 ratio within the collapsed domain in a repeating pattern of (FG:GLFG:FG:GLFG:FG:GLFG:FG). Since there may not be a full number of repeats of this pattern within the collapsed domain, the final ratio of FG:GLFG motifs in the design could be slightly lower than 4:3. By spacing each FG and GLFG-motif by 13 residues, the average density of FG-motifs and GLFG-motifs within GLFG-Nup collapsed domains was reproduced. Compared to our previous work^31^, the FG-spacing, C/H-value and FG:GLFG ratio more closely match the statistical average found in native GLFG-Nups. Spacer residues within the collapsed domain were based on the collective spacer sequence statistics of yeast GLFG-Nups (Supplementary Table 2). The used Nups are Nup49, Nup57, Nup100, Nup116, and Nup145N for the low C/H ratio domain and Nup100, Nup116, and Nup145N for the extended domain, respectively.

In line with previous work, designs were constructed iteratively and checked for intrinsic disorder using PONDR^82^ and DISOPRED3^83^ until a satisfactory disorder profile was obtained. Further structure prediction using ROBETTA^84,85^, Phyre2^86^ did not yield any consistent or high-confidence structures, confirming the disordered nature of the template protein.

### Sequence analysis of yeast (GL)FG-Nups and artificial FG-proteins

Sequences of native FG-Nups were obtained from UniProt and categorized into high-charge domains or low-charge (collapsed) domains according to definitions by Yamada *et al.*^35^ (Supplementary Table 2). Charge-to-hydrophobic (C/H) amino acid ratios *f*_CH_ for FG-Nup segments were defined as the ratio of the number of charged residues (*N_c_*) divided by the sum of all *N* hydrophobicity values (ϵ_n_ for amino acid *n*). This ratio can be written as:

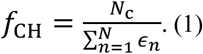

The values for *ϵ_n_* were taken similarly to those used in our 1-BPA computational model^49,87^ (Supplementary Table 4).

### Design of FG-ratio variants and C/H spacing variants

We generated a collection of 800-residue proteins with C/H-values and FG/GLFG-motif spacings (d_FG_) that range from 0.5-12 times (0.024-0.57) and 0.5-8 times (7-117 residues) the average found in the collapsed domains of GLFG-Nups. The design of these variants took place in a multi-step process where variants with different d_FG_-values were designed by mutating the NupY(13;0.05)-protein while preserving the C/H-value of the collapsed domain. The selected d_FG_-variants were then re-used as inputs for generating C/H-variants, where the spacing between FG-motifs was maintained.

To design d_FG_-variants, we first selected pairs of non-aromatic amino acid residues that together comprise a similar hydrophobicity score (using a normalized hydrophobicity scale, Supplementary Table 4) as the amino acids Phe-Gly (‘FG’) or Gly-Leu-Phe-Gly (‘GLFG’). Resulting iso-hydrophobic pairs (AA, MQ, MP, IG, LG IS, LS, VS, VT) were randomly selected to replace all FG and GLFG-motifs (two pairs) within the input sequence. FG and GLFG-motifs were reinserted according to the newly assigned d_FG_-value in a 4:3 ratio, using the same alternating pattern as NupY(13;0.05) (see section on NupY-design). Any residues displaced by the FG and GLFG-motifs at their new position were stored and randomly selected to replace previously inserted iso-hydrophobic pairs, to preserve the C/H of the collapsed domain. The design of NupY(7;0.05) formed an exception to this method, since C/H could only be preserved when creating variants with increased d_FG_. To generate this variant, we applied the design steps for the original NupY(13;0.05) protein, but reduced the spacing to 7 residues. Since the increased density of FG-motifs now yielded a C/H-value that was lower than the GLFG-Nup native average, mutations to the spacers in the cohesive domain were required. The method for generating C/H-variants (next paragraph) was applied to match the C/H-value of the NupY(7;0.05) collapsed domain to that of the other d_FG_-variants.

C/H-ratio variants were designed by employing NupY or d_FG_-variants thereof as templates. Via an in-house code, spacer residues were selected randomly on an iterative basis. Based on the hydrophobicity and charge of the selected amino acid and the direction of the variation (e.g., increasing or decreasing the C/H ratio, resp.), a suitable pool of residues is pre-selected based on the hydrophobicity and charges of the residues. This pool contained disorder-promoting non-aromatic residues^35^ (A, G, Q, S, P, M, T, H, E, K, D) that maintain the protein’s net charge upon substitution. This process was repeated until the C/H of the 1:610-domain reached the assigned value within a certain tolerance.

### Selection of artificial FG-Nups using disorder prediction and coarse-grained modeling

For each combination of C/H and d_FG,_ 10^4^ variants were generated using the methods described earlier. All sequences were checked for intrinsic disorder in the 1:610-domain (the high C/H ratio 611:800 domain was not mutated after the initial template design) using SPOT-Disorder-Single^88^, based on its ability for high-throughput disorder prediction and high accuracy specifically for long amino acid sequences without sequence conservation^89,90^. A subset of 50 proteins with the highest average disorder score in the 1:610 domain was selected. After assessing the similarity between the designs regarding polymer properties using coarse-grained molecular dynamics simulations, a second round of disorder prediction was performed using a local installation of DISOPRED3^83^. A final design was chosen from the three designs with the highest DISOPRED score.

### Coarse-grained modeling of IDPs and folded proteins

Coarse-grained molecular dynamics simulations were carried out using our earlier-developed coarse-grained model for intrinsically disordered proteins^49,91^, where we refer to earlier work (Refs. ^49,87,91^) for an in-depth explanation of the model parameters. Concisely, the model distinguishes between all twenty amino acids and considers hydrophobicity and Coulombic interactions as the main non-bonded interactions, with corrections for residues involved in cation-pi interactions. Backbone interactions (bond stretching, bending, and torsion) are assigned based on the rigidity of the amino acids, where we distinguish between three categories (flexible-glycine, stiff-proline, and all others). We employed a simple coarse-graining procedure for the folded proteins Kap95, ubiquitin, and BSA. Based on the individual crystal structures (Kap95: 3ND2^92^, ubiquitin: 1OTR^93^, BSA: 4F5S^94^), a single bead is placed at the C_α_-position of each residue. A bonded network is then applied, consisting of harmonic potentials with a binding constant *k*_-_= 8000 kJ/mol/nm^2^ for any residues separated by less than 1.2 nm. Non-bonded interactions are set, depending on the type of residue, to represent charged interactions, cation-pi interactions^95^, or volume exclusion. Kap95 binding site regions^55^ that interact specifically with FG-motifs were modeled using a description from earlier work^96^. All simulations in the current work were carried out using the GROMACS molecular dynamics package, versions 2016.3 and 2018.4.

### Genetic optimization of artificial FG-proteins

A subset of artificial FG-proteins (Supplementary Table 1) was selected for gene synthesis. Initially, reverse translation of the amino acid sequence to a genetic sequence pre-optimized for expression in *E.Coli* was done using a freely-available tool (https://www.novoprolabs.com/tools/codon-optimization). Following initial translation, codons were optimized^97^ to replace any remaining rare codons that could reduce expression. Finally, the sequence was checked against the presence of rho-independent terminator regions, regions with high dyad symmetry, highly stable RNA stem-loops, or restriction enzyme sites. These analyses used FindTerm^98^, ARNOLD^99,100^, and CloneManager v.10, respectively. After codon analysis, further codons were added to incorporate an N-terminus Protease cleavage site (for purification), a C-terminus Cys-residue (for end-grafting to surfaces), and a stop codon. These modifications added a’GP’-sequence to the N-terminus, and a C-residue to the C-terminus, causing the final NupY-variants to comprise 803 residues.

### Expression and purification of the NupY variants

NupY proteins were expressed and purified essentially as described for NupX^31^, with the following modifications: The strain used for overexpression (ER2566) also contained pRARE2 (Merck-Millipore), and for NupY(7;0.05) the SP sepharose column was replaced by a 1 ml phenyl sepharose column. To reach high stock concentrations for the LLPS experiments, NupY proteins were precipitated using ethanol containing 40 mM potassium acetate. Pellets were washed three times before being resuspended in low volumes (100-200 µl) of storage buffer (50 mM Tris/HCl pH 7.5, 150 mM NaCl, 6 M GuHCl, 100 µM TCEP) to final concentrations between 10-100 µM, depending on the expression yield of the variants. Proteins were snap-frozen in liquid nitrogen and stored at-80 °C until further use.

### Expression, purification, and labeling of Kap95 and cargoes

Kap95 was expressed and labeled with AZDye647 or AZDye488 (Vector Laboratories, structurally identical to AlexaFluor647and AlexaFluor488, respectively) at the C-terminus using sortase mediated ligation (degree of labeling ∼40%) as described previously^33^. The cargoes IBB-GFP and 2xMBP-IBB were expressed as described previously^81^ with the following modifications for 2xMBP-IBB. An amylose resin was used for affinity purification (NEB, E8021S), the imidazole wash was omitted, and elution was performed using 10 mM maltose. See Supplementary Table 7 for the amino acid sequences of the proteins used in this study.

### Simulating morphology and transport through nanopores

We simulated the morphology of solid-state nanopores coated with our artificial FG-Nups by tethering copies of our FG-Nups to a cylindrical scaffold (diameter of 55 nm) consisting of sterically inert, 3 nm diameter beads, see Figure 5a. FG-Nups were tethered to the interior of the nanopore wall by their C-terminus Cys-residue in a triangular lattice in four rows, using a grafting distance of 5.5 nm. We did not match the grafting distance in these nanopore simulations to the grafting distance in our QCM-D measurements, since QCM-D does not give an unambiguous estimate of the grafting distance^40^. Rather, we chose values for the grafting distance and pore diameter in line with our earlier work on SiN^29,31^, which was confirmed via SPR measurements^31,32^ and agrees well with known estimates of the grafting distance in the nuclear pore complex^16^. Systems were equilibrated by iteratively performing simulations of several ns, where the temperature and timestep increased gradually to 300K and 20 fs, respectively, followed by a longer equilibration run of 2.5×10^7^ steps (500 ns), where data was stored every 5000 steps (0.1 ns). The production runs in NupY-coated nanopores in the absence of cargo took place for 2.5×10^8^ steps (5 µs).

Transport studies were carried out using ten copies each of Kap95, BSA, and ubiquitin, which were inserted into the output structures of the previously mentioned nanopore simulations. Ten copies of each protein were simultaneously pulled into the NupY-coated nanopores until the ten proteins’ center of mass crossed the nanopore’s center. Kap95 was pulled from the top side, whereas BSA and ubiquitin were pulled from the bottom. Each protein type was restrained in the z-direction (but free to move in the lateral direction) while the next set of proteins was pulled into the pore network. A cylindrical compartment, consisting of 3 nm sterically inert beads, with a height of 45 nm and diameter of 90 nm was added on either side of the nanopore: the compartment interacted sterically with the Kap95, BSA, and ubiquitin proteins (Supplementary Table 5) to confine these proteins to the vicinity of the nanopores, but did not interact with FG-Nups. After pulling, relaxation of the FG-Nup network took place for 2.5×10^7^ steps (500 ns) using a decreased Langevin friction coefficient (by increasing the coupling time τ_T_ to 500 ps rather than 50 ps) while restraining the Kap95, BSA, and ubiquitin proteins. Production runs were then performed for 2.5×10^8^ or 2.8575×10^8^ steps (5 µs) for a timestep of 20 or 17.5 fs, respectively (Supplementary Table 3).

### Simulating IDP-Kap95 binding using coarse-grained MD simulations

Kap95 and our artificial FG-Nups were placed in a 45×45×45 nm^3^ periodic box, where the position and orientation of Kap95 were restrained. Based on a cumulative simulation time of approximately 300 microseconds (of 20 replicas of ∼15 microseconds each), we calculated the dissociation constant *K_D_*between Kap95 and FG-Nup-variants using a relation^101^ that considers the fraction of bound configurations and the fraction of configurations where two proteins are in close proximity of each other but not necessarily bound. Following the definition of Jost Lopez *et al*.^101^, we define *V_sub_*, a spherical volume centered around the center-of-mass of Kap95 with radius 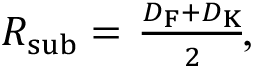, where *D_F_* and *D_K_* are the average largest diameters (largest internal distance between any residue) of the FG-Nup and Kap95, respectively. A binding affinity is then defined as:

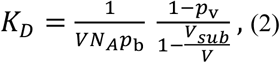

where *p*_b_ is the fraction of bound configurations with the minimum distance *d_ij_* < 0.8 nm^101^, *p_v_* the fraction of configurations where the center of mass of NupY-variants resides in the sub-volume *V_sub_*_-_ centered around Kap95, *N_A_* is Avogadro’s number and *V* the box volume (fixed at 45^3^ nm^3^ throughout). Importantly, we note that this method assumes a binary (1:1) binding interaction, and accounts for multivalent interactions between Kap95 binding sites and FG-motifs from an individual FG-Nup. Since other effects that contribute to binding (e.g., allostery, effects of local concentration variations), are not considered in this method, we used the term binding affinity, rather than avidity when describing the results from this *K_D_*-calculation. We calculated the number of contacts required for the calculation of *p_b_*, using the MDAnalysis Python package^102^, version 1.9.

### Determining pore translocations

We calculated the number of translocations from the z-component of the center-of-mass of the inert proteins (BSA or ubiquitin). We first determined whether the z-coordinate is below the pore membrane (*z* < −10) or above the pore membrane (*z* > 10). The number of crossings was then found by determining the number of downward crossings (*i.e.*, from *z* > 10 to *z* < −10) and upward crossings (from *z* < −10 to *z* > 10) and calculating the sum of crossing events in either direction. We found that the number of translocation events or the scaling with FG-spacing or C/H is insensitive to modest (up to 25%) increases or decreases in the chosen thickness of the pore membrane.

### Calculating time-averaged, axi-radial protein density profiles

The time-averaged (z,r) density profiles were recorded using the `gmx densmap` utility in GROMACS. The number densities for a group of amino acid residues (*e.g.*, FG-Nups, FG-motifs, individual cargo) for each trajectory frame were binned using 0.5 nm-sized bins on a cartesian grid, converted to polar coordinates, and averaged over the azimuthal direction and the total simulation duration. To obtain a radial density profile, an additional averaging step over the pore height (|z| < 10 nm) was performed, and only bins that fell within the pore diameter were considered. Density profiles for cargo molecules were calculated for each molecule and were subsequently averaged rather than calculating the cumulative density profile.

### Mass flux analysis from simulation trajectories

We calculated the mass flux of cargo molecules from their spatial (axi-radial) density and (scalar) velocity distributions. We first obtained axi-radial density maps for individual molecules for all three cargo types, respectively as described in the previous paragraph. Next, we obtained the center-of-mass trajectories for all individual cargo molecules (Kap95, BSA, ubiquitin, resp.) using the GROMACS built-in `gmx traj` and calculated the displacement between each trajectory frame. Using a central differences approach, the velocity vector for each center of molecule’s center of mass could be found for each trajectory frame. The scalar velocity (speed) followed from the norm of the velocity vector. A smoothening step was performed by employing a moving average of 250 frames, which we found to represent the trajectory of the particles well while filtering out high-frequency oscillating movements due to the stochastic dynamics integrator. Finally, the axi-radial velocity distributions were found by mapping the cartesian center of mass coordinates (corresponding to each velocity data point) to an axi-radial coordinate system, and spatially binning the velocities (0.5 nm bin size). The axi-radial distribution of the mass flux was finally obtained by multiplying the density and velocity distributions, where the mass density distribution was interpolated such that it was defined on the same (r,z) grid points as the scalar velocity distribution. Averaging was then performed over the pore dimensions (|z| < 10 nm, r < 27.5 nm) for each molecule and finally over all the copies of a cargo type. In the selectivity score (mass flux ratio of Kap95 over BSA, as a function of d_FG_ and C/H), any ill-defined ratios (*e.g.*, where the BSA mass flux is zero for low C/H) were set to 0 instead.

### QCM-D experiments, materials, sample preparation, and data analysis

For our QCM-D experiments, we employed the Qsense Analyzer platform (Biolin Scientific, Västra Frölunda, Sweden). Gold-coated quartz crystals were employed as substrate for all the coating and binding experiments. Normalized resonance frequency (*Δf_i_/i*) and dissipation (Δ*D_i_*) shifts were acquired at odd harmonics *i* = 3,5,7,9,11, and monitored in real-time using Qsoft (provided by Biolin Scientific with the machine). The seventh harmonic for frequency 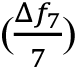 and dissipation (Δ*D*_7_) were chosen for display and analysis. Flow rates for the various experiments were: 30 μL/min for coating the chips with NupY proteins, 20 μL/min for coating with a 1-mercapto-1-undecylte-tra(ethyleneglycol) molecule (MUTEG, Sigma-Aldrich) for passivation of the remaining exposed gold surface after NupY coating, and 60 μL/min when flushing ultrapure BSA (Sigma-Aldrich) or Kap95. All experiments were carried out at room temperature. Prior to the NupY coating step, chips were cleaned using base piranha. Briefly, a solution of 30% Ammonium Hydroxide, 30% Hydrogen Peroxide, and deionized water (DI) in a ratio of 1:1:5 is pre-heated at 75°C in a water bath. The solution was then taken out of the water bath, and chips were immediately submerged and let to react for ∼15-30 minutes. After the base piranha treatment, chips were thoroughly washed in DI and sonicated in pure ethanol for ∼30 minutes. Chips were then blow-dried with a nitrogen flow and mounted into the flow cells, which were previously disassembled, sonicated in 2% sodium dodecyl sulfate (SDS, Sigma-Aldrich), washed in DI, blow-dried with a nitrogen flow, and reassembled. The running buffer for all the experiments was Phosphate-Buffered Saline (PBS, Sigma-Aldrich) at pH 7.4. Before the NupY coating, NupY proteins were incubated with TCEP (Sigma-Aldrich), a reducing agent that breaks the disulfide bridges between the cysteines on the C-terminus of the proteins. FG-Nup proteins were then flushed into the flow cell and immobilized onto the cleaned gold-coated quartz sensor (Figure 2a-b, Supplementary Figure 3a-h) *via* a self-assembly process based on thiol-gold chemistry. The process was monitored in real-time with QCM-D by measuring the resonance frequency shift (Δf), which is proportional to the adsorbed mass, and the change in dissipation factor (ΔD), which scales with the level of hydration and softness of the brush^40^. For the sake of this study and given the known limitations of the QCM-D technique (*e.g.*, mass transport limitations^103^, secondary effects due to entrapped water^40^), we limited ourselves to merely comparing Δf and ΔD among the different experiments, without directly estimating the precise amount of adsorbed dry protein mass per unit area through, *e.g.*, *via* the Sauerbrey relation^104^. Raw data were exported in Excel using Qtools (Biolin Scientific). All analysis, plotting, and fitting were done with custom-written Matlab code. Fit curves shown in Figure 2h-i and Supplementary Figure 6 correspond to two-exponential functions of the form: 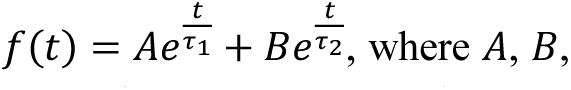 τ_1_, τ_2_ are fit parameters. Extracted parameters from the fits for all NupY variants and Kap95 concentrations are reported in Supplementary Table 6. The brush softness or hydration level was estimated by measuring the dissipation to frequency ratio at the seventh harmonic (Δ*D*_7_/Δf_7_). This is generally expressed^40^ as

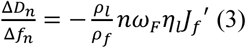

where *ρ_l_* and *ρ_f_* are the densities of the liquid and deposited film, respectively, *n* is the harmonic number, ω_E_ is the angular fundamental resonance frequency, η_F_ is the viscosity of the liquid, and 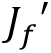 is the elastic component of the compliance of the film. Hence a higher 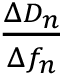 corresponds to softer brushes, as it is directly proportional to the film compliance 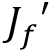 which is a measure of the intrinsic viscosity of the film^105^.

### Phase separation experiments

Condensates of NupY were formed by rapid dilution into denaturant-free buffer (50 mM Tris/HCl pH 7.5, 150 mM NaCl) containing 10% w/v PEG-8000 (Promega) to a final concentration of 200 nM unless specified otherwise (corresponding to a dilution factor of 1:50 to 1:100, depending on the stock concentration of the different variants). For the sedimentation assay, NupY condensates were allowed to form for 1 h before centrifugation at 16.000 rcf for 20 min to pellet the condensates. The supernatant was carefully removed, and the protein contents of the pellet and supernatant were analyzed by SDS-PAGE using Coomassie staining (InstantBlue® Coomassie Protein Stain, ISB1L, abcam), The saturation concentration c_sat_ was taken as the concentration of the soluble protein in the supernatant. To quantify c_sat_, band intensities were compared to a BSA concentration series loaded on the same gel. Sedimentation assays were repeated at least two times. For microscopy, freshly formed condensates were transferred into chambered coverslips (µ-Slide 15-well with ibiTreat surface modification, ibidi, Germany) and allowed to sediment for 1 h prior to imaging. Kap95 and cargo were added at concentrations of 1 µM within 5 min after phase separation had been initiated and prior to loading the sample into the measurement chamber. Brightfield and fluorescence images were acquired on a Nikon A1R confocal laser scanning microscope equipped with a 100x oil-immersion objective. Laser power, detector gain, and pixel dwell times were adjusted to avoid signal saturation for the brightest condensates (usually those of the template NupY(13;0.05)) for each measurement series and kept constant throughout. The signal of Kap95 and cargo within the condensates was measured at the center of the particle where the signal plateaued. Only condensates that were large enough to show a plateauing intensity of the client (approximate size above 2 µm diameter) were considered for the analysis. For FRAP experiments, circular regions of interest were selected at the center of condensates and bleached at maximum laser power for 20 s. No bleaching occurred for particles outside of the selected regions. For influx experiments, condensates were formed as described above and 1 µM of fluorescently labeled Kap95 was added immediately before imaging. For the competition assay, condensates were first incubated with 1 µ M Kap95-Alexa488 for 1 h, after which 1 µM of Kap95-Alexa647 was added to the solution and incubated for 1 h before images were acquired.

For the sedimentation assays shown in Supplementary Figure 7d, 1 μl of NupY protein (60 μM protein in 2 M Gu– HCl, 100 mM Tris–HCl pH 8, purified as described earlier^106^ was pipetted in a low-protein-binding tube and diluted to a final protein concentration of 3 µM in assay buffer (50 mM Tris–HCl and 150 mM NaCl, pH 8.0) containing 0%, 5%, 10% or 20% 1,6-hexanediol, and incubated for 1 hour at room temperature. The insoluble fractions were then separated by centrifugation (17,900g for 10 min at room temperature), separated with SDS-PAGE and stained with Brilliant Blue G overnight. Band intensities were determined with Fiji (Image J, National Institute of Health). Insoluble fractions were calculated compared to the total protein (insoluble + soluble).

## Acknowledgements

This work was financially supported by Dutch Research Council grants OCENW.GROOT.2019.068 (to P.R.O., C.D and L.M.V.), VI.C.192.031 (to L.M.V.), and the ‘Projectruimte’ grant 16PR3242-1 (P.R.O and C.D) of the Netherlands Foundation of Scientific Research Institutes (NWO-I). Computing resources were provided by the University of Groningen CIT, the Berendsen Centre for Multiscale Modeling and the Dutch national e-infrastructure with the support of SURF Cooperative, grant no. EINF-972 (H.W.d.V.). We thank Eva Bertosin, Nils Klughammer and Jaap van der Vis for discussions.

