## Supplementary Information for "Spatial control of karyopherin binding avidity within NPC mimics revealed by designer FG- Nucleoporins"

| C/H | d_FG_ |  |
| --- | --- | --- |
| **0.0478 (1x)** | **7 (0.5x)** | **GPSQIMNSTFGPNNTGNQGLFGAPGSPSTFGTNQTNGTGLFGQAQSSSNFGQGSPLTTGLFGLSLAPQEFGNSAMVQSFGPNTGASTGLFGRKGGQKSFGKFNNIMMGLFGQSSPQTQFGQNSANNTGLFGQALTTGAFGQPFSANNFGLGNNPTSGLFGNGGKVGTFGSGSPGNPGLFGNSGNGQTFGLGLKTGSGLFGNGNHKTSFGAAGASLNFGQNQAGSNGLFGKGGAASNFGSSKQSKNGLFGMQSSGSTFGAQGNNQQGLFGTGGSTTNFGHHQSSTMFGTMINNQQGLFGNQNNLTQFGNTSSQPSGLFGHTQMFGKFGTNVNPLVGLFGGGQPMTSFGAGNIMGQFGTNSQGGSGLFGQVTNGMNFGHSMLNSMGLFGTANSGPSFGMNSSHSTGLFGQIVLPGTFGNGGASHNFGNQNSNSQGLFGMMSSDGNFGPTQSQTSGLFGQTLTPGPFGSGQHASSGLFGMNNGSFQFGAGTTNMQFGNQTGTNSGLFGQGSLAPNFGSPKASTAGLFGSGATAGTFGSSLSNTTGLFGFQYGAGSFGPTSMSGQFGNPPQSTAGLFGAPGGTAPFGNNSAGKSGLFGPSRPNQHFGSPNNTTTGLTKLKSSIGKDFTLAQVNKTTNNASIIEAIPYYNLVKKSSNKRRSYNEDDPRLGENASSAKAKKFQLDREQLNSAGKDDFDIQLLSPHESVFMIMLNAGNLDARLDANNKVESLNSNDAQRSNGENPKPRAMTPLTISSEKDSISSSMKIDPAKHLTEPSQDIEQKKLTTTWKSSDEKPVSVPKSTNSSEEC** |
| 0.1912 (4x) | 7 (0.5x) | GPSQIMNSTFGGNKTGNQGLFGAPGSPSTFGTKQANGTGLFGQQQSSSNFGQGKPTTTGLFGKSLAPQEFGESAMHQSFGHNTGASTGLFGRKEGQKSFGKFNNISMGLFGQSSPQTQFGQNSGNNTGLFGQALTTGAFGSRFSAKRFGLGENPTSGLFGRGGKVGGFGSESGRDHGLFGDSENGQTFGLGLKTGSGLFGNGEHKTSFGATGAELNFGQNGAGSRGLFGKDRAPENFGSSKQGKNGLFGMHSGGSTFGEQGNNQQGLFGQGESETNFGMHQSSKMFGTGINDQQGLFGEQNNLTQFGNTSSQPSGLFGGTQGFGKFGTNVNPLPGLFGGGQPPTSFGARNIGGSFGKNSQGGRGLFGQSDNGMNFGGSMLNSMGLFGTANGGPSFGMNSGHSTGLFGQIVLPGGFGNKDSSTNFGNQNSNSQGLFGPMSSDGNFGPTQSETSGLFGQTLTPRPFGSGQHASSGLFGGENGGFQFGARTTNGHFGNQTGTNSGLFGQGSLAPKFGSPKTSTAGLFGSKATAATFGSSDKRTTGLFGFQHGAGSFGSGSMSGQFGNPPQGTAGLFGSPEGRAPFGNASHRKSGLFGPSRPNQHFGSPENTTTGLTQLKSSIGKDFTLAQVNKTTNNASIIEAIPYYNLVKKSSNKRRSYNEDDPRLGENASSAKAKKFQLDREQLNSAGKDDFDIQLLSPHESVFMIMLNAGNLDARLDANNKVESLNSNDAQRSNGENPKPRAMTPLTISSEKDSISSSMKIDPAKHLTEPSQDIEQKKLTTTWKSSDEKPVSVPKSTNSSEEC |
| 0.3824 (8x) | 7 (0.5x) | GPGQIMNRGFGPKDPGNSGLFGAPESPSTFGTNMTNGDGLFGESQESSEFGTDSHLREGLFGGDAAPQEFGNSMDQQSFGPKSGKSRGLFGRKRRKKSFGKHNKIMPGLFGQSSTQTDFGQKSEENTGLFGQAETTGHFGQPKSADRFGSGENPTSGLFGNGGKVGGFGKGSPGEPGLFGNKDNGQTFGSEMKTGEGLFGNGNHKTSFGAAGSSLNFGSNTKGSKGLFGKGGKKDNFGSSKQDKEGLFGHQSSGDTFGEQEMNQQGLFGKGGSTKRFGDHQSSTTFGTQHSEQGGLFGDHNNHTQFGNRSSQPRGLFGHKQPFGKFGGNVEPPVGLFGKKEPASSFGEGNHMEQFGGDSSGGQGLFGQHGRRPNFGHSMLNRMGLFGTANSKPSFGTNDSHKKGLFGQIMMPKTFGNRKAQTNFGNDNSNSQGLFGMMSSDGNFGDDQRQKSGLFGQGMTPGPFGSEQHDSSGLFGMDKGGFHFGDDSTPMQFGEGGHGNSGLFGQGSLDMKFGGTKTSTTGLFGSGRKHGSFGSGLSNGTGLFGFGSAAGRFGPTGMEGQFGNPRQSGAGLFGAPGETHSFGRKSAGKSGLFGRDRPRQKFGSPNNDTTGSTKLKSSIGKDFTLAQVNKTTNNASIIEAIPYYNLVKKSSNKRRSYNEDDPRLGENASSAKAKKFQLDREQLNSAGKDDFDIQLLSPHESVFMIMLNAGNLDARLDANNKVESLNSNDAQRSNGENPKPRAMTPLTISSEKDSISSSMKIDPAKHLTEPSQDIEQKKLTTTWKSSDEKPVSVPKSTNSSEEC |
| **0.5736 (12x)** | **7 (0.5x)** | **GPSEMHNSDFGGDERRETGLFGAEGSPSSFGTNGTNRKGLFGQGGRSSEFGQRRGQRTGLFGAKEGGQEFGESAQMQSFGSNTGRSSGLFGRKGGQKSFGKFRNIKMGLFGSDGPRTQFGQNEAKKKGLFGQRMSERSFGQPMKGNNFGQKENKDDGLFGNKKKAGMFGKGSTKNPGLFGDSGEGEKFGLDAKDGSGLFGNGKGKSSFGAAGAGHEFGQEQAGKKGLFGKKEDASAFGSKKQDKEGLFGSEESGETFGRQRNESQGLFGTGEKKENFGDHQSEDQFGTPKNNQGGLFGHGKDLSQFGETSEGTDGLFGHTKMFGKFGREQNPLEGLFGGGEHAGPFGTGEPMGQFGTRESGGSGLFGQVDNGGEFGHRTLRSHGLFGTTNSDPEFGMNSSHRSGLFGQGELSRRFGDGDHTGNFGNGNSKSGGLFGRTDEDRKFGTGQSQDGGLFGQRGTKESFGSRSEAKKGLFGDENDSKQFGPRQDKMQFGNQSGSKSGLFGSGGHAPNFGRPKQDGKGLFGSGATAKKFGSSHRKTTGLFGRTSGAKSFGGTSKSEMFGDDPQDSPGLFGAPGDTSPFGEESSKKEGLFGPKRTNQEFGKPKNETTGSTKLKSSIGKDFTLAQVNKTTNNASIIEAIPYYNLVKKSSNKRRSYNEDDPRLGENASSAKAKKFQLDREQLNSAGKDDFDIQLLSPHESVFMIMLNAGNLDARLDANNKVESLNSNDAQRSNGENPKPRAMTPLTISSEKDSISSSMKIDPAKHLTEPSQDIEQKKLTTTWKSSDEKPVSVPKSTNSSEEC** |
| 0.0239 (0.5x) | 13 (1x) | GPTPGPSANSHAMGMFGAASHASTNSFSGSGLFGSNTQMAANMLTQMFGQPAGSNNTMNTSQGLFGGMNQTFMVKMSSNFGNGNGRGQNQQAMLGLFGQPQTHPTGSIITHFGGLSTSKSQSSQVAFGSSTQMAGMHMQTNGLFGSMNNNTNLSMNPMFGTAVSQAAVASYAMGLFGISSGHTQTGAQNVFGPQPAANSPQSMTGGLFGHANGTTKANMNQPFGAAFSPAGSMAFPNFGQLMANTANGPMGAGLFGPPNQTASPSNTPPFGQNPLPPVPQTQAPGLFGSAQSHANTTNGNMFGHTMSGTKGNMINAGLFGQPQNQGGMQTTQIFGTALPAPMQMMNSQFGSNRTMAAHMQATGGLFGALMQSSLLNPQATFGQSNQASHLSMQMMGLFGAGAMTTQSNSNMMFGGSQTAQPSSMGNMGLFGTHMAGPALAMKMGFGMSQQSMPFTGGQVFGLNLQQMQQGTNTTGLFGTNVLTMAFQAPQPFGGKLGGSSQSNSNLGLFGSQTMGTPLGQNSNFGAGQQSPNSSATGGGLFGANSPMSPANGQATFGQNGNAMMMPMGKPFGTQTTMNNASSNGIGLFGAFANSQQAPMSNTFGSTQPTSASQNLKSSIGKDFTLAQVNKTTNNASIIEAIPYYNLVKKSSNKRRSYNEDDPRLGENASSAKAKKFQLDREQLNSAGKDDFDIQLLSPHESVFMIMLNAGNLDARLDANNKVESLNSNDAQRSNGENPKPRAMTPLTISSEKDSISSSMKIDPAKHLTEPSQDIEQKKLTTTWKSSDEKPVSVPKSTNSSEEC |
| **0.0478 (1x)** | **13 (1x)** | **GPSSGPSANSGGTNQFGTASTSGNNNFSGSGLFGSNTQPAKNQLSTNFGQPAGSNNKMNTSQGLFGGQNGTFPVNSGSNFGNNNGRGQNQQAQLGLFGSPQGTGTGSIITNFGGLGGSKSQGSQVQFGSSTQASNPNTQTNGLFGSQNNNTNLSTNSQFGTAVSQSAVASYAKGLFGISSGNTQTGAQNVFGTQNAANSNTSTNGGLFGNANGNNKANTNQPFGAAFSQAGSQTFPNFGQLAPNTANGPQGAGLFGQGNQTSSPSNSSAFGTNTLPGVPQTQKSGLFGSANSSANGENGNMFGNGTSGTKGNPINAGLFGQPSNKGGKQTTSIFGTALGAGSQNSNSNFGSNRTMTSGSQATNGLFGQLQTSGLLNTQTTFGQSNTSSGLSGQMGGLFGAGTQTNQSNSNASFGGSQTTQPSSAGNNGLFGTSAGGNPLATNQKFGTSQQSPQFTNGQVFGLNLQNTGQGTNTTGLFGTNVLNKSFQAPQNFGNKLNGSSNSNSNLGLFGSGTTGTPLGQNSNFGAGQQSPNSSATGGGLFGSNSPASQTNGSATFGTNGNNGMMPMGKTFGTSTNNNNANSNGIGLFGTFNNSQQTPNGNTFGSTQNTNPSTNLKSSIGKDFTLAQVNKTTNNASIIEAIPYYNLVKKSSNKRRSYNEDDPRLGENASSAKAKKFQLDREQLNSAGKDDFDIQLLSPHESVFMIMLNAGNLDARLDANNKVESLNSNDAQRSNGENPKPRAMTPLTISSEKDSISSSMKIDPAKHLTEPSQDIEQKKLTTTWKSSDEKPVSVPKSTNSSEEC** |
| **0.0956 (2x)** | **13 (1x)** | **GPSSGPSANSGGTNQFGTASTSGNNNHSGSGLFGENTQPAKNQLSTNFGQPAGSRNKMNTSQGLFGGQNGTDPVNSGSNFGNNNGRGQNQQAQLGLFGSPQGTGTGSGITNFGGLGGSKSQGGQVKFGSSTQASNPNTQTNGLFGSRNNNTNLSTNSQFGTAVSQEAVASYAKGLFGISSGNTQTGAQNVFGTQNAANSNTETNGGLFGNANGNNKANTNQPFGAAFSQAGSQTFPNFGQLAPNTANGPQGAGLFGQGNQTSSPSNSSAFGTNTLPGVPQTRKSGLFGSANSSANGENGNMFGNGTSGTKDNPINAGLFGQPSNKGGKQTTSIFGTALGAGSQNSNSNFGSNRTMTSGSQATNGLFGQLQTSGLLNTQTTFGQSNTSSGLSGQMGGLFGAGTQTNQSNSNASFGGSQTTQPSSAGNNGLFGTSAGGEPLATNQKFGTSQQGPQTTNGSVFGSNAQNTGSGTNTDGLFGTNMLNKSFQAPQKFGNKLNGSSNSNSNLGLFGSKTTGTPLGQNSNFGAGQQSPNSSATGGGLFGGNSPASQTNGSATFGTNGNNGMMPMGKTFGTSTNNNNANSNGTGLFGTFNDSQKSPNGNTFGSTQNTNPSTNLKSSIGKDFTLAQVNKTTNNASIIEAIPYYNLVKKSSNKRRSYNEDDPRLGENASSAKAKKFQLDREQLNSAGKDDFDIQLLSPHESVFMIMLNAGNLDARLDANNKVESLNSNDAQRSNGENPKPRAMTPLTISSEKDSISSSMKIDPAKHLTEPSQDIEQKKLTTTWKSSDEKPVSVPKSTNSSEEC** |
| **0.1912 (4x)** | **13 (1x)** | **GPSSGPSANSGGTNQFGTASTSGNNNFGGSGLFGSNSQAAKNQASTNFGQPAGSNNKMNTEQGLFGGQNKSFPVNSGSRFGNNNHRGQNQQAQPGLFGGPQGTETGSIITPFGGLGEDKSQGSQVQFGSSTRASNPNTQKNGLFGSQNNNSNMSSNSQFGTAVSTSAVASYEKGLFGISSGNSHTGMQGVFGTQNMANSNTSTPGGLFGNENGNKKANSDSPFGAAESQGGSQSFPNFGQMKPNTANGPQDHGLFGHGNQTSRPRNSSGFGTNTLPGVPSTQKSGLFGSANSEAEKENGNMFGRGSSGRKGNPIKAGLFGQPSNKGGKQTTKMFGTALGAGSQNSNSNFGGNRTMTSGSQADNGLFGGLQSSGGLNTQTTFGQQNTEEGLSGQMGGLFGAGTQTNQGNSNASFGGPQTKQPSSAGNNGLFGTSAGGNPLAGNQKFGTSQQSPQFTNGQTFGLNRQNTDGGTDSSGLFGGEVQNKSFQPPQNFGNKLNKSSNSNSNLGLFGSDTTGTPLGQNSAFGAGQQSPNSKGTGEGLFGRESPRSQTNGSATFGTNGNNGMMPMGKSFGSSTNNNKPNSNGIGLFGTFNNSQQTPNGDTFGSKHRTNPSDNLKSSIGKDFTLAQVNKTTNNASIIEAIPYYNLVKKSSNKRRSYNEDDPRLGENASSAKAKKFQLDREQLNSAGKDDFDIQLLSPHESVFMIMLNAGNLDARLDANNKVESLNSNDAQRSNGENPKPRAMTPLTISSEKDSISSSMKIDPAKHLTEPSQDIEQKKLTTTWKSSDEKPVSVPKSTNSSEEC** |
| **0.3824 (8x)** | **13 (1x)** | **GPSEGPSAKKGGTKGFGTSSTSGNNDFSGEGLFGSNEQPAKNQGATNFGQPQKSRNKTETSQGLFGKSNEGFPSNGGSDFGNENGRGQNQGSELGLFGKTQGEGTGKIITRFGGLGGSKSSESQSSFGSSPSASNPNTQSNGLFGDRNENTRRSTNSSFGTAVSQKAPASSAKGLFGISSGNTQDERTNVFGDQNAADDNTETEGGLFGNMNERNKANTNHPFGAAPSQHGSQGFPEFGQAQQNTANKQTDAGLFGHRERTSSPKKSDAFGTNTLKEVSQTGKSGLFGSHNSSANGEKGNMFGNGESGTKGNPKEAGLFGQSKNKGKKQKGSIFGTATGAGSQRSNEGFGSNRTDDEGSQGTNGLFGQEQTSELLNDQSTFGQSNTSGELSGGMGGLFGTGTHTEKSNSNASFGDGQTKQPKSTGNNGLFGTSTKGNQLSTNQKFGGSQQSPTTTNGHVFGMNGHRTGGGTRTTGLFGEKVKNKGFQAPQDFGNKHEDSSNSNSNAGLFGSGRTKRPLGSNSNFGTGQQRPNRSQSGGGLFGRKRTGSQREGSAGFGTNGNNGMMPMGKTFGTSENNNNANSEGQGLFGSFNNSQETPNGNTFGSKGNTKGETNLKSSIGKDFTLAQVNKTTNNASIIEAIPYYNLVKKSSNKRRSYNEDDPRLGENASSAKAKKFQLDREQLNSAGKDDFDIQLLSPHESVFMIMLNAGNLDARLDANNKVESLNSNDAQRSNGENPKPRAMTPLTISSEKDSISSSMKIDPAKHLTEPSQDIEQKKLTTTWKSSDEKPVSVPKSTNSSEEC** |
| 0.5736 (12x) | 13 (1x) | GPKKRPSTRDGRTNQFGTASGSKNNNKSDSGLFGSNTHPTKNSLDRRFGGRAEDNDKMDTSEGLFGGKEGTTPDNSGSNFGRDKERGQEDQAKPGLFGSPTGDMTGSSTSEFGRLRKSKSKDRQPQFGSSDGRSDGKTQTRGLFGEEDNNTNLSKNKEFGTAVGSGDVGSKAKGLFGSSSGETQTDAQNGFGTQNHANSEEGTNEGLFGNANMNRKARTNQSFGGAFKQAGSQEQPKFGGLAQEGGNGSSKAGLFGQGKQGSSPENSSDFGENRSPEVRKTQKSGLFGSANSSANDENDKMFGNGTPKTKQNPINAGLFGQDENSKEPQDTSIFGRPLRQGSRNSNSNFGGNRTMRSGGHASNGLFGGLTTSGLGNTQRTFGHERTSSGKGGQRKGLFGAGTQRRQSKSRAGFGGDETEKPSKTRNRGLFGGSQGGRPKRSNQKFGSSSQSQQATREQVFGTNAQNTEQGTNTTGLFGTNPLEKDFQQSGRFGNKQRESSNGRGNLGLFGTEMTDEALGQNSRFGAGTQDPNGKRDGDGLFGSPKPASSTEDSATFGENGNERSHPMGKDFGSGGENNNHESEEIGLFGTHNESEHKRKGNKFGSEQPENPSTNLKSSIGKDFTLAQVNKTTNNASIIEAIPYYNLVKKSSNKRRSYNEDDPRLGENASSAKAKKFQLDREQLNSAGKDDFDIQLLSPHESVFMIMLNAGNLDARLDANNKVESLNSNDAQRSNGENPKPRAMTPLTISSEKDSISSSMKIDPAKHLTEPSQDIEQKKLTTTWKSSDEKPVSVPKSTNSSEEC |
| **0.0478 (1x)** | **26 (2x)** | **GPSSGPSANSGGTNQAATASTSGNNNFSFGSVISSNTQPAKNQLSTNVSQPAGSNNGLFGSQSLMPGQNGTFPVNSGSNGSNNNGRFGNQQAQLLNTQSPQGTGTGSIITNMPGGLFGKSQGSQVQSASSTQASNPNTQTNPMLFGQNNNTNLSTNSQVSTAVSQSAVASYAGLFGSISSGNTQTGAQNVGQTQNAANSNTSFGGKSVLNANGNNKANTNQPQNAAFSQAFGQTFPNIGQLAPNTANGPQGATVMPQGGLFGSPSNSSAMQTNTLPGVPQTQKSSVLGFGNSSANGENGNMTNNGTSGTKGNPINAGLFGQPSNKGGKQTTSIQTTALGAGSQNSNFGNQSNRTMTSGSQATNLGGSQLQTSGLGLFGTTSNQSNTSSGLSGQMGKMNTAGTQTFGSNSNASSAGSQTTQPSSAGNNNQTSTFGGGNPLATNQKMPTSQQSPQFTNGQVMGLFGQNTGQGTNTTSLAATNVLNKSFQAPQFGTNKLNGSSNSNSNLNNNASGTTGTPLGLFGNGSAGQQSPNSSATGGPLNLSNSPASFGNGSATAATNGNNGMMPMGKTGSTSTNGLFGNSNGIGQNSTFNNSQQTPNGNTNVSTFGTNPSTNLKSSIGKDFTLAQVNKTTNNASIIEAIPYYNLVKKSSNKRRSYNEDDPRLGENASSAKAKKFQLDREQLNSAGKDDFDIQLLSPHESVFMIMLNAGNLDARLDANNKVESLNSNDAQRSNGENPKPRAMTPLTISSEKDSISSSMKIDPAKHLTEPSQDIEQKKLTTTWKSSDEKPVSVPKSTNSSEEC** |
| 0.0239 (0.5x) | 52 (4x) | GPSSGPSMSSGGTNQNHTMSQMGNNNFQSSITNLSNTQPAKSQLAAALGQPMMSFGQMPTSQPMMQGANMTFAVNSGSTLGNTHGTMMQQQAQLGNKLSMQMPGTGMIGLFGGGLGAMKTMGSMVQQTSSTPASTPMTQTNGIMQSQPNNTNLSMNSALGTAVSFGAVMSYAKALMPISSGNPQTHAQNVLSTQNMANMATQMNMGLLSNANGNNKANGLFGQPAMFSQAGSQTFAPVSQLMMAPANGAAGAPMVTQTAQASPPAMHQAMPMNTFGGVPQTQKSAAMMHMQSSANGMSGNMMQNGMSGTQHNPINMGLLSAPSNKGGTGLFGIVSTMLGAPSQMSNQNMPSNRPMTQGSMATNQMTQQLQTAGLLNTQTPMQQSFGQMGLHGQMMAMSTAGTMTNAMNSQASLSPSAQTQPSSAGPNMLVSASAGPNPFGTHQSNMTSQQSPQFTSGQVLPLNLQMPGMGTNQQPMMQQNVLNKSFQAPQHIGLFGNGSSNSQSALMLLGSGTQGTPLGMNSGLAAGQPQANSAATGGQMLSMMMAAPFGNQSAPVSTNGGSAMMPMGHTVMTSTNNNAMNSQTITNQPTFNNSQQTPPGNPGLFGQNTMPHTNLKSSIGKDFTLAQVNKTTNNASIIEAIPYYNLVKKSSNKRRSYNEDDPRLGENASSAKAKKFQLDREQLNSAGKDDFDIQLLSPHESVFMIMLNAGNLDARLDANNKVESLNSNDAQRSNGENPKPRAMTPLTISSEKDSISSSMKIDPAKHLTEPSQDIEQKKLTTTWKSSDEKPVSVPKSTNSSEEC |
| **0.0478 (1x)** | **52 (4x)** | **GPSSGPSANSGGTNQNTTASTSGNNNFSGSITNLSNTQPAKNQLSTNLGQPAGSFGKMNTSQPMMQGQNGTFPVNSGSNLGNNNGRGQNQQAQLGNKLSPQGTGTGSIGLFGGGLGGSKSQGSQVQQTSSTQASNPNTQTNGIMQSQNNNTNLSTNSQLGTAVSFGAVASYAKGLMPISSGNTQTGAQNVLSTQNAANSNTSTNGGLLSNANGNNKANGLFGQSAAFSQAGSQTFPNVSQLAPNTANGPQGAPMVTQGNQTSSPSNSSAMPTNTFGGVPQTQKSAAMQSANSSANGENGNMMQNGTSGTKGNPINAGLLSQPSNKGGKGLFGIVSTALGAGSQNSNSNMPSNRTMTSGSQATNQTTSQLQTSGLLNTQTTMQQSFGSSGLSGQMGAASTAGTQTNQSNSNASLSGSQTTQPSSAGNNSLVSTSAGGNPFGTNQKNNTSQQSPQFTNGQVLPLNLQNTGQGTNTTPMMQTNVLNKSFQAPQNIGLFGNGSSNSNSNLGLLGSGTTGTPLGQNSNLAAGQQSPNSSATGGQMLSSNSPASFGNGSATVSTNGNNGMMPMGKTVTTSTNNNNANSNGITNQPTFNNSQQTPNGNTGLFGQNTNPSTNLKSSIGKDFTLAQVNKTTNNASIIEAIPYYNLVKKSSNKRRSYNEDDPRLGENASSAKAKKFQLDREQLNSAGKDDFDIQLLSPHESVFMIMLNAGNLDARLDANNKVESLNSNDAQRSNGENPKPRAMTPLTISSEKDSISSSMKIDPAKHLTEPSQDIEQKKLTTTWKSSDEKPVSVPKSTNSSEEC** |
| 0.1912 (4x) | 52 (4x) | GPSSAQSANSGGTNQNTTASTSGNNNSSGSISNGSNTGPAKDQLTTNLGQPAGSFGKMNTSQPMMQGQNGTFPVNSGSNLGERNGRGQDEQAQLGNKLSPQGTGTGSIGLFGGGAGGSKSQESQVQQTSSTGASDPNTQTNGIMQSQNNNTNLSTNSSLGTHVSFGPVASYAKGLMPIQSGNTQTGEQNTLRTQNAANSNTETNGGLLSNANGNKKARGLFGQSGAFSQAGDQTFPNVSQLAPNTANGPRGEPDVTQGNQTSSPSNSSKMPGNTFGGVPQTTKGTAMQSANSSARGENGNMMQNETSGSKPNHINAGLLSQGSNKGKKGLFGPVSTALGAGSHKSNSNMKSNRTMTSPSQATNQTTSHLQTGGLLKTQTTMQQSFGSSGLGGQMGASSTAGTQTNESNSNASLSGSHTTQPSSMGENSLVSTSAGGNPFGTNQKNRTSQQGPQFTNGQVLPGQLSRTGQGTNTTPPMQTNSLNKSPQAPQKIGLFGNGSSNSNSNLGLLRSGSTKTPLGHKSNLAAGQESPGQSATGEQMLESNSPESFGNGSATVSTNGDNGMMPMDKTVKRSTNNENANSRRPGNQPTFNNSQKTPNGNTGLFGQNTNPSTELKSSIGKDFTLAQVNKTTNNASIIEAIPYYNLVKKSSNKRRSYNEDDPRLGENASSAKAKKFQLDREQLNSAGKDDFDIQLLSPHESVFMIMLNAGNLDARLDANNKVESLNSNDAQRSNGENPKPRAMTPLTISSEKDSISSSMKIDPAKHLTEPSQDIEQKKLTTTWKS |
| 0.3824 (8x) | 52 (4x) | GPGEKPSANDGGTNQNTTQSTGENDNFSGKQTNQSNSTPAKNQAETELGQPTGSFGKMNTSQPMTQGQNGKFPVNSGSRLGDNNGRGRNDQAQLGNKLSPTGTGEGSTGLFGKDLGESKSQRSQVQETSSSQASNPETTTEGIMDGQNNNTRKGKNSTKETAVSFGAVASAAKKLMPISSGNDQTRAQNVDGGQMAANSNEDGNGGSLSNANGNNKQNGLFGQREAFSQAERHTFPNRSQAAPNTANGPHGEPMMTQGNQTSSPSNSSAMPTNKFGGSQQTQKSAAHSSANSGKNGENGNMMQDGTSGTKGRPGEGGLSDQPSNKGDKGLFGIVGTAKGAGSQNKNSNMPSNRTDTSGSQATNQTGSQTQTSGDLNTQETMDQGFGKSKPSGQMGKAKTADTDTRQSNRNRSLSGSQTRQPSSPGNNSLASTDAGGNPFGKNQKNNTRQRSPQFSNGEKLPLETQNTRQGTNTTEMMKTRVANKSTQAPQNTGLFGNGSGNSNSNMGLLGSGTTDTPLGQKDNTSAGQQSPNSEAGKGQMLSSESPAKFGRGSATPEGKKNEGMMGMGKSVTTSTNNNDANSNKITNGPSSRNSRQTENGRTGLFGGNTNPSTELKSSIGKDFTLAQVNKTTNNASIIEAIPYYNLVKKSSNKRRSYNEDDPRLGENASSAKAKKFQLDREQLNSAGKDDFDIQLLSPHESVFMIMLNAGNLDARLDANNKVESLNSNDAQRSNGENPKPRAMTPLTISSEKDSISSSMKIDPAKHLTEPSQDIEQKKLTTTWKSSDEKPVSVPKSTNSSEEC |
| 0.5736 (12x) | 52 (4x) | GPSSGPSANSEETEQRSTRDTKEDDNMSGSMGELDKTGPRKNTLSTNLEQPAGSFGKMNTGQPMMKKRNGTFPVNDGEELDNNRRRGQEKGHSLKEKLGPQKTRGGSIGLFGDRSGESKRQRSSVQQTSSTRASNSETKTEGIMQSQKNNRNLSRNDQTETHEKFGAVASYAKGLGPRSGEETSGKASNMLSSTNGANENTSKKGKLLEDHDGNNKARGLFGESSTMSQSKSSTFQNTSQEAPKTGNGKQRAPMGGQGNETGSPEEESKPPSNTFGGVPQTQKGATMQSSNDSAKGENGNAMQNGTKEEKKKPINAKRLSQPSEKGGKGLFGPVSTALGAGESKEESNMDKNRRATSGSQAGEQTTSQLQTSRLPDTTSDMQQGFGSSGLSGGMDKPSTAGTQTNQSNSNARSSKSQTTQTSSAGNESLVSTGAKERPFGGNQKNDDSQEEDQFTNGQVLPTNMRETGQGTNKTPRQQENQLNKSKQRPQNSGLFGNRSSKGNKNLKALGSGETRTHSGQNSETATRQESPEGGATDGSHLSKNGPAEFGNGKATGGSNENRGMMGMDKGVTTGKNNNNAKSKGQTNQKTGNNGTSTTNGETGLFGGNEEPESNLKSSIGKDFTLAQVNKTTNNASIIEAIPYYNLVKKSSNKRRSYNEDDPRLGENASSAKAKKFQLDREQLNSAGKDDFDIQLLSPHESVFMIMLNAGNLDARLDANNKVESLNSNDAQRSNGENPKPRAMTPLTISSEKDSISSSMKIDPAKHLTEPSQDIEQKKLTTTWKSSDEKPVSVPKSTNSSEEC |
| 0.0239 (0.5x) | 104 (8x) | GPQSAPSANSMGTNQLMMASTSAQMGFTAQAAISHNTQAMPPMLSPNLGMPAGSNQHMNPSMNMLLGQHGPFPVMSSSSAMHNNQRTQNQQMQLPMMASPMMTGHGFGITNISGLGGSPSQGSQVQVSATPQASHMNMQTNAALSPQGNNTNLSTATQMPTAVMMSMVMSYASQMMQISSGNTQTMAQGVLTQQNMANSNTSTSMQMMPNMNGLFGANTNQPMPAAFSAAGSMPFPNSIPLAPNTANGPQQAPMLTQGNPTSSPANQSAMPTPTLAGVAMTQGTPMLGSANTMANGENGTMIPNGTSMTQGNPINAMVRQFGSSKGGKATTSILSTALTAGSPATHSNLPMNRPMTSMSQAMNSMMPQLPTMPLLQTQTTQPQMPTSSGLQGQMHPVMQAGAMAGQSNSNAALPGSMTAQPSSAGNGLFGTTSAGQNPLATNMKMAMSAQSPQFMMGAVMPLNLPNTGQGHNTTPMMQTNVLPPPFQAPKNISNTLQGTHPSNSNLTLISTPTTGHPLGANMNMQMGQASPNSSFGGGGLAASHSPMPPTNPSATLSQHGNNTMMPMPAAMQTSTNAAPMNSNSIQNAKPFNNSAPPMNGNTISSQQQTMMPPNLKSSIGKDFTLAQVNKTTNNASIIEAIPYYNLVKKSSNKRRSYNEDDPRLGENASSAKAKKFQLDREQLNSAGKDDFDIQLLSPHESVFMIMLNAGNLDARLDANNKVESLNSNDAQRSNGENPKPRAMTPLTISSEKDSISSSMKIDPAKHLTEPSQDIEQKKLTTTWKSSDEKPVSVPKSTNSSEEC |
| **0.0478 (1x)** | **104 (8x)** | **GPSSGPSANSGGTNQLSTASTSGNNNFSGSAAISSNTQPAKNQLSTNLGQPAGSNNKMNTSQNSLLGQNGTFPVNSGSNATNNNGRGQNQQAQLPMAASPQGTGTGFGITNISGLGGSKSQGSQVQVSSSTQASNPNTQTNAALSSQNNNTNLSTNSQMPTAVSQSAVASYAKQMMQISSGNTQTGAQNVLSTQNAANSNTSTNGQMMPNANGLFGANTNQPMPAAFSQAGSQTFPNSIQLAPNTANGPQGAPMLSQGNQTSSPSNSSAMPTNTLPGVPQTQKSPMLGSANSSANGENGNMISNGTSGTKGNPINATVMQFGSNKGGKQTTSILSTALGAGSQNSNSNLGSNRTMTSGSQATNAAMPQLQTSGLLNTQTTQPQSNTSSGLSGQMGTVMQAGTQTNQSNSNASLSGSQTTQPSSAGNGLFGGTSAGGNPLATNQKMQTSQQSPQFTNGQVMPLNLQNTGQGTNTTPMMQTNVLNKSFQAPQNISNKLNGSSNSNSNLGLISSGTTGTPLGQNSNMQAGQQSPNSSFGGGGLAASNSPASQTNGSATLSTNGNNGMMPMGKTMQTSTNNNNANSNGIGNNKTFNNSQQTPNGNTISSTQNTNPSTNLKSSIGKDFTLAQVNKTTNNASIIEAIPYYNLVKKSSNKRRSYNEDDPRLGENASSAKAKKFQLDREQLNSAGKDDFDIQLLSPHESVFMIMLNAGNLDARLDANNKVESLNSNDAQRSNGENPKPRAMTPLTISSEKDSISSSMKIDPAKHLTEPSQDIEQKKLTTTWKSSDEKPVSVPKSTNSSEEC** |
| 0.1912 (4x) | 104 (8x) | GPSSGPGADSRGTNQLSTASTSGENNFSGSAAISSNTKPAKNQLSGNLGQPAGSNNKMNTSQNSLLGQNGSFPVNSGSNATNNNGRGQNQQAQLPMAASPQGTMTDFGPTNISGLGGSKDQGSTVQVSSSTQAKNPNTQTNPQLQSQNNNTELSTKSQHPTAVSQSAVASAAKGMMQISSGNTSTGAQNVLSTQEAAESNTSRNGQMQPNARGLFGANTNQPMQTDFSQSGSQSSPNSIELAPNTSKRPQGAPMQSQGNQTSSPSNSSSMPTNTLPGQGQTQKSPMLGSPNSSGDGENGNMISNGTEGTKGNPINASVMQFGSNKGGKQTTSHLSTPLGAGSQRHNSNLGRNRTMTSGSQATNAAMPQLQDSGLLNTQSTQPQSNSESKLSDQMGTVMQAGTQKNQSNSNMSLSGSQTTQPSSAGNGLFGGTKARGKPLATNSKMMTSHQSPQFSNGQMMPLNGQNTGQGDETTPMMQKNVMNKSFQASQNKSNKLNGSSNHNSNLGLISSGTTGTPLGHNSNMQAEQQSPNSSFGGEDLDASNSPAGQTNGSAELSTNGNPGMDSMGKTMQTSRNNNKAKSNGIGNRKTFNNGQQTPNGNTISSSQNTRPSTNLKSSIGKDFTLAQVNKTTNNASIIEAIPYYNLVKKSSNKRRSYNEDDPRLGENASSAKAKKFQLDREQLNSAGKDDFDIQLLSPHESVFMIMLNAGNLDARLDANNKVESLNSNDAQRSNGENPKPRAMTPLTISSEKDSISSSMKIDPAKHLTEPSQDIEQKKLTTTWKSSDEKPVSVPKSTNSSEEC |
| 0.3824 (8x) | 104 (8x) | GPSRGPSANGKGTNQGPTASTSENGNFSGSKQISENEQSAKEQLSTNMGQGGGDNNKMDTGQNSHMGQNRGSQVNSTSSAGNNRGRGQNQGAELPMAPSTQGTGTPFGDTNISGHREHKRQGSQVTGSSSTTASNPRTQDNAALKSGNRKTNSETNGQMPTQGSQSEVPSAAKQMMQIGSGNTQTGATRVLSSQRAAEENTSGNGQMAPNSRGLFGANSNQPMKAAFRRAGSGTFPNGSQLATNTMNGPQGAPMLSQENQGSKRSNSSKMPDRTLPKVPQDQKSPMTGQPNSSANGEDGESESNGTSGGKGSPMRARMHQFGKRKGGKQTTSQGSTASGAGKDESEGNLGKERTTTSGETQTEAAMDQLQTSGLLNTHTKHPQSNTDKGLSEHMGSDGSAGTQGNQSNSKASLSGSQTKKPSRRENGLFGGDSAGGKPGATNQKMQTDQGSPSTTNRQVPDLRPGNTRQGSNTRPSMTPEMLNKSFQAPQNISNKLNGGSKSNSNLTQIESGTEGTPLEQNSEMQMGQQGENSSFGKGGQRGSRSPESSTNGSATPMDNGNNGMMPMKKTGQTSTNNENANSNGIGKNKTFNNSQQTPDGDSTSGTGETNPSTNLKSSIGKDFTLAQVNKTTNNASIIEAIPYYNLVKKSSNKRRSYNEDDPRLGENASSAKAKKFQLDREQLNSAGKDDFDIQLLSPHESVFMIMLNAGNLDARLDANNKVESLNSNDAQRSNGENPKPRAMTPLTISSEKDSISSSMKIDPAKHLTEPSQDIEQKKLTTTWKSSDEKPVSVPKSTNSSEEC |
| **0.5736 (12x)** | **104 (8x)** | **GPSKESGADSGGTKQLSTAEDSEKDKFSDSSGHSSETQPAKDEMGTNPKQPHESNNKPNTSQNSLTKQNGTMPVNSGGRDTNNNGRKQKHEAQSGMAPSPQGTGSKFGIENESRLGGSKRHDSHVQTSSSEQASEPNSQTNDTLGSQNENGELSTNSTMSSRDEQSHHASYAKQMMQISRGRTQTKAQNPLSTQDAPNSETSGEGQKGPKANGLFGARGDQPMPAAMSQQGSQTKPNSIHLAPNSARKPRGAPMLRHKNKTSKPSESSAMPDNTLPGVPSGHKSQQLGSANSSGNGENDEMTENGTGGTKDNTIEAKPMQFGREKGKKSTTKGLDTALGADSQNKNSNMGSRRTATSKSGATNHRMPQLQTSKLARTSTTGTQSNTSSGLDRQTGTVMQEGSQSNESRSDASASRSRKSQPSSQENGLFGGRKAGGKPEDTNSKMTTERRSPQFTNEQVSPQRLEETGQETKTESHMEGNTLNKDEQAPQNHSNKLDGSSRRKSRAESISSGTTGTELGQNKNSHHDQQKPNDSFGRGGMAAERSHQSQTRGSATASKKGEERMQPMGKTHQTSTNNDNANSNRKEEEKTPNNKDKRPNGRTAESTQNGDPSTNLKSSIGKDFTLAQVNKTTNNASIIEAIPYYNLVKKSSNKRRSYNEDDPRLGENASSAKAKKFQLDREQLNSAGKDDFDIQLLSPHESVFMIMLNAGNLDARLDANNKVESLNSNDAQRSNGENPKPRAMTPLTISSEKDSISSSMKIDPAKHLTEPSQDIEQKKLTTTWKSSDEKPVSVPKSTNSSEEC** |

Supplementary Table 1 – Amino acid sequences of artificial FG-Nups used in simulations and experiments. The final amino acid sequences include a Cys-residue at the C-terminus, and a Gly-Pro-motif that remains after cleaving off the Protease tag at the N-terminus, putting the total sequence length at 803 residues. Sequences optimized for expression and purified for use in the experiments in this work are highlighted using bold text.

| **FG-Nup** | **Residues** | **C/H** | **Average FG/GLFG-motif spacing** |
| --- | --- | --- | --- |
| Nup159 | 387–1071 | 0.51 | 24.42 |
| Nup42 | 1–382 | 0.071 | 11.17 |
| Nup116-CD | 172–764 | 0.042 | 12.61 |
| Nup116-ED | 765–960 | 0.83 | 196* |
| Nup100-CD | 1–610 | 0.042 | 11.1 |
| Nup100-ED | 611–800 | 0.56 | 190* |
| Nsp1-CD | 1–186 | 0.041 | 11.4 |
| Nsp1-ED | 187–617 | 0.748 | 18.6 |
| Nup2 | 120-600 | 0.600 | 34.9 |
| Nup49 | 1–251 | 0.051 | 11.2 |
| Nup57 | 1–255 | 0.05 | 12.81 |
| Nup145N-CD | 1–242 | 0.06 | 17.17 |
| Nup145N-ED | 243–433 | 0.49 | 191* |
| Nup1-ED | 220–797 | 0.574 | 46.33 |
| Nup1-CD | 798–1076 | 0.072 | 38.14 |
| Nup60 | 389–539 | 0.539 | 151* |

Supplementary Table 2 - Overview of Saccharomyces Cerevisiae FG-Nup domains. Domain definitions (*e.g.* collapsed or extended) follow those outlined by Yamada et al^1^. The FG-motif spacing (in number of residues) is calculated by determining the average amino acid spacing between any FG or GLFG-motifs. The charged-to-hydrophobic amino acid ratio (C/H) was calculated using Equation 1 and the normalized hydrophobicity values in Supplementary Table 4. The domains marked with an asterisk (*) comprise a high charge content and do not contain any ‘FG’ or ‘GLFG’ motifs, hence the spacing coincides with the full length of the domain.

| System | System size (no. of beads) | Number of simulations | Forcefield | Timestep (ps) | Simulation time (steps) |
| --- | --- | --- | --- | --- | --- |
| Single FG-spacing or C/H ratio variants | 803 | 1400 (50 per d_FG_,C/H) | 1-BPA | 0.02 | 1.5 μs (7.5·10^7^) |
| FG-Nup-coated nanopores in absence of cargo | 102731 | 21 | 1-BPA-CP^2^ | 0.02 | 5 μs (2.5·10^8^) |
| FG-Nup-coated nanopores in absence of cargo | 102731 | 21 | 1-BPA-CP^2^ | 0.0175 | 5.0 μs (2.86·10^8^) |
| FG-Nup-coated nanopores in presence of cargo, equilibration | 142841 | 18 | 1-BPA-CP^2^ | 0.02 | 500ns (2.5·10^7^) |
| FG-Nup coated nanopores in presence of cargo | 142841 | 18 | 1-BPA-CP^2^ | 0.02/0.0175 | 5.0-12μs  (up to 6·10^8^) |
| Binding simulations of FG spacing or C/H ratio variants with Kap95 | 1663 | 20 per FG-spacing and C/H (420 in total) | 1-BPA-CP^2^ | 0.02 | Up to 300 μs (1.5·10^10^) |

### Supplementary Table 3 – Overview of simulations in this work.

| **AA** | A | R* | N | D* | C | Q | E* | G | H | I |
| --- | --- | --- | --- | --- | --- | --- | --- | --- | --- | --- |
| $\boldsymbol{\varepsilon}_{\mathbf{i}}$ | 0.7 | 0.005 | 0.33 | 0.005 | 0.68 | 0.64 | 0.005 | 0.41 | 0.53 | 0.98 |
| **AA** | L | K* | M | F | P | S | T | W | Y | V |
| $\boldsymbol{\varepsilon}_{\mathbf{i}}$ | 1 | 0.005 | 0.78 | 1 | 0.65 | 0.45 | 0.51 | 0.96 | 0.82 | 0.94 |

Supplementary Table 4 – Hydrophobicity scale used in our 1-BPA model. Hydrophobicity-values of charged residues have been increased with respect to the original version^3^. This hydrophobicity scale is used to calculate the C/H-ratio values.

| **Particle type** | **Consists of:** | $\boldsymbol{\sigma}$ **(nm)** |
| --- | --- | --- |
| AA | All residues not involved in cation-pi (RK/FYW) or binding site (B-RK, B-FYW, F1, G1) interactions | 0.6 |
| RK | R or K-residues that are not part of a Kap95 binding site | 0.6 |
| FYW | F, Y or W residues that are not part of a Kap95 binding site or FG-motif (in case of F). | 0.6 (hydrophobic) or 0.45 (cation-pi) |
| B-RK | R or K-residues that are part of a Kap95 binding site | 0.6 or 0.45 |
| B-FYW | F, Y or W residues that are part of a Kap95 binding site | 0.6 or 0.45 |
| F1 | F-residue within an FG-motif | 0.6 |
| G1 | G-residue within an FG-motif | 0.6 |
| WALL | Sterically beads that constitute the nanopore scaffold | 3 |
| BOUNDS | Sterically inert beads that confine only Kap95 to a cylindrical region on either side of the nanopore membrane | 3 |

Supplementary Table 5 – Overview of interaction types.

|  | **62.5 nM Kap95** | | | | **500 nM Kap95** | | | |
| --- | --- | --- | --- | --- | --- | --- | --- | --- |
|  | $\boldsymbol{A}$**(Hz)** | $\boldsymbol{B}$**(Hz)** | $\boldsymbol{\tau}_{\boldsymbol{1}}$**(s)** | $\boldsymbol{\tau}_{\boldsymbol{2}}$**(s)** | $\boldsymbol{A}$**(Hz)** | $\boldsymbol{B}$**(Hz)** | $\boldsymbol{\tau}_{\boldsymbol{1}}$**(s)** | $\boldsymbol{\tau}_{\boldsymbol{2}}$**(s)** |
| **NupY(13-0.0478)^1^** | -13.23 | -4.823 | 32.88 | 2.577 | -2.394 | -16.21 | 1.213 | 21.76 |
| **NupY(52-0.0478)** | -3.904 | -3.145 | 0.9054 | 8.391 | -0.5391 | -1.611 | 0.7623 | 9.389 |
| **NupY(104-0.0478)** | -4.265 | -1.707 | 0.7191 | 7.46 | -0.04891 | -0.635 | 0.009876 | 0.4711 |
| **NupY(104-0.5736)** | -2.791 | -5.727 | 2.162 | 14.97 | -9.784 | -1.998 | 15.35 | 1.215 |
| **NupY(13-0.0478)^2^** | -2.764 | -11.96 | 0.863 | 18.73 | -9.252 | -21.73 | 11.12 | 36.28 |
| **NupY(13-0.0956)** | -2.758 | -8.144 | 0.7711 | 18.29 | -0.2957 | -20.04 | 0.1647 | 19.78 |
| **NupY(13-0.1912)** | -9.502 | -10.62 | 1.813 | 52.19 | -2.683 | -14.14 | 0.4817 | 20.73 |
| **NupY(13-0.3824)** | -6.146 | -11.78 | 1.923 | 12.99 | -13.18 | -1.483 | 17.66 | 1.061 |

### Supplementary Table 6 – Extracted fit parameters from fitting frequency shifts.

The reported parameters $A$, $B$, $\tau_{1}$, $\tau_{2}$ resulted from fitting the function $f\left( t \right)=Ae^{\frac{t}{\tau_{1}}}+Be^{\frac{t}{\tau_{2}}}$ to the frequency shifts shown in shown in Figures 5l,m and 6j,k. The start of the fitted curves was zeroed before applying fitting to reduce the number of free parameters. ^1,2^: Parameters for Kap95 binding to NupY_13-0.0478_ in the two comparisons shown in Figure 5 and 6, respectively.

| **Protein** | **Sequence** |
| --- | --- |
| **Kap95** | MSTAEFAQLLENSILSPDQNIRLTSETQLKKLSNDNFLQFAGLSSQVLIDENTKLEGRILAALTLKNELVSKDSVKTQQFAQRWITQVSPEAKNQIKTNALTALVSIEPRIANAAAQLIAAIADIELPHGAWPELMKIMVDNTGAEQPENVKRASLLALGYMCESADPQSQALVSSSNNILIAIVQGAQSTETSKAVRLAALNALADSLIFIKNNMEREGERNYLMQVVCEATQAEDIEVQAAAFGCLCKIMSLYYTFMKPYMEQALYALTIATMKSPNDKVASMTVEFWSTICEEEIDIAYELAQFPQSPLQSYNFALSSIKDVVPNLLNLLTRQNEDPEDDDWNVSMSAGACLQLFAQNCGNHILEPVLEFVEQNITADNWRNREAAVMAFGSIMDGPDKVQRTYYVHQALPSILNLMNDQSLQVKETTAWCIGRIADSVAESIDPQQHLPGVVQACLIGLQDHPKVATNCSWTIINLVEQLAEATPSPIYNFYPALVDGLIGAANRIDNEFNARASAFSALTTMVEYATDTVAETSASISTFVMDKLGQTMSVDENQLTLEDAQSLQELQSNILTVLAAVIRKSPSSVEPVADMLMGLFFRLLEKKDSAFIEDDVFYAISALAASLGKGFEKYLETFSPYLLKALNQVDSPVSITAVGFIADISNSLEEDFRRYSDAMMNVLAQMISNPNARRELKPAVLSVFGDIASNIGADFIPYLNDIMALCVAAQNTKPENGTLEALDYQIKVLEAVLDAYVGIVAGLHDKPEALFPYVGTIFQFIAQVAEDPQLYSEDATSRAAVGLIGDIAAMFPDGSIKQFYGQDWVIDYIKRTRSGQLFSQATKDTARWAREQQKRQLSLLPETGG |
| **IBB-GFP** | MSKHHHHSGHHHTGHHHHSGSHHHTGENLYFQGSDNGTDSSTSKFVPEYRRTNFKNKGRFSADELRRRRDTQQVELRKAKRDEALAKRRNFIPPTDGTSKGEELFTGVVPILVELDGDVNGHKFSVSGEGEGDATYGKLTLKFICTTGKLPVPWPTLVTTLTYGVQCFSRYPDHMKQHDFFKSAMPEGYVQERTIFFKDDGNYKTRAEVKFEGDTLVNRIELKGIDFKEDGNILGHKLEYNYNSHNVYIMADKQKNGIKVNFKIRHNIEDGSVQLADHYQQNTPIGDGPVLLPDNHYLSTQSALSKDPNEKRDHMVLKEFVTAAGITLGMDELYKTGC |
| **2xMBP-IBB** | MKIEEGKLVIWINGDKGYNGLAEVGKKFEKDTGIKVTVEHPDKLEEKFPQVAATGDGPDIIFWAHDRFGGYAQSGLLAEITPDKAFQDKLYPFTWDAVRYNGKLIAYPIAVEALSLIYNKDLLPNPPKTWEEIPALDKELKAKGKSALMFNLQEPYFTWPLIAADGGYAFKYENGKYDIKDVGVDNAGAKAGLTFLVDLIKNKHMNADTDYSIAEAAFNKGETAMTINGPWAWSNIDTSKVNYGVTVLPTFKGQPSKPFVGVLSAGINAASPNKELAKEFLENYLLTDEGLEAVNKDKPLGAVALKSYEEELVKDPRIAATMENAQKGEIMPNIPQMSAFWYAVRTAVINAASGRQTVDEALKDAQTNSSSNNNNNNNNNNLGIEGRISHMLEVLFQGPMKIEEGKLVIWINGDKGYNGLAEVGKKFEKDTGIKVTVEHPDKLEEKFPQVAATGDGPDIIFWAHDRFGGYAQSGLLAEITPDKAFQDKLYPFTWDAVRYNGKLIAYPIAVEALSLIYNKDLLPNPPKTWEEIPALDKELKAKGKSALMFNLQEPYFTWPLIAADGGYAFKYENGKYDIKDVGVDNAGAKAGLTFLVDLIKNKHMNADTDYSIAEAAFNKGETAMTINGPWAWSNIDTSKVNYGVTVLPTFKGQPSKPFVGVLSAGINAASPNKELAKEFLENYLLTDEGLEAVNKDKPLGAVALKSYEEELVKDPRIAATMENAQKGEIMPNIPQMSAFWYAVRTAVINAASGRQTVDEALKDAQTCCGPCCDNGTDSSTSKFVPEYRRTNFKNKGRFSADELRRRRDTQQVELRKAKRDEALAKRRNFIPPTDHHHHHHHH |

Supplementary Table 7 – Amino acid sequences of Kap95 and cargoes used in the study.

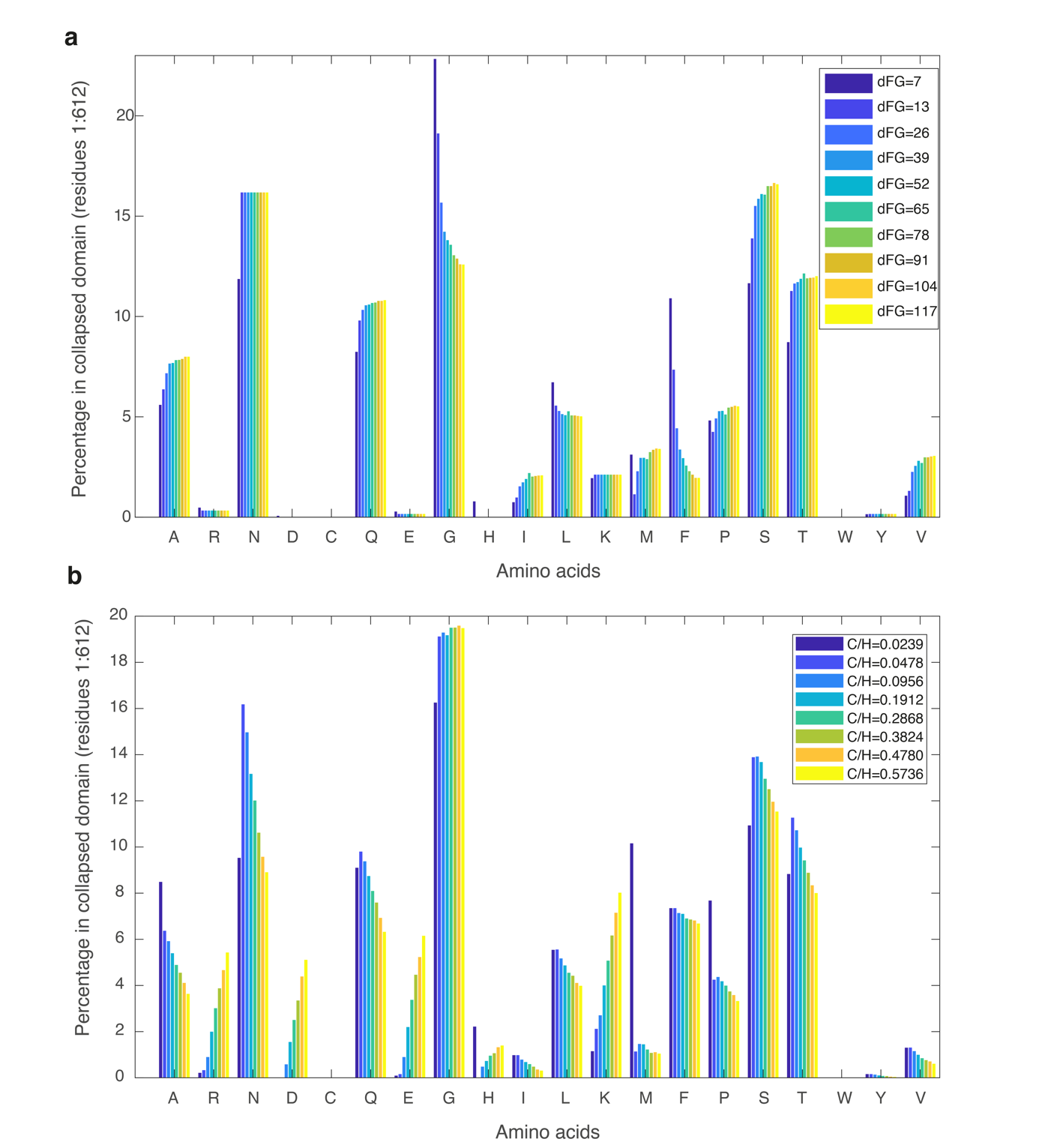

Supplementary Figure 1 – Sequence histograms for FG-spacing variants and C/H-ratio variants. **a:** The average occurrence of different amino acids in the 50 highest-ranking designs for FG-spacing variants (with fixed C/H-ratio at the native GLFG-Nup average of 0.0478). Increasing the FG-spacing leads to an exchange of FG and GLFG-motifs motifs for other isohydrophobic pairs of amino acids (AA, MQ, MP, IG, LF, IS, LS, VS, VT). **b:** The average occurrence of different amino acids in the 50 highest-ranking designs for C/H-ratio variants (with fixed FG-spacing at the native GLFG-Nup average of 13). To achieve higher C/H-ratios, additional charge content (D,E,K,R) is introduced that replaces polar uncharged residues (N,Q,T,S). To reduce the hydrophobicity below the native GLFG-Nup average of 0.0478, additional hydrophobic residues were introduced that do not strongly enhance secondary structure (*e.g.*, A, M, or P).

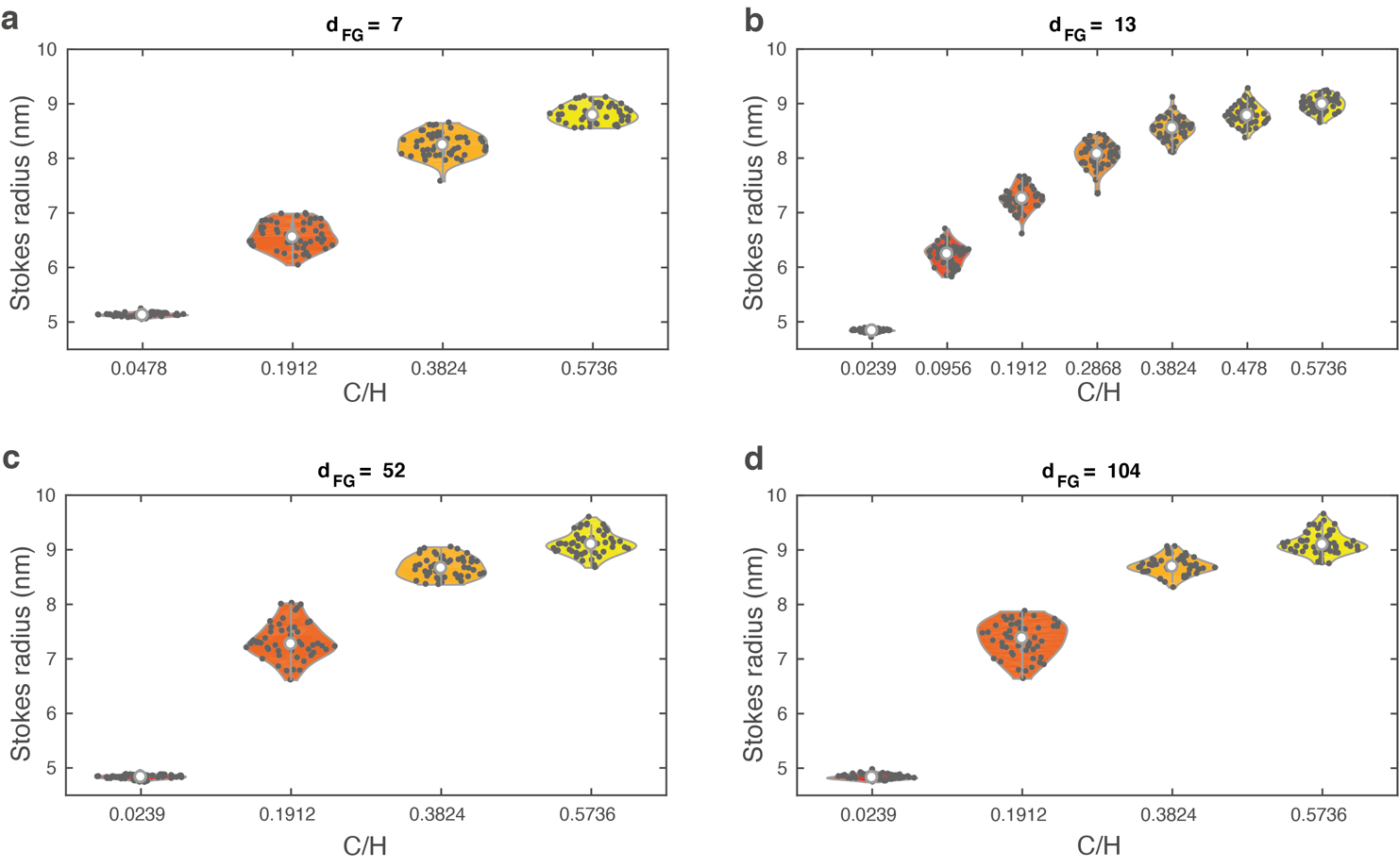

Supplementary Figure 2 – Violin plots of the Stokes radii of 50 different designs for different combinations of C/H-ratio and FG-spacing. Compared to the Stokes radii for designs comprising C/H and d_FG_-values close to the GLFG-Nup average (*i.e.,* panel (b) is reproduced from Figure 1g in the main text), we observe similar trends for other combinations (a,c,d). Namely, increases in C/H lead to a monotonic increase in FG-Nup dimensions, the polymer dimensions are similar between C/H-variants with the same d_FG_, and the variability in Stokes radii between designs is small (~10%).

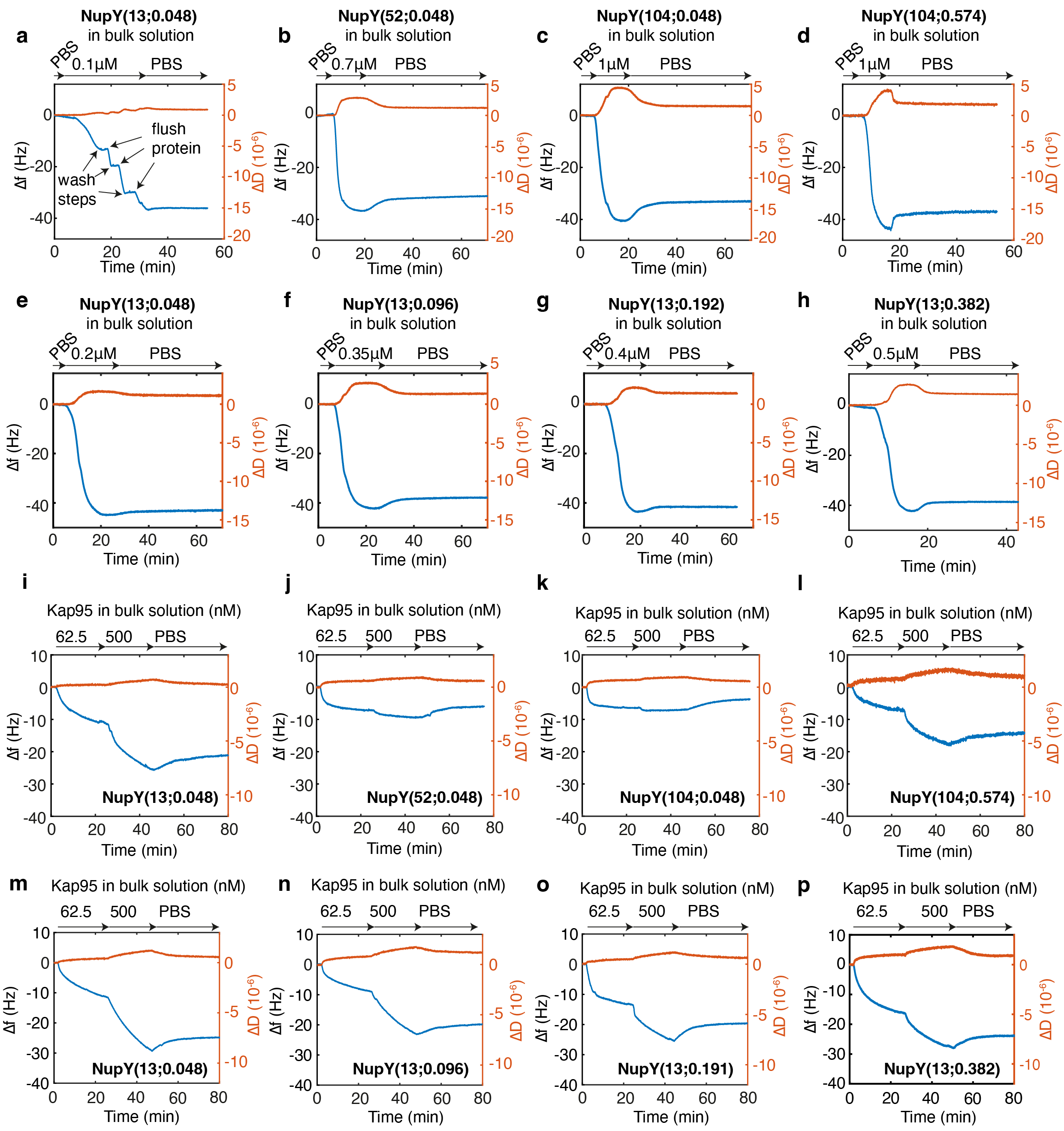

### Supplementary Figure 3 – Overview of QCM-D experiments on NupY brushes.

**a-h)** Coating of the QCM-D sensor with the indicated NupY variants. The concentration was individually adjusted to achieve similar frequency response Δf for all variants. **i-p)** Addition of Kap95 to the pre-formed NupY brushes at concentrations of 62.5 and 500 nM shows different degree of interaction for the tested NupY variants.

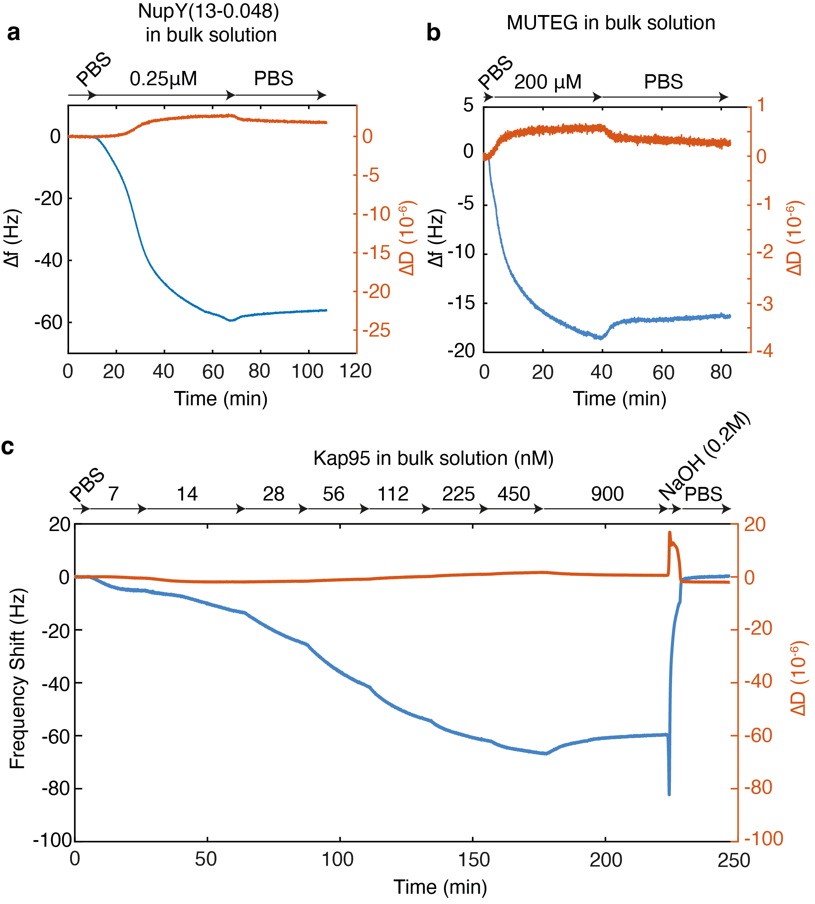

### Supplementary Figure 4 – MUTEG coating and NaOH dissociation of NupY-bound Kap95 in QCM-D experiments.

**a)** Coating of a sensor with our template NupY(13;0.05) at 0.25 μM concentration. **b)** Passivating with MUTEG at 200 μM of the remaining exposed gold surface following NupY coating. **c)** Titration of Kap95 from concentrations 7–900 nM showing efficient binding to the NupY(13;0.05) brush followed by complete dissociation by 0.2 M NaOH.

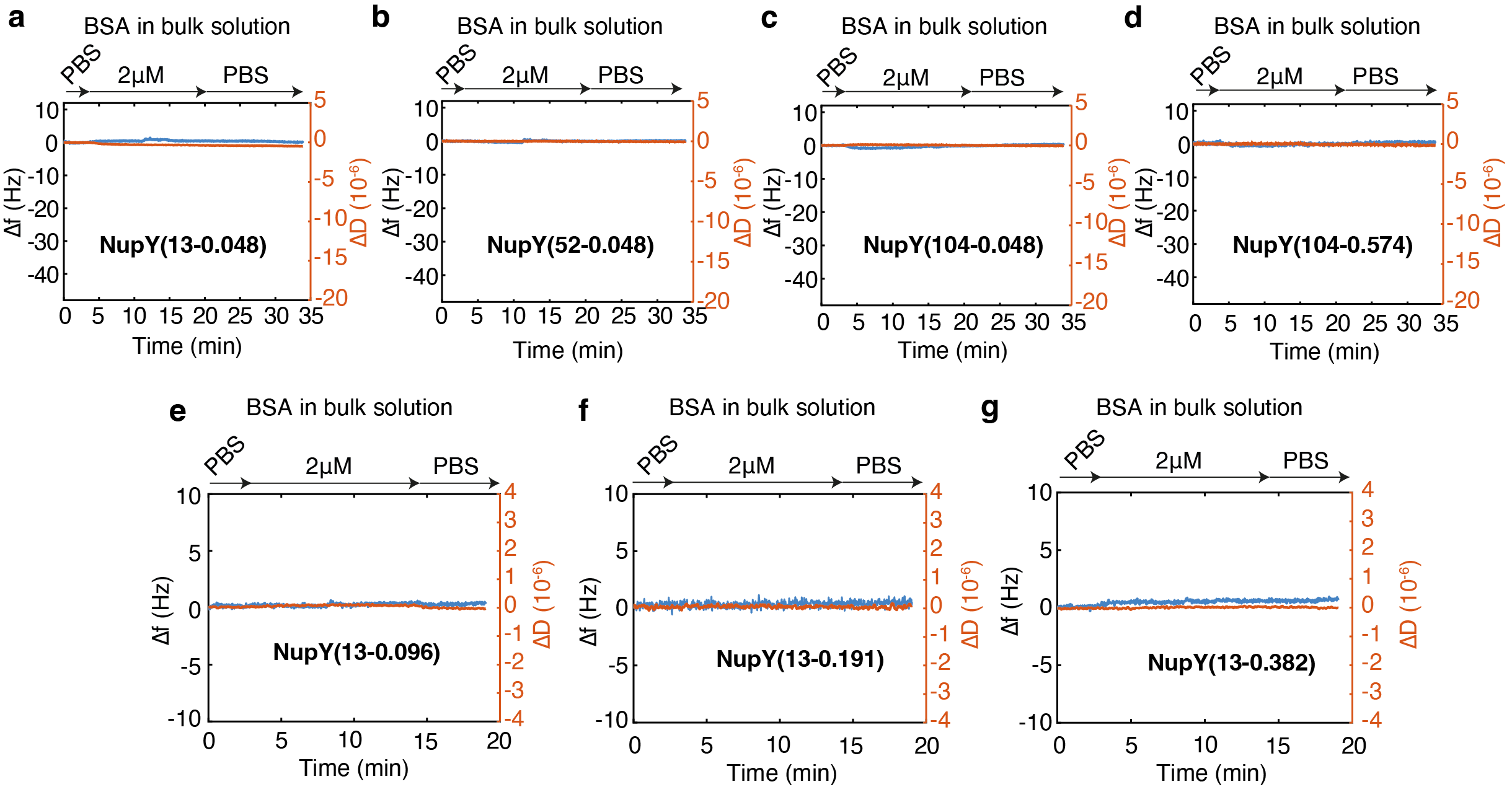

### Supplementary Figure 5 – Response of NupY-coated QCM-D chips to BSA.

Upon flushing high concentration (2 μM) of BSA onto the NupY-coated sensors no significant change in the resonance frequency nor dissipation was detected, indicating a complete lack of interaction between all NupY variants and BSA.

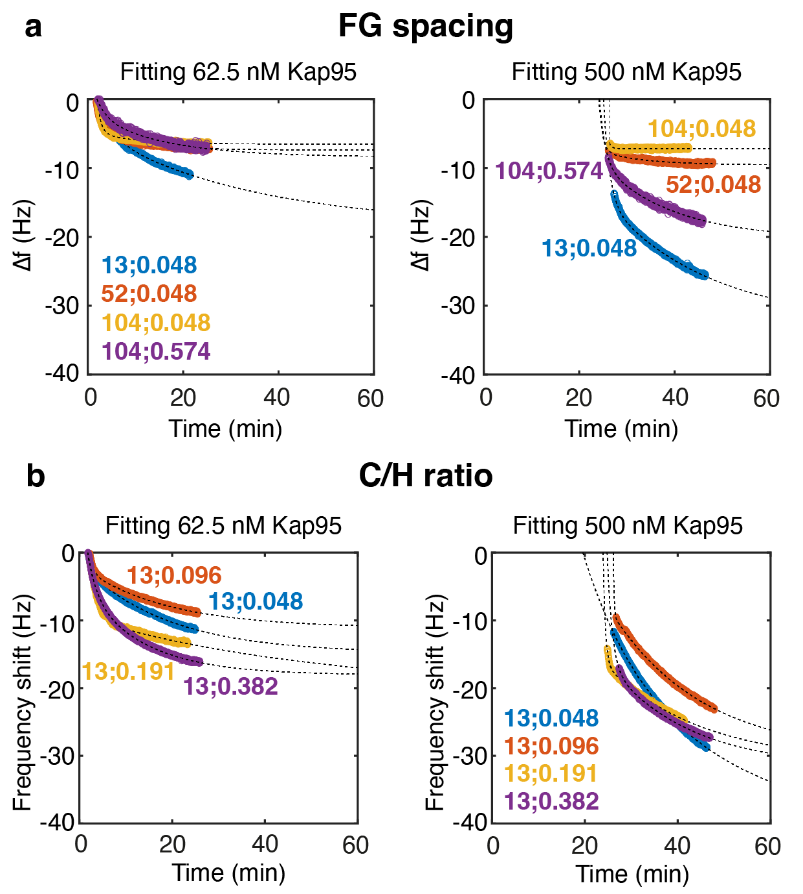

### Supplementary Figure 6 – Fitting of the QCM-D frequency response upon Kap95 binding.

The frequency shift Δf for the FG-spacing (**a**) and C/H variants (**b**) after sequential flushing of 62.5 and 500 nM Kap95 was fitted to a bi-exponential model function to extract the half-time t_1/2_ and saturation value Δf_max_.

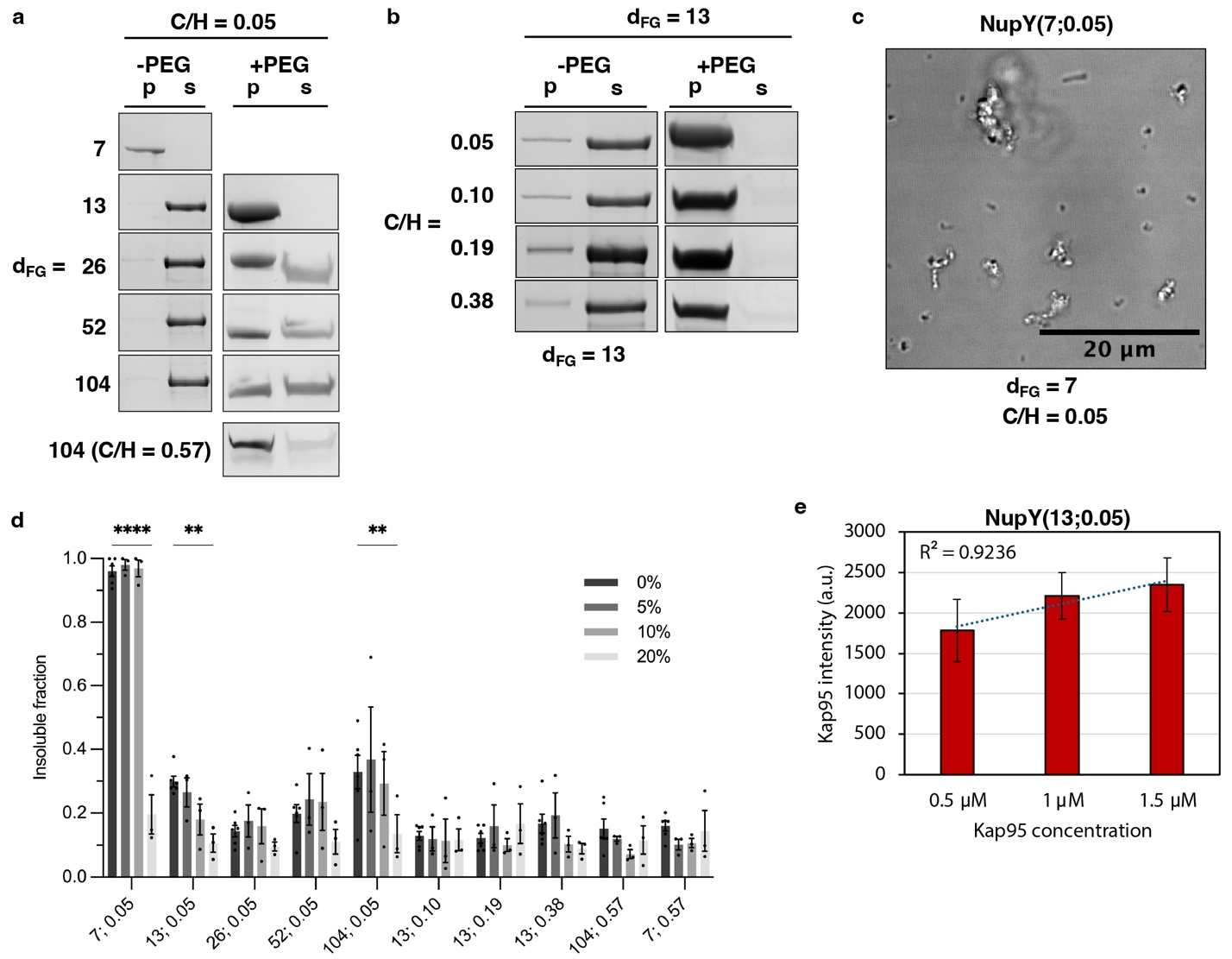

Supplementary Figure 7 – Characterization of the liquid-liquid phase separation propensity and aggregation of the different NupY variants.

**a-b)** The degree of condensation at a concentration of 200 nM in the absence or presence of 10% w/v PEG-8000 in 50 mM Tris, 150 mM NaCl, pH 7.5 is assessed by a sedimentation assay. p: pellet, s: supernatant. **c)** The variant NupY(7;0.05) formed amorphous aggregates at 200 nM in the absence of PEG. **d)** Quantification of the insoluble fraction of NupY condensates formed at room temperature after 1 hour (3μM protein in 50 mM Tris–HCl pH 8.0, 150 mM NaCl, and indicated concentrations of 1,6-hexanediol). Graph shows mean ± SEM (n=3-6). Two-way ANOVA with Dunnett's multiple comparisons test compared to 0% 1,6-hexanediol for each NupY variant. **** p < 0.0001, ** p <0.01 e) Saturation of condensates formed by the variant NupY(13;0.05) under increasing concentration of Kap95. The marginal increase from 1 µM to 1.5 µM indicates that condensates are close to reaching saturation under the experimental conditions.

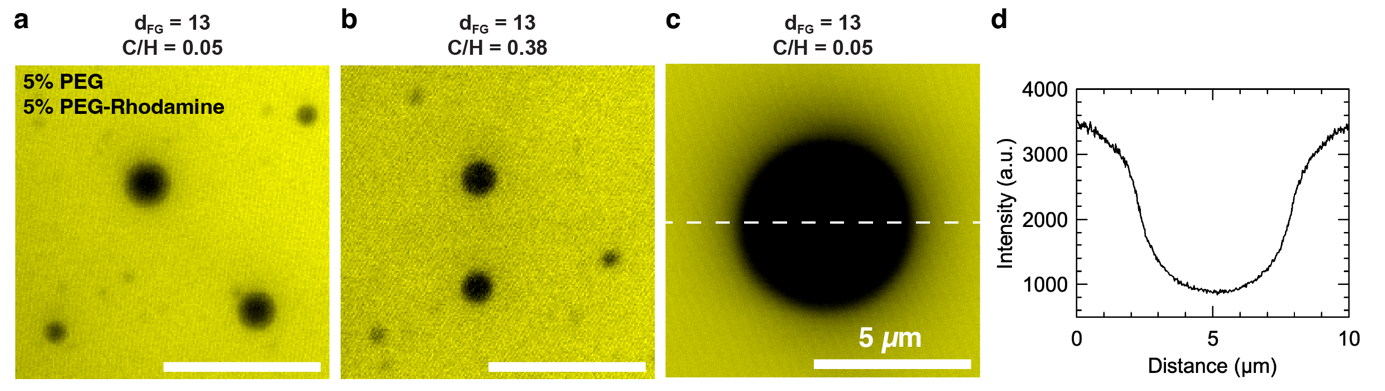

Supplementary Figure 8 – PEG does not partition into NupY condensates. Condensates of the template NupY(13;0.05) and variant NupY(13;0.38) were formed in 5% w/v PEG-8000 and 5% w/v PEG-8000-Rhodamine in 50 mM Tris, 150 mM NaCl, pH 7.5. PEG is excluded from the condensates. a-c) Confocal fluorescence images of NupY condensates. Scalebar in a-b: 10 µm. d) Line profile of the PEG-Rhodamine intensity as indicated in c.

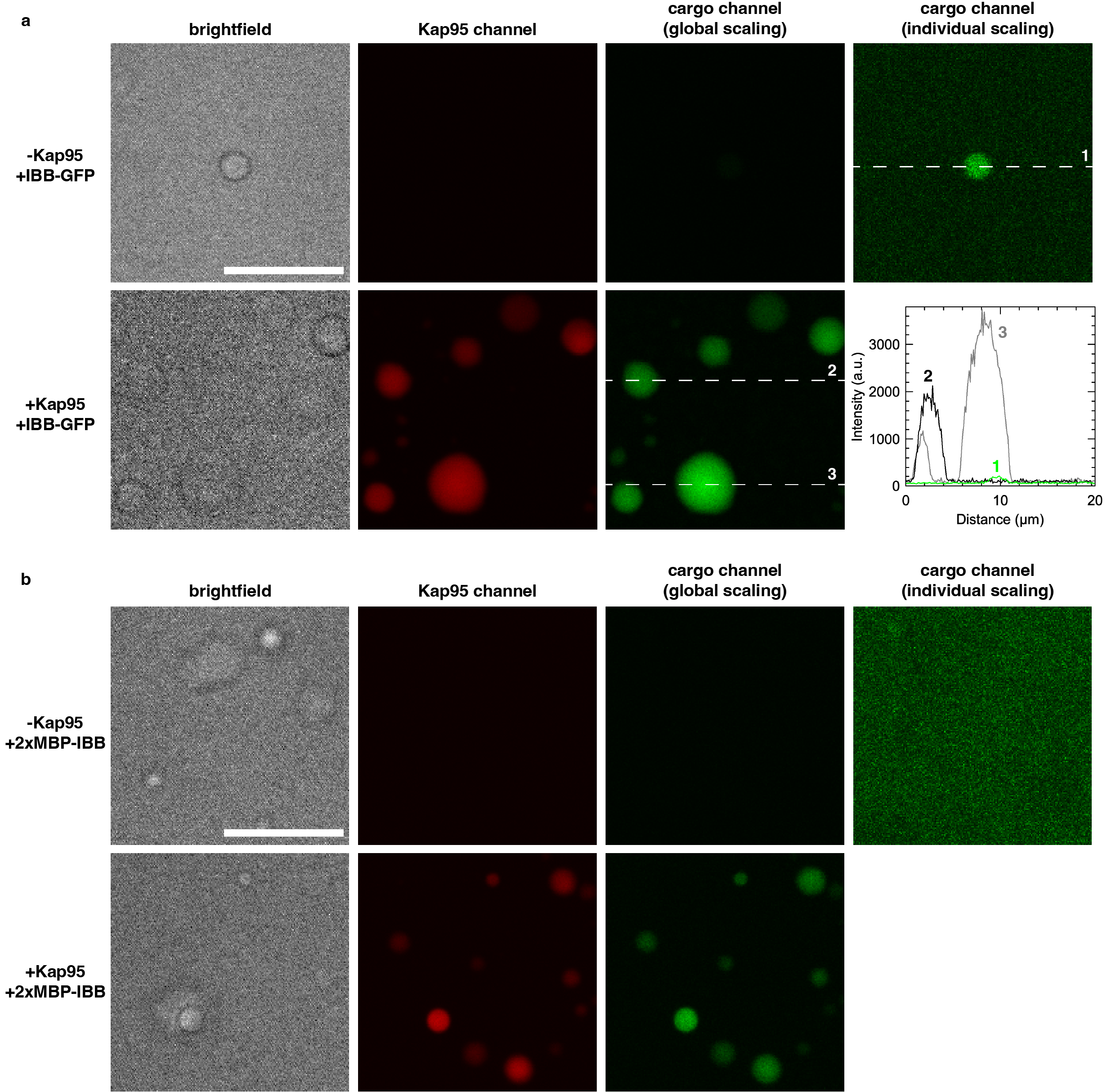

Supplementary Figure 9 – Minimal interactions of model cargoes with NupY condensates. Condensates of the template NupY(13;0.05) were formed in 10% w/v PEG-8000 in 50 mM Tris, 150 mM NaCl, pH 7.5, and supplemented by 1 µM cargo (**a**: IBB-GFP, **b**: 2xMBP-IBB) in the absence (top) or presence (bottom) of 1 µM Kap95. Scalebars: 10 µm. The cargo channel in the presence and absence of Kap95 is shown in global scaling between the two conditions (third column) and individually scaled (fourth column). A minimal amount of interaction is detected for the IBB-GFP cargo (**a**, right) which is negligible compared to the signal in the presence of Kap95 (see line profiles). No significant interaction is detected for the large cargo 2xMBP-IBB.

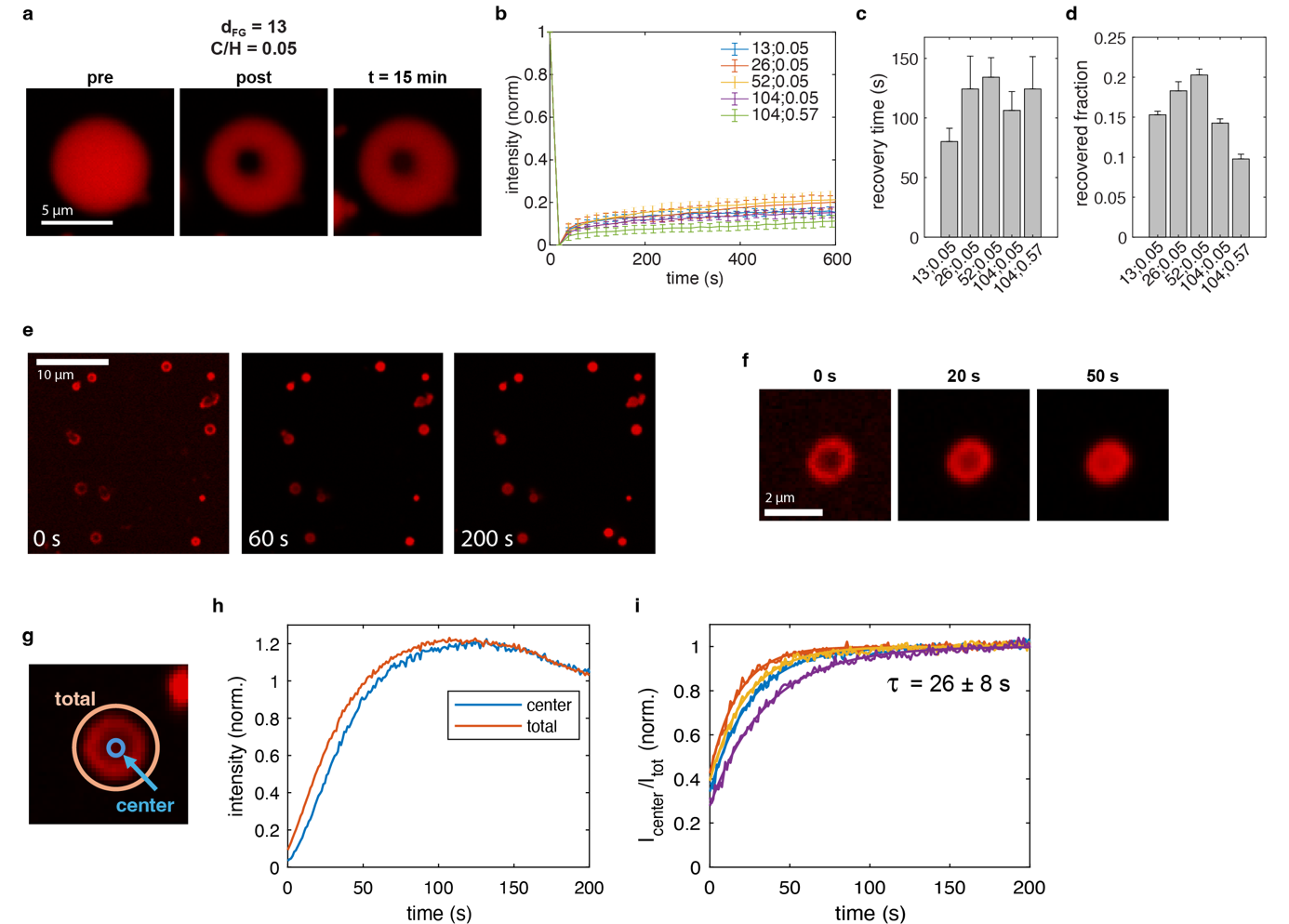

Supplementary Figure 10 – Diffusion kinetics of Kap95 in NupY condensates. Condensates of the different NupY variants were formed at 200 nM in 10% w/v PEG-8000 in 50 mM Tris, 150 mM NaCl, pH 7.5. **a)** FRAP experiment of a condensate formed by the template NupY loaded with 1 µM Kap95. Minimal recovery occurs over the timescale of 15 min. **b)** Recovery curves of the different NupY variants showing low recovery over 10 min. **c-d)** Recovery time and recovered fraction of the data shown in b. **e)** An influx experiment where Kap95 was added to fresh condensates of the template NupY_13-0.05_ at a concentration of 1 µM. The contrast of the images was individually adjusted. **f)** Zoom-in of an exemplary particle from e. **g)** The influx of Kap95 was quantified by comparing the the total intensity to the intensity in the center of the condensate. **h)** Exemplary quantification of the influx as shown in h. Curves were each normalized to their respective signal at the end of the experiment. **i)** Comparison of the influx curves of four different particles. Each curve was normalized to the intensity ratio at the end of the experiment. Kap95 influx occurs on the timescale of 26±8 s.

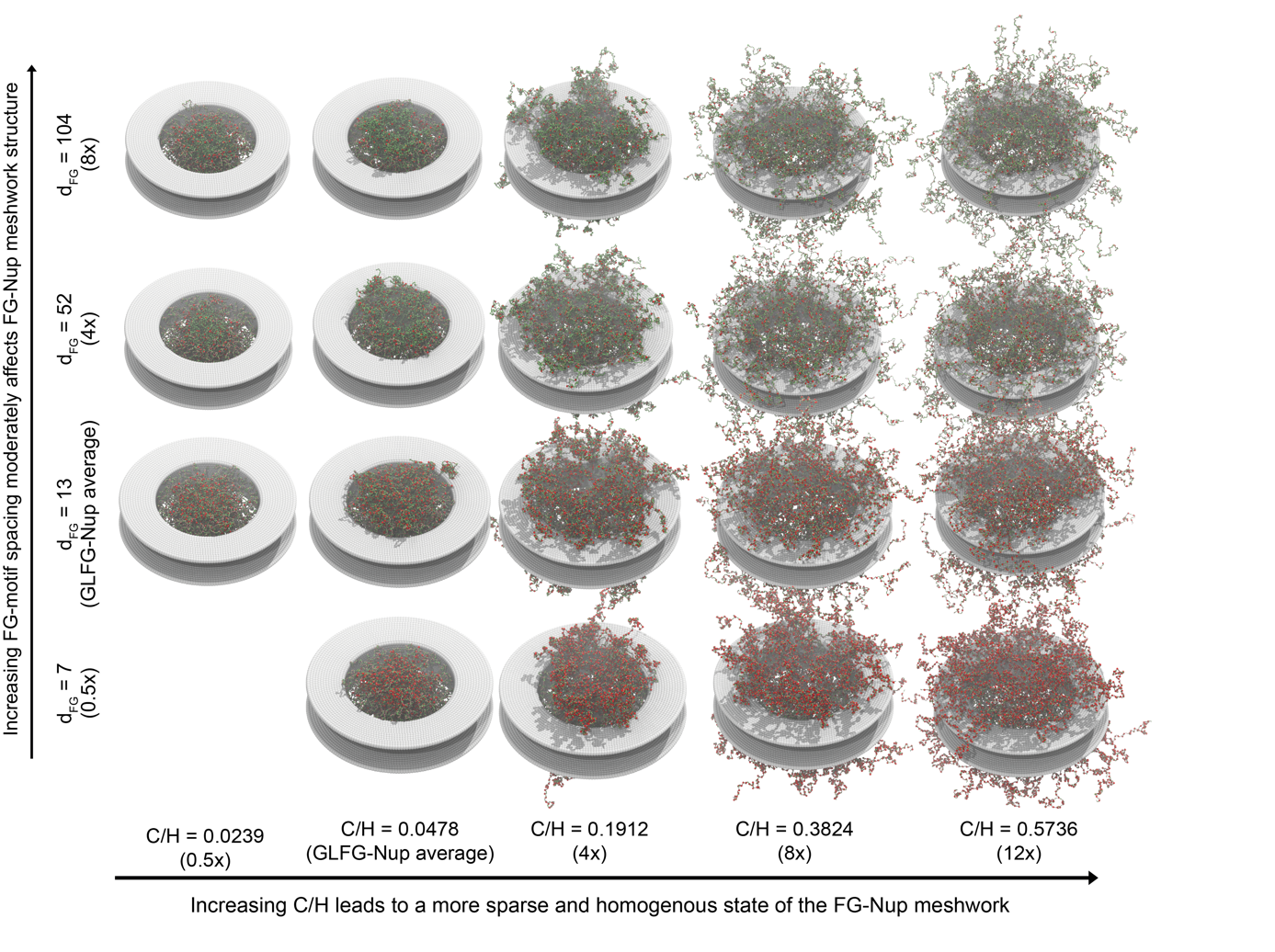
Supplementary Figure 11 – Snapshots of nanopores coated with FG-spacing and C/H-variants. Generally, the distribution of FG-Nups (green) and FG-motifs (red) is strongly affected by the value of C/H (horizontal axis). For low C/H-values, our FG-Nups seal the pore interior, where the collapsed domains (residues 1-610) form dense regions near the pore center. At larger values of C/H (beyond 4x the native average or 0.1912) a homogenous and sparse meshwork of FG-Nups forms, extending outwards from the pore region. The distribution of FG-Nups is less strongly affected by the FG-spacing (vertical axis): in general, the distribution of FG-Nups remains similar for FG-spacing variants, as the patterning of hydrophobic and C/H is conserved. The 0.5x-spacing (d_FG_=7)-variants provide an exception to this rule, as the enhanced patterning of hydrophobic residues leads to an additional collapse of the FG-Nup meshwork for all variants.

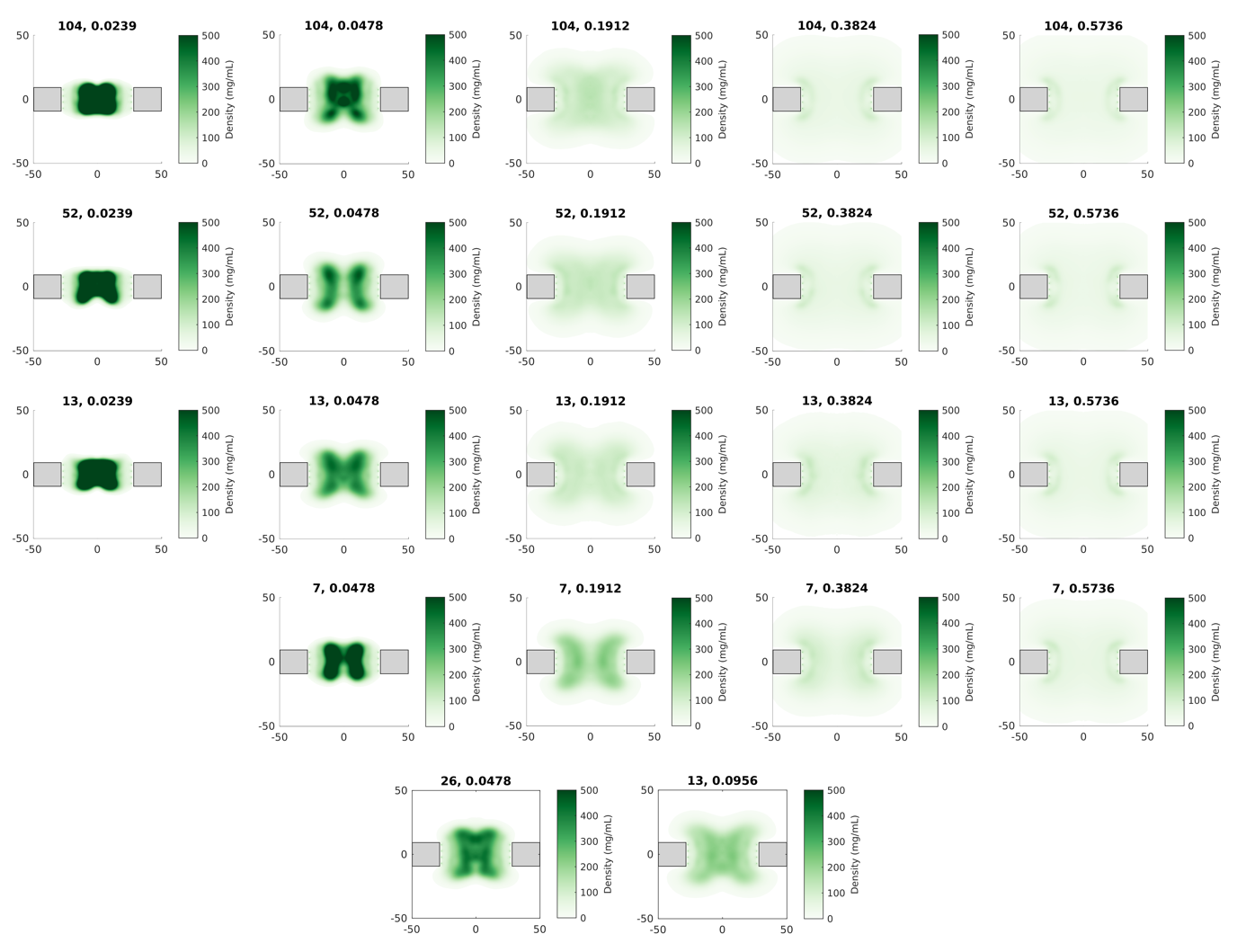

Supplementary Figure 12 – Time-averaged, axi-radial density graphs of FG-Nups in nanopores coated with artificial FG-Nups, grouped by C/H and FG-spacing. Scaffolds are highlighted in gray. Additional variants with 2xFG-spacing and 2xC/H are shown separately on the bottom row. The value of C/H (horizontal direction) strongly affects the distribution of FG-Nups in the nanopore lumen: for the native GLFG-Nup average (0.0478), a dense, ring-like or ‘doughnut-like’ structure is formed by the FG-Nups, with densities exceeding 500 mg/mL. A further reduction in C/H (0.0239) leads to a strongly collapsed meshwork with densities exceeding 1000 mg/mL. Increases (C/H=0.0956 and higher) lead to the formation of a more homogenous meshwork, where the density inside the pore lumen steeply drops, from 300 mg/mL to ~100 mg/mL. Increases in FG-spacing (vertical direction) w.r.t. the native GLFG-Nup average (dF_G_=13) do not notably affect the distribution of the FG-Nups, as the patterning of hydrophobic residues and overall C/H are conserved. For the variants with a decreased spacing (d_FG_=7, top row), we find that regardless of conserved C/H, the increased number and degree of patterning of hydrophobic residues leads to an additional collapse of the FG-Nup mesh.

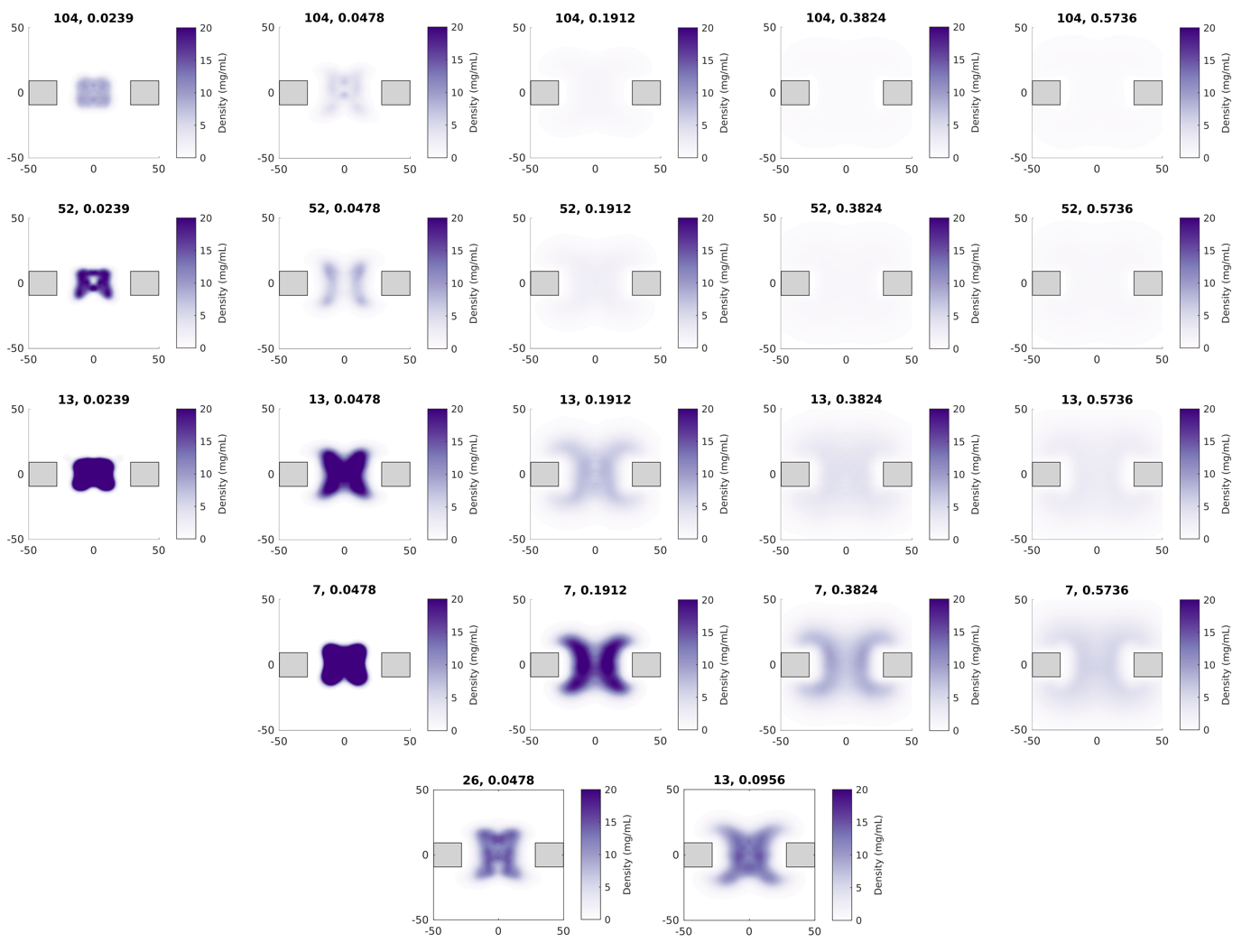

Supplementary Figure 13 – Time-averaged, axi-radial density graphs of F(G)-motifs in nanopores coated with artificial FG-Nups, grouped by C/H and FG-spacing. Scaffolds are highlighted in gray. Additional variants with 2xFG-spacing and 2xC/H are shown separately on the bottom row. The FG-motif distribution coincides with the distribution of the NupY-domains that are varied in this study (residues 1-610). Changes in the localization of this domain by virtue of changing C/H (horizontal direction) similarly affect the localization of the FG-motifs, where a dense, ring-like or ‘doughnut-like’ central region is formed that comprises all the FG-motifs for low values of C/H, and the FG-motifs are spread out rather evenly for higher C/H-values, albeit with a significantly reduced density. Changes in the FG-spacing (vertical direction), which occur under similar C/H, generally lead to a preserved shape of the FG-motif distribution. At the same time, the density of the FG-motifs in the nanopore scales with the change in d_FG_: compared d_FG_=13, variants with spacings of 26, 52 and 104 residues display densities that are approximately 2, 4 and 8 times lower, respectively. This trend does not hold fully for a reduced FG-spacing (d_FG_=7, top row), as the FG-Nup meshwork experiences an additional collapse by means of the enhanced patterning of hydrophobic residues.

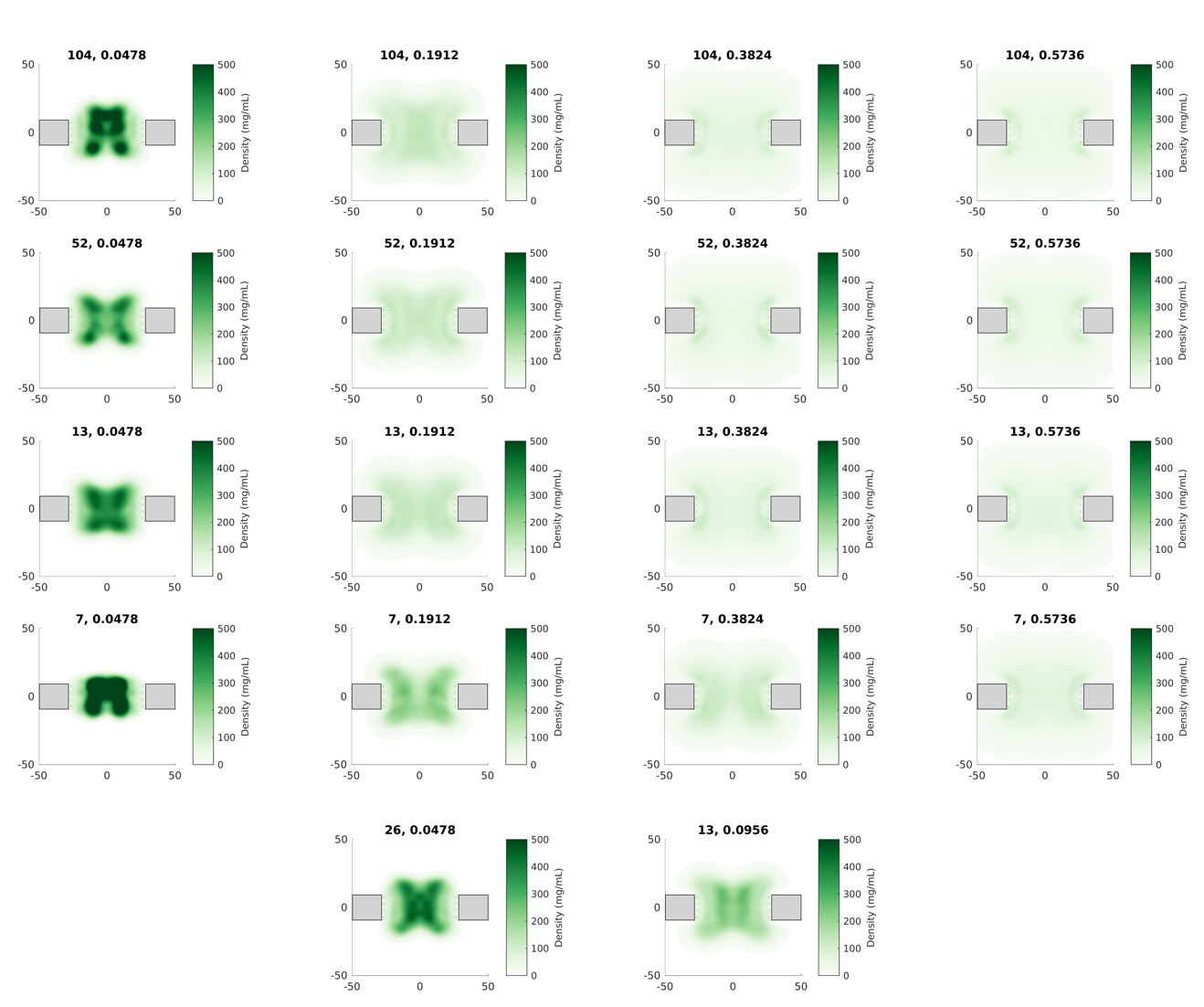

Supplementary Figure 14 – Time-averaged, axi-radial density graphs of FG-Nups in nanopores coated with artificial FG-Nups in presence of Kap95 and inert proteins, grouped by C/H and FG-spacing. Scaffolds are highlighted in gray. Additional variants with 2xFG-spacing and 2xC/H-ratio are shown separately on the bottom row. C/H=0.0239 (0.5x native average) variants were not studied in presence of cargo and NTRs due to their high FG-Nup densities. The overall FG-Nup density distributions closely follow those in Supplementary Figure 4, indicating that the presence of the studied amount of NTRs and inert molecules (10 copies of Kap95, BSA and Ubiquitin, resp.) did not appreciably alter the average structure of the FG-Nup meshwork.

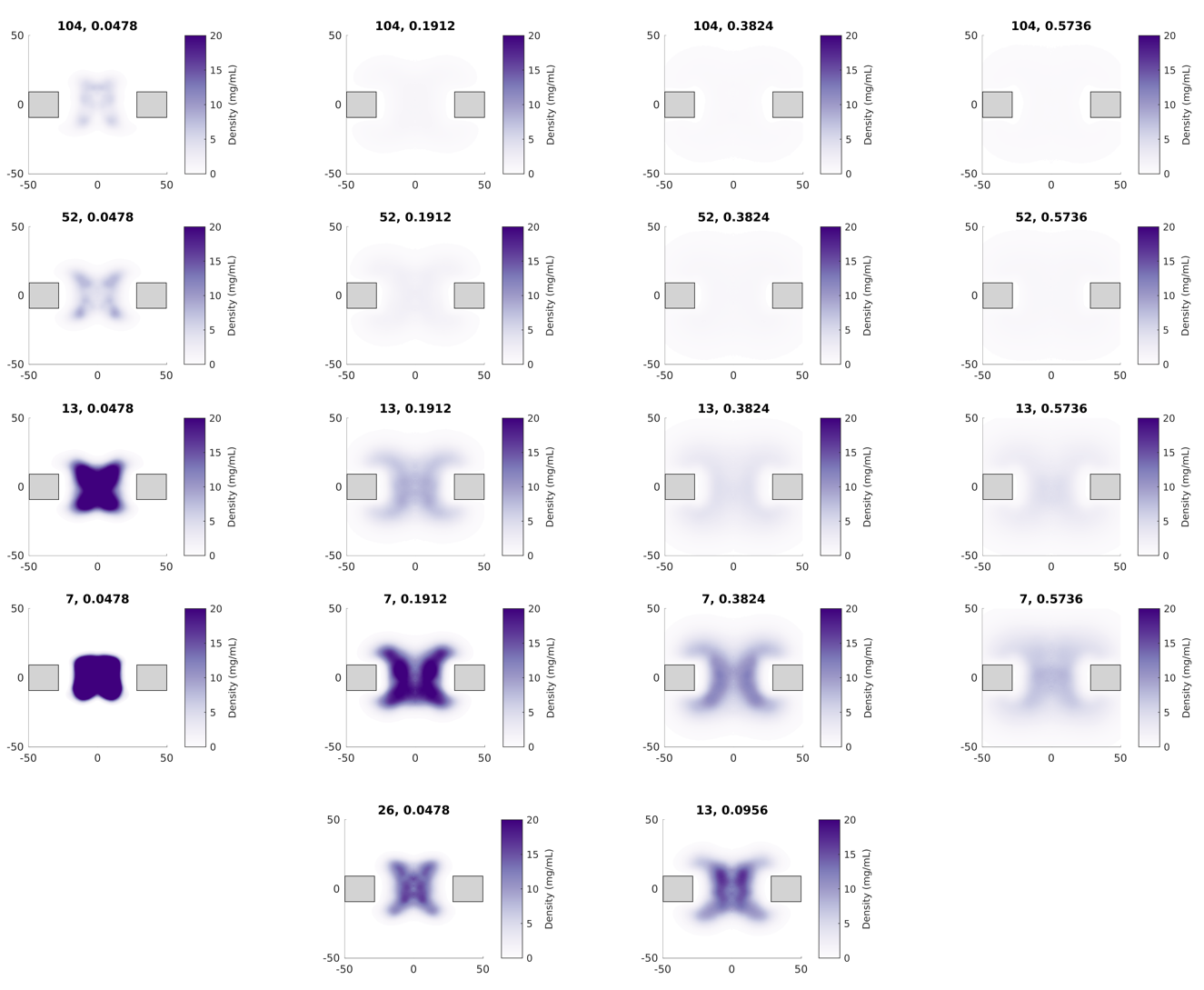

Supplementary Figure 15 – Time-averaged, axi-radial density graphs of the F(G)-motifs in nanopores coated with artificial FG-Nups in presence of Kap95 and inert proteins, grouped by C/H and FG-spacing. Scaffolds are highlighted in gray. Additional variants with 2xFG-spacing and 2xC/H-ratio are shown separately on the bottom row. C/H=0.0239 (0.5x native average) variants were not studied in presence of cargo and NTRs. Similar to the overall FG-Nup density distribution (Supplementary Figure 4, we do not find that the presence of NTRs and inert proteins has not notably affected the distribution of FG-motifs.

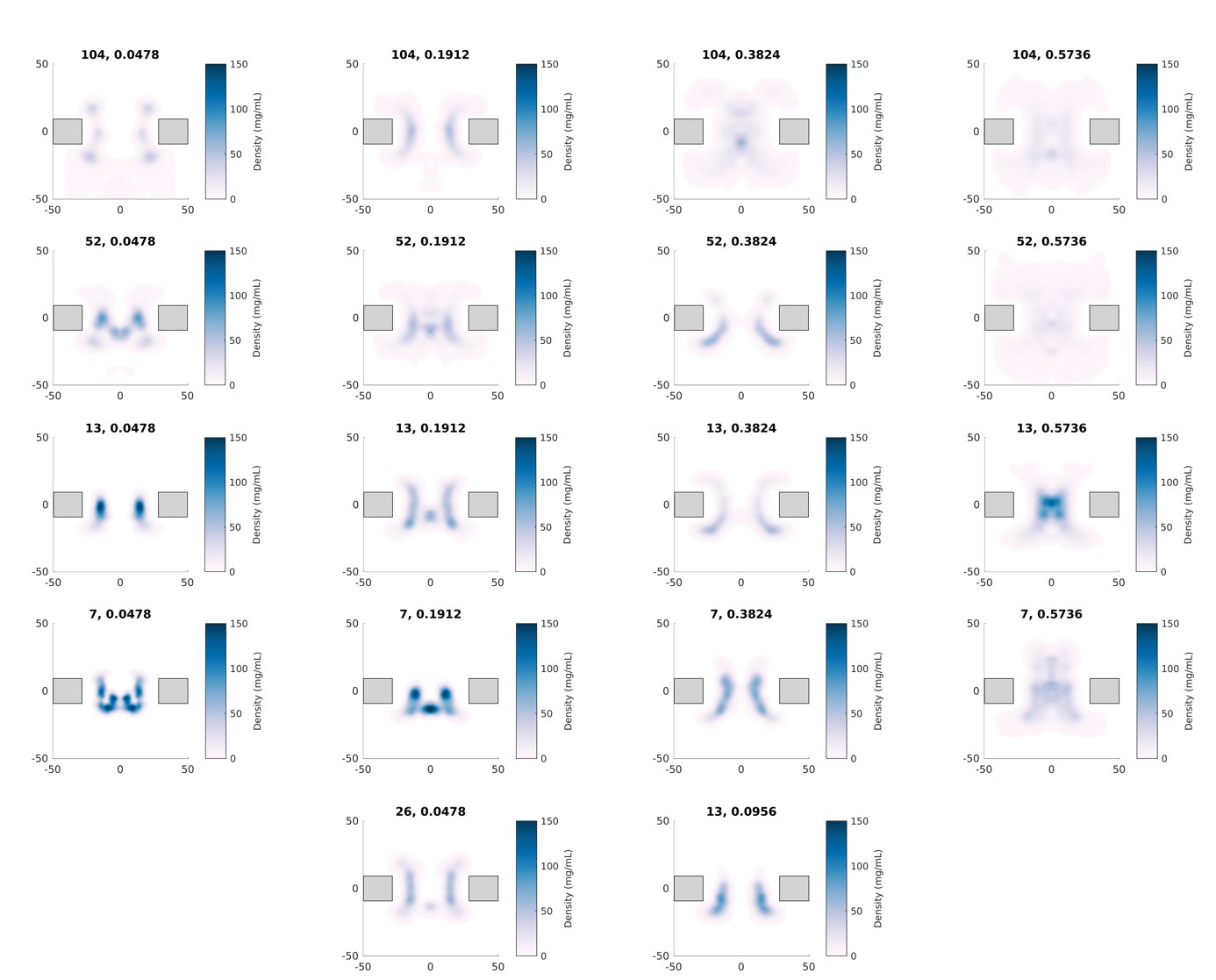

Supplementary Figure 16 – Time-averaged, axi-radial density graphs of the Kap95 protein in nanopores coated with artificial FG-Nups in presence of inert proteins, grouped by C/H-ratio and FG-spacing. Scaffolds are highlighted in gray. Additional variants with 2xFG-spacing and 2xC/H-ratio are shown separately on the bottom row. C/H=0.0239 (0.5x native average) variants were not studied in presence of cargo and NTRs. Following the changes in the FG-Nup distribution (Supplementary Figure 4 with varying C/H (horizontal direction), the distribution of Kap95 changes in a similar fashion. For low values of C/H, Kap95 interacts with FG-motifs in the dense, central bands, ring-like or ‘doughnut-like’ regions formed by the artificial FG-Nups, without fully localizing inside the dense regions. Indeed, the density distribution of Kap95 for low C/H (left column) does not fully coincide with the density distributions of FG-motifs and FG-Nups in Supplementary Figures 4-5. Rather, it is shifted radially outward. For higher values of C/H (beyond 0.0956), the FG-Nup density reduces, and the Kap95 distribution more closely follows that of the FG-motifs (more homogenous with a slight presence of two ‘bands’). Expectedly, the FG-spacing (vertical direction) affects the Kap95 density in the pore lumen: reducing or increasing the FG-spacing leads Kap95 to localize near the densest regions or spread out more evenly throughout the pore region, respectively. Interestingly, increasing the FG-spacing to large values (d_FG_=104) enhances the portion of the pore volume that is sampled by Kap95.

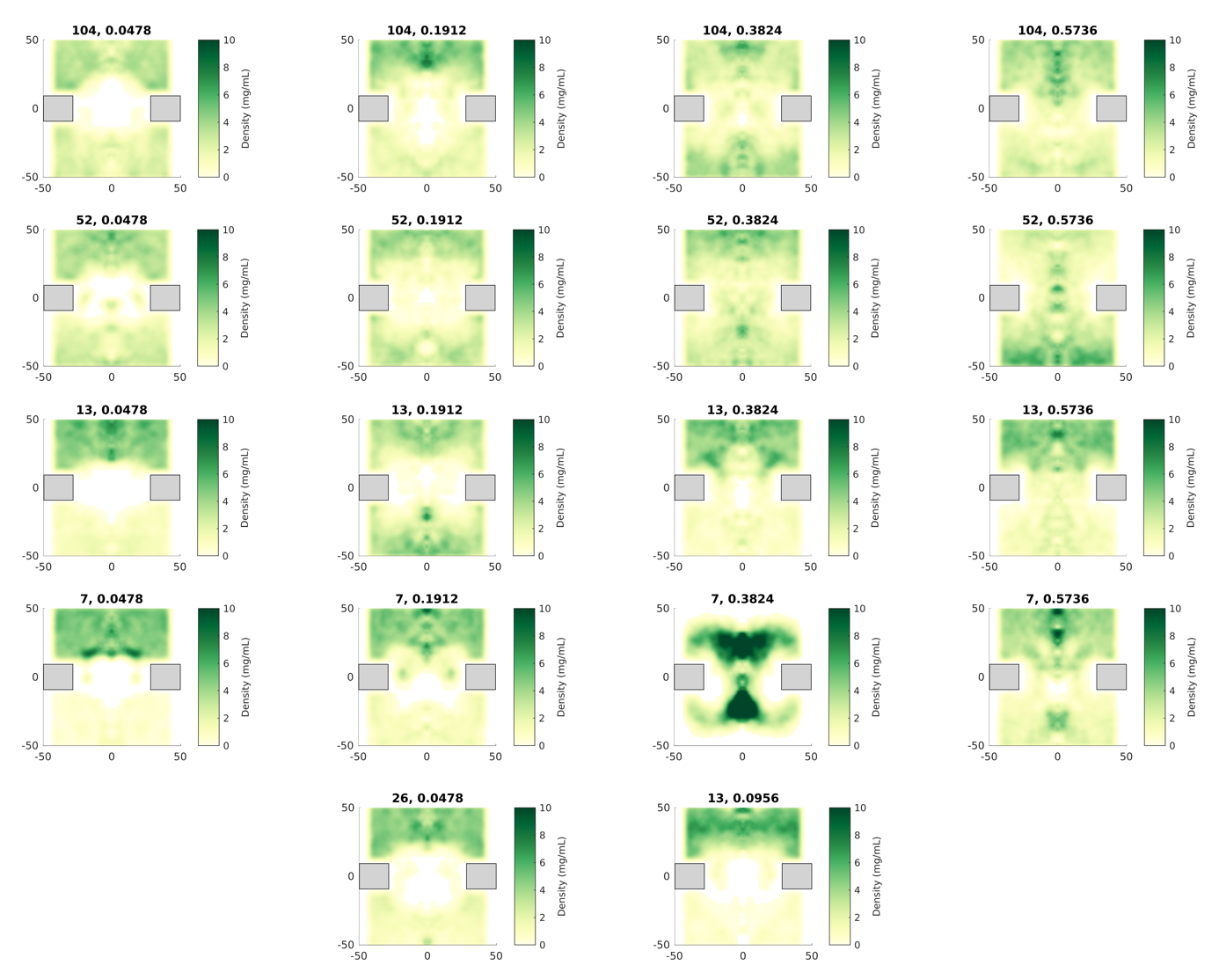

Supplementary Figure 17 – Time-averaged, axi-radial density graphs of the BSA protein in nanopores coated with artificial FG-Nups in presence of Kap95 and inert proteins, grouped by C/H and FG-spacing. Scaffolds are highlighted in gray. Additional variants with 2xFG-spacing and 2xC/H-ratio are shown separately on the bottom row. C/H=0.0239 (0.5x native average) variants were not studied in presence of cargo and NTRs. The density distributions for BSA (initially inserted from the top side of the nanopore, and subsequently expelled to the top side) indicate that changes in C/H (horizontal direction) generally lead to a reduction of barrier function. Any BSA permeation events for lower C/H-values (0.0478, 0.0956, 0.1912) take place via the sparser regions near the pore wall, formed by the extended anchoring domain of our artificial FG-Nups. When C/H is increased further, translocation events start to take place via the central channel of the nanopore as well, as the ‘dense plug’ formed by artificial FG-Nups for low C/H is dissolved. For variations in FG-spacing, only a decrease (d_FG_=7) seems to affect the distribution of BSA, due to the increased FG-Nup density in such cases.

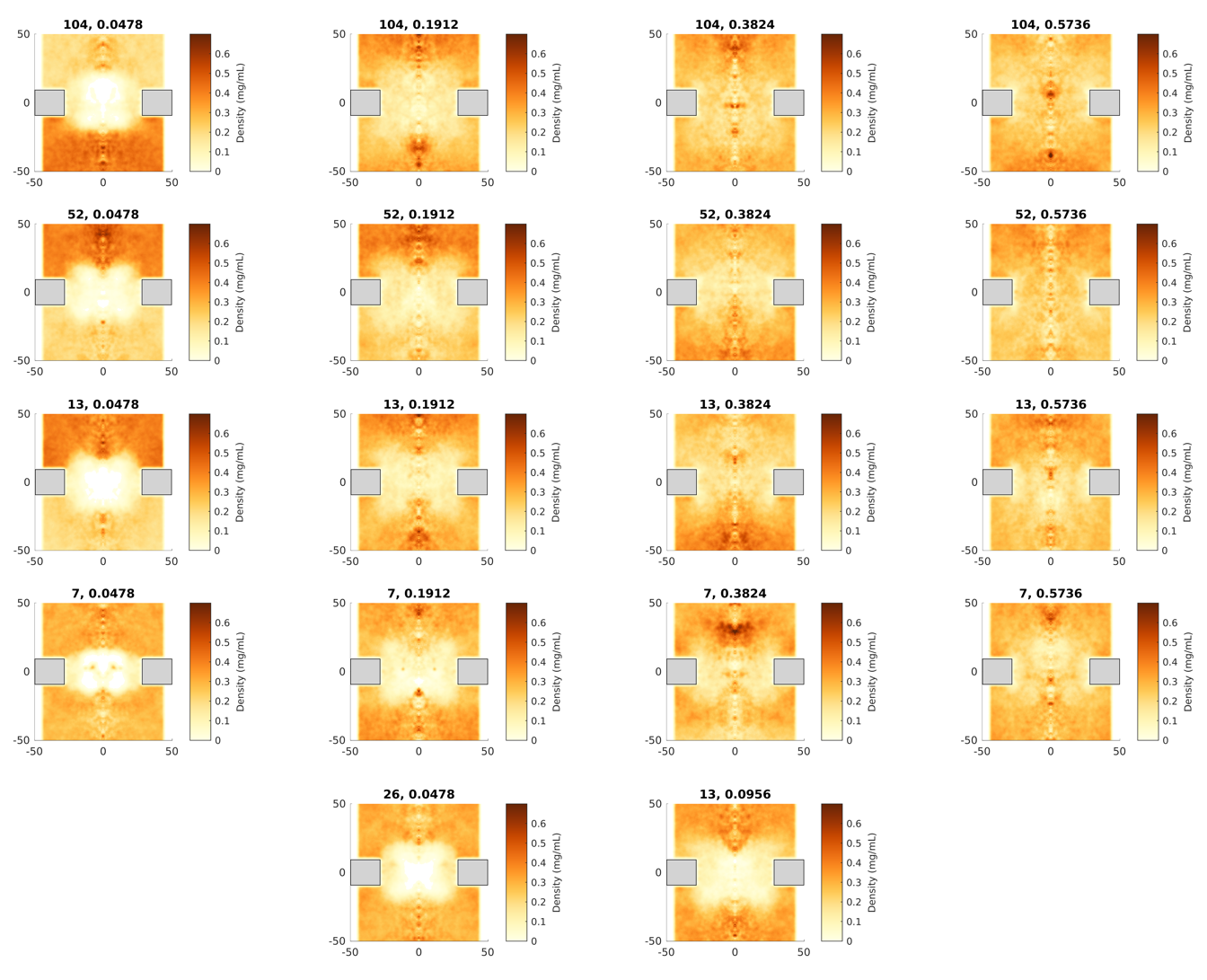

Supplementary Figure 18 – Time-averaged, axi-radial density graphs of the Ubiquitin protein in nanopores coated with artificial FG-Nups in presence of Kap95 and inert proteins, grouped by C/H and FG-spacing. Scaffolds are highlighted in gray. Additional variants with 2xFG-spacing and 2xC/H-ratio are shown separately on the bottom row. C/H=0.0239 (0.5x native average) variants were not studied in presence of cargo and NTRs. Similar to the results for BSA (Supplementary Figure 9), the permeability of FG-Nup-coated nanopores for small inert proteins increases with C/H (horizontal direction). Localization of Ubiquitin in FG-Nup coated pores takes place near the sparse, extended anchoring domains close to the pore wall for low C/H (0.0478, 0.0956). As the value of C/H increases beyond 0.0956, Ubiquitin localizes rather evenly throughout the nanopore system. The effect of FG-spacing on the distribution of Ubiquitin is modest, with only a reduction (d_FG_=7) w.r.t. the native GLFG-Nup average leading to a noticeable change in the average Ubiquitin distribution.

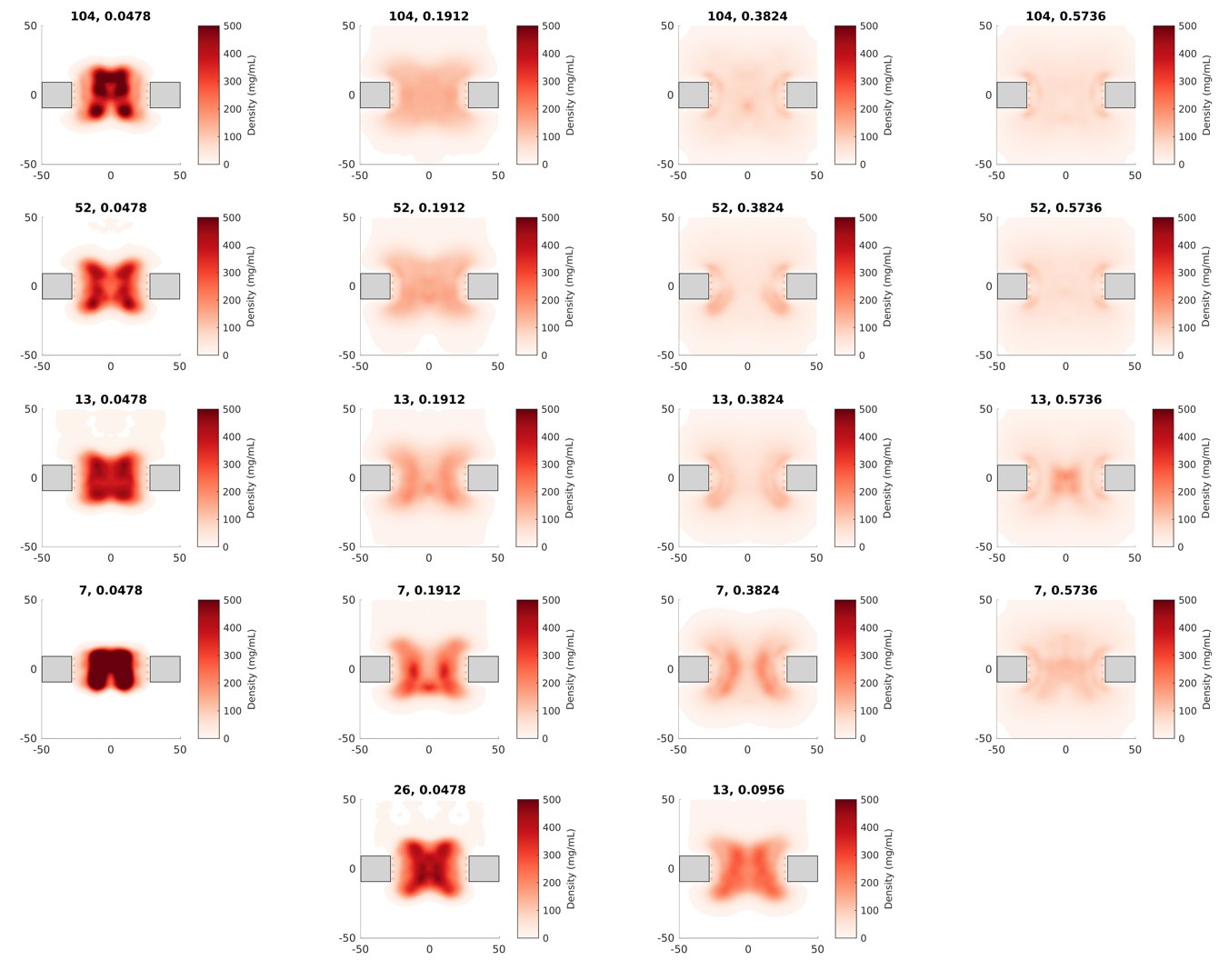

Supplementary Figure 19 – Time-averaged, axi-radial density graphs of the entire protein mass (FG-Nups and inert proteins and Kap95) in nanopores coated with artificial FG-Nups in presence of Kap95 and inert proteins, grouped by C/H-ratio and FG-spacing.

Scaffolds are highlighted in gray. Additional variants with 2xFG-spacing and 2xC/H-ratio are shown separately on the bottom row. C/H=0.0239 (0.5x native average) variants were not studied in presence of cargo and NTRs.

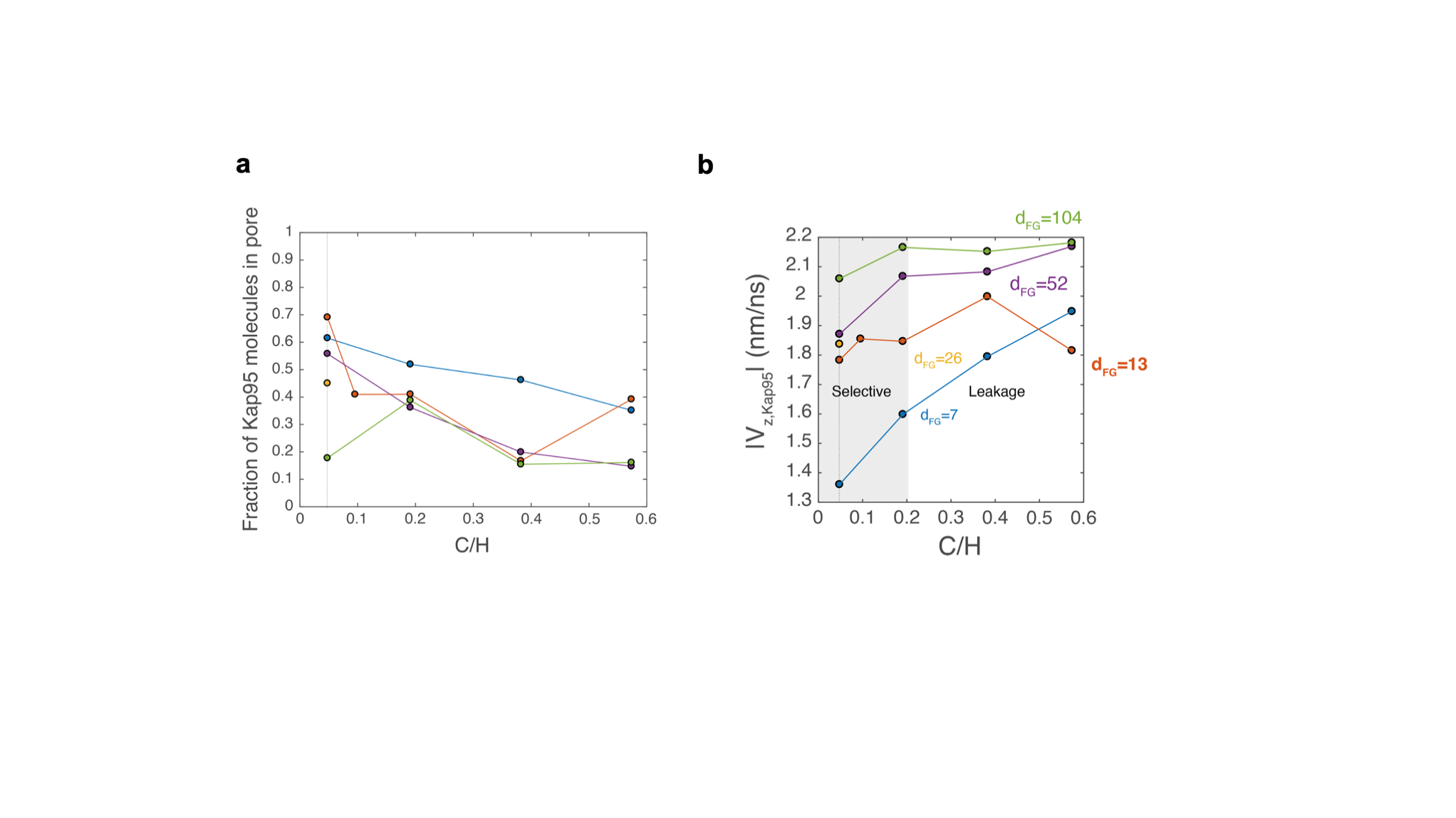

Supplementary Figure 20 – Occupancy and on-axis velocity magnitude of Kap95.
**a:** Time-averaged fraction of Kap95 molecules that reside within the pore. With increasing C/H, the occupancy decreases. **b**: Velocity magnitude, averaged over the pore interior and over time, for Kap95 molecules in NupY-variant-coated pores. Mobility of Kaps increases with C/H (reduced steric hindrance) and with dF_G_ (less frequent binding).

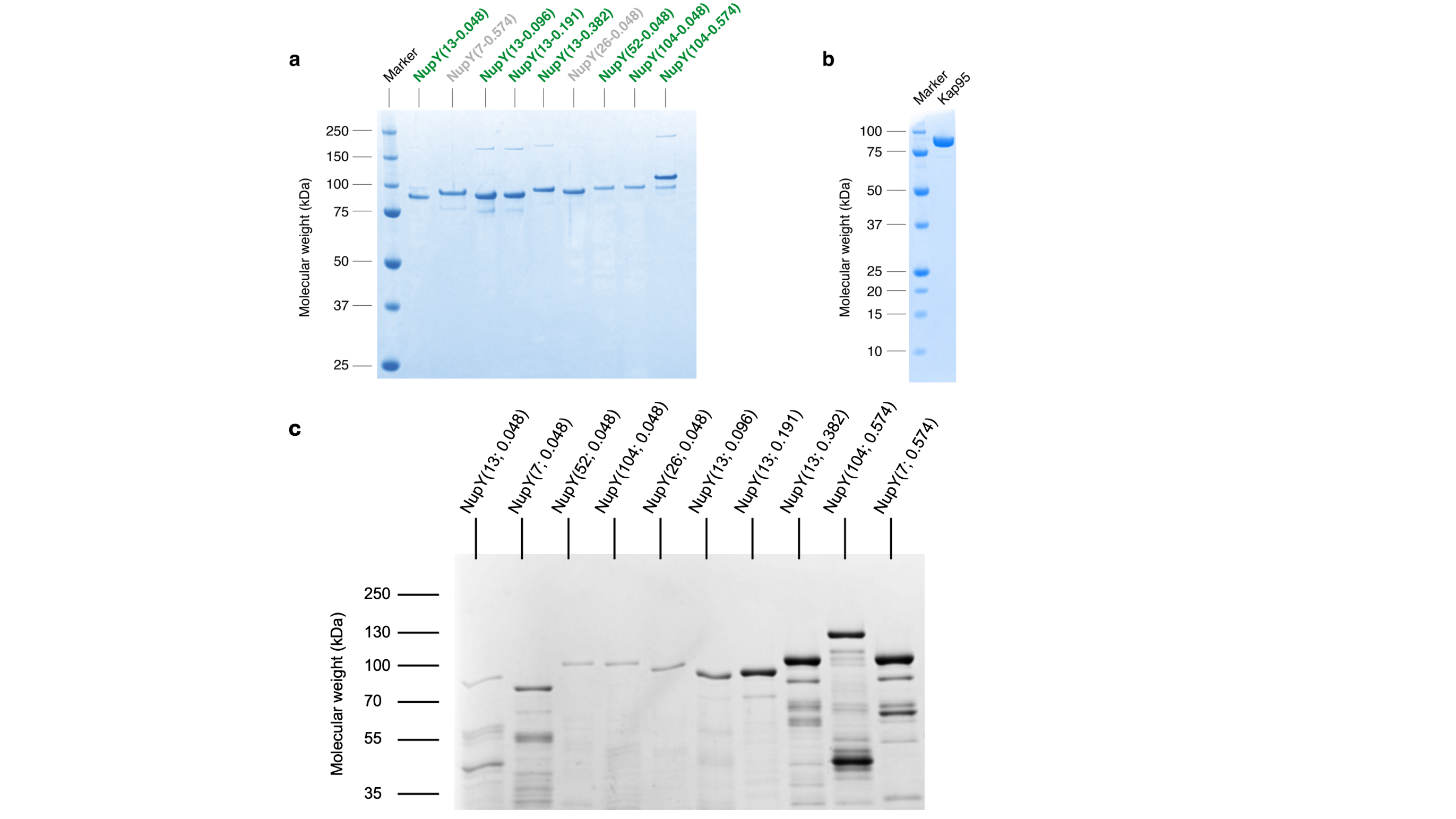

### Supplementary Figure 21– SDS-PAGE gel of purified NupY variants and Kap95.

**a)** Coomassie-stained SDS-PAGE gel showing purified NupY variants running between ~80–120 kDa. Green labels indicate the NupY variants used for QCMD-D and LLPS experiments, whereas the grey ones (NupY(7;0.574) and NupY(26;0.048)) could not be used due to unsatisfactory expression levels. **b)** Coomassie-stained SDS-PAGE for Kap95 confirms the high purity of the Kap95 sample. **c)** Coomassie Brilliant Blue G250 SDS-PAGE gel of purified NupY variants used in Figure S7D.
